# A longitudinal single-cell and spatial multiomic atlas of pediatric high-grade glioma

**DOI:** 10.1101/2024.03.06.583588

**Authors:** Jonathan H. Sussman, Derek A. Oldridge, Wenbao Yu, Chia-Hui Chen, Abigail M. Zellmer, Jiazhen Rong, Arianne Parvaresh-Rizi, Anusha Thadi, Jason Xu, Shovik Bandyopadhyay, Yusha Sun, David Wu, C. Emerson Hunter, Stephanie Brosius, Kyung Jin Ahn, Amy E. Baxter, Mateusz P. Koptyra, Rami S. Vanguri, Stephanie McGrory, Adam C. Resnick, Phillip B. Storm, Nduka M. Amankulor, Mariarita Santi, Angela N. Viaene, Nancy Zhang, Thomas De Raedt, Kristina Cole, Kai Tan

## Abstract

Pediatric high-grade glioma (pHGG) is an incurable central nervous system malignancy that is a leading cause of pediatric cancer death. While pHGG shares many similarities to adult glioma, it is increasingly recognized as a molecularly distinct, yet highly heterogeneous disease. In this study, we longitudinally profiled a molecularly diverse cohort of 16 pHGG patients before and after standard therapy through single-nucleus RNA and ATAC sequencing, whole-genome sequencing, and CODEX spatial proteomics to capture the evolution of the tumor microenvironment during progression following treatment. We found that the canonical neoplastic cell phenotypes of adult glioblastoma are insufficient to capture the range of tumor cell states in a pediatric cohort and observed differential tumor-myeloid interactions between malignant cell states. We identified key transcriptional regulators of pHGG cell states and did not observe the marked proneural to mesenchymal shift characteristic of adult glioblastoma. We showed that essential neuromodulators and the interferon response are upregulated post-therapy along with an increase in non-neoplastic oligodendrocytes. Through *in vitro* pharmacological perturbation, we demonstrated novel malignant cell-intrinsic targets. This multiomic atlas of longitudinal pHGG captures the key features of therapy response that support distinction from its adult counterpart and suggests therapeutic strategies which are targeted to pediatric gliomas.

## Main

Pediatric high-grade glioma (pHGG) is a devastating brain malignancy accounting for approximately 11% of central nervous system (CNS) tumors in children from infants to adolescents^1^. Although the incidence of this tumor is relatively low (1.78 per 100,000 population)^2^, pHGG holds an exceptionally dismal prognosis, with a median overall survival of 14 to 20 months.^3^ Despite decades of research and over 1,500 clinical trials, there remains no cure for pHGG. Standard therapy includes maximal safe resection, high-dose radiotherapy, and chemotherapy^4^, yet this multimodal therapy does little to change the course of the disease^5^. Although childhood and adult HGG, including glioblastoma multiforme (GBM), share many histopathological and clinical features, the advent of genomic, transcriptomic, and epigenomic profiling has led pHGG to be recognized as a distinct disease entity with substantial differences in its molecular characteristics^2,6–10^. Most prominently, mutations in the histone H3 gene (*H3F3A* and *HIST1H3B*) define important anatomically-distinct subtypes of pediatric gliomas^6^. The H3K27M mutation occurs frequently in tumors arising in the brainstem and other midline structures including the thalamus and cerebellum, while the H3G34R/V mutation is found most frequently in adolescent pHGGs of the cerebral cortex^2,7^. Other mutations in genes such as *BRAF* and *ACVR1* are found predominantly in pediatric, rather than adult gliomas, yet their implications for diagnosis and treatment have not been established^7,11^. However, despite advances in delineating genomic subtypes, pHGG remains extremely heterogeneous with a desperate need for improved therapeutic options.

Recent advances in single-cell multiomics and spatial profiling have greatly informed our understanding of the intra-tumoral and inter-tumoral heterogeneity of adult and pediatric brain tumors^12–23^. Collectively, these studies have identified patterns of neoplastic cell differentiation states and metabolic programs, proposed detailed models for tumor initiation and oncogenesis, characterized the tumor immune microenvironment (TIME), and identified actionable avenues for targeted chemotherapeutic and immunotherapeutic strategies. Importantly, recent studies using bulk and single-cell transcriptomics have identified key cellular and microenvironmental changes during adult glioma progression under standard therapy, such as a shift in neoplastic cell states from a proneural to mesenchymal phenotype^23,24^, which has been implicated in glioma treatment resistance^25^. However, current single-cell characterization of pHGG is largely limited to the neoplastic cell compartment^17,12,16,20,22^, and the extent to which pHGG progression under therapy differs from that of adult HGG is unknown. To address this, we present an integrated multimodal analysis of matched primary-recurrent patient specimens (16 patients) across histologic and molecular subtypes using single-nucleus RNA-sequencing (snRNA-Seq), single-nucleus assay for transposase-accessible chromatin via sequencing (snATAC-Seq), whole genome sequencing (WGS), and Co-Detection by Indexing (CODEX) spatial proteomics. Overall, this longitudinal multiomic atlas of pHGG captures key features of therapy response that support its distinction from adult HGG and suggests therapeutic strategies which are targeted to pediatric gliomas.

## Results

### Single-cell profiling of longitudinal pHGG specimens

We profiled pHGG samples obtained through the Children’s Brain Tumor Network (CBTN)^26^ from 16 patients across therapeutic time points via snRNA-Seq (15 pairs) and snATAC-Seq (11 pairs) (**Figure 1a-c**). All patients received radiotherapy and surgical resection, and some received pharmacological treatment including temozolomide, immunotherapy (e.g., pembrolizumab), and cytotoxic chemotherapy (**Extended Data Fig. 1a, Supplementary Fig. 1, Supplementary Table 1)**. Patients in the cohort ranged from 4 to 24 years in age, had a male-to-female ratio of 2.2, and tumors included a range of genomic alterations (**Extended Data Fig. 1b**). The tumor specimens were resected from multiple anatomic locations including cortical lobes and midline structures including the thalamus and cerebellum. The cohort included three H3K27M-mutated cases, one H3G34V-mutated case, and one IDH1-mutated case; the remainder were IDH1/H3 wildtype (WT) (**Figure 1d)**. Post-therapy time points were further delineated as progressive/recurrence, where samples were obtained through a secondary resection, and autopsy, where samples were collected post-mortem. Collectively, over 400,000 cells were profiled via snRNA-Seq, and over 110,000 cells were profiled via snATAC-Seq after quality assessment and filtering, capturing a mean of 2,280 genes and 19,094 unique chromatin fragments per cell respectively (**Extended Data Fig. 2a**, **Extended Data Fig. 3a**). Samples were integrated to remove batch effects and cell types were annotated (**Extended Data Fig. 2b-e**, **Extended Data Fig. 3b-e, Supplementary Fig. 1, 2, Methods**). We captured the major cell types present in gliomas, including normal mature neurons and oligodendrocytes, myeloid cells (macrophages/microglia), T cells, endothelial cells, mural cells, and a diverse population that we have termed “other neural and glial cells,” including a mix of inferred neoplastic and non-neoplastic subpopulations. (**Figure 1b-c)**. There was significant heterogeneity between patients and time points (**Figure 1d, Extended Data Figure 2f-g**, **Extended Data Figure 3f-h**). Notably, the majority of mural cells were captured within two patients, and T cells were captured largely in a single patient (**Figure 1d)**. Examining the longitudinal shifts in cell type composition in the snRNA-Seq data revealed a significant increase in non-neoplastic oligodendrocytes (p*=*0.0067) and mature neurons (p*=*0.029) within patient-matched pairs (**Figure 1e**). Oligodendrocytes were concordantly enriched post-therapy in the snATAC-Seq data (p*=*0.019) (**Figure 1f**), consistent with prior observations of oligodendrocyte expansion in adult glioblastoma multiforme (GBM)^23^. This trend occurred primarily in the secondary resection samples, suggesting this is not simply an artifact of wider normal margins in autopsy specimens **(Extended Data Fig. 2f**, **Extended Data Fig. 3f)**.

**Figure 1.**
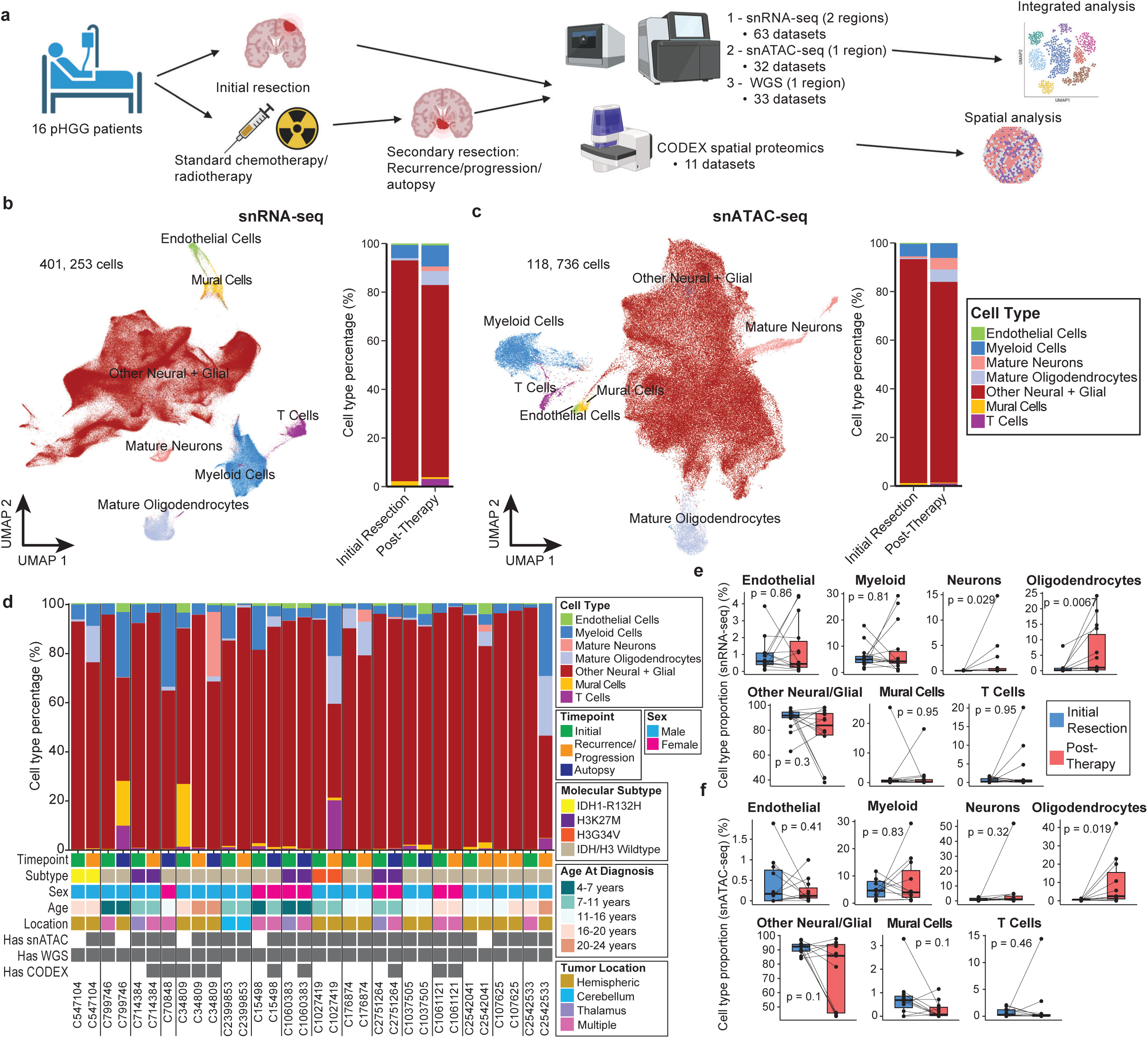
Longitudinal single-cell RNA and ATAC atlas of pediatric high grade glioma (pHGG) **a)** Overview of the multiomics studies on patient-matched longitudinal pHGG specimens. **b-c)** Uniform manifold approximation and projection (UMAP) of (**b**) snRNA-Seq data (401,253 cells) and (**c**) snATAC-Seq data (118,736 cells) annotated by major cell type category (left) and stacked bar plot of cell type proportions across dataset comparing initially resected pHGG samples with post-therapy samples. **d)** Cell type proportions in snRNA-Seq data across each patient and therapeutic time point, along with a summary of patient demographics and molecular subtype. **e-f)** Shifts in cell type proportions for each patient between initial resection and post-therapy time points in (**e**) snRNA-Seq and (**f**) snATAC-Seq; *n* = 15 paired samples profiled by snRNA-Seq and *n* = 11 paired samples profiled by snATAC-Seq, including an initial resection and at least one post-therapy sample. Post-therapy samples were merged for one patient with three longitudinal samples. A two-sided Wilcoxon signed-rank test for paired samples was used.

**Figure 2.**
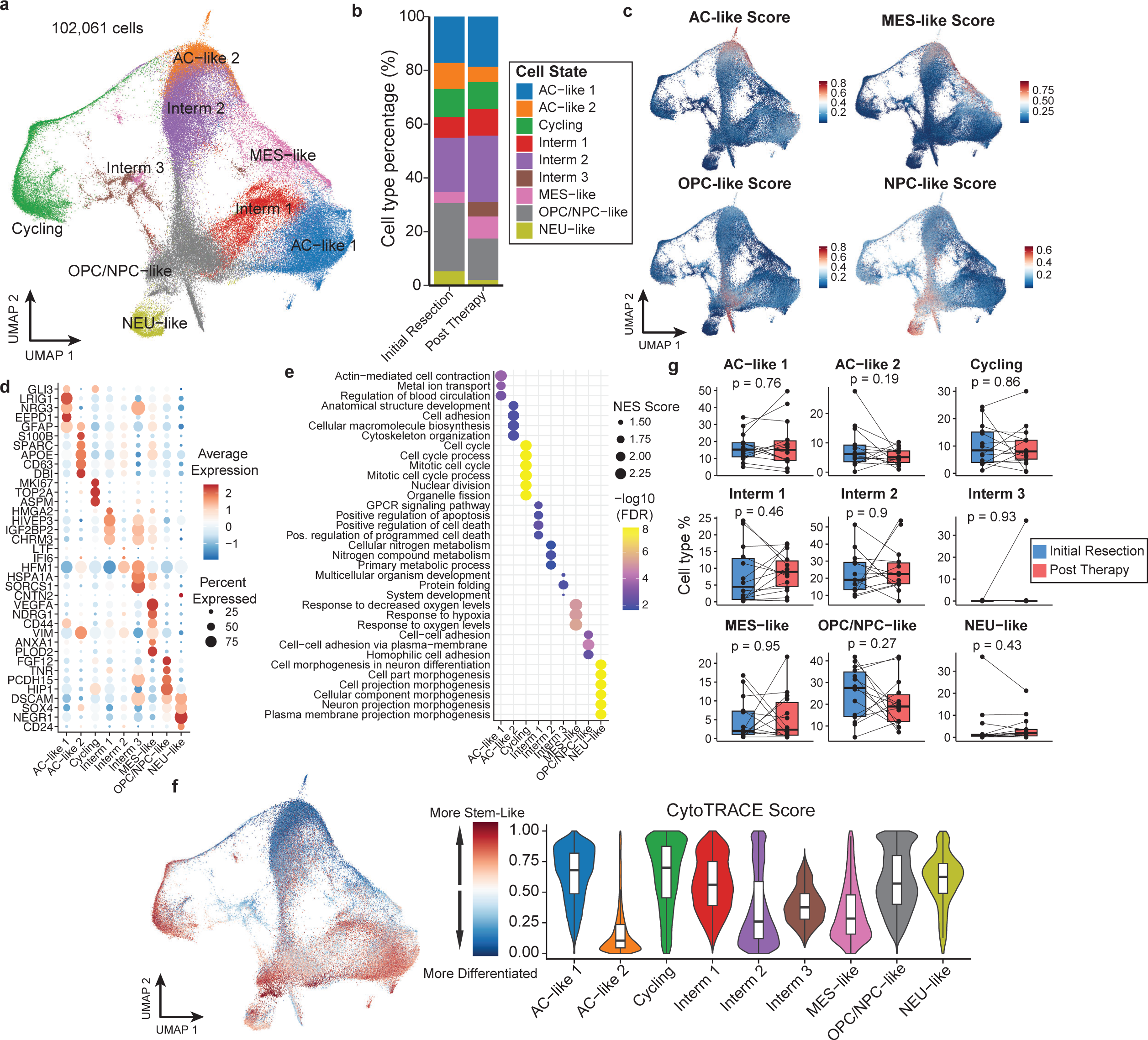
Transcriptional states of pHGG neoplastic cells. **a)** UMAP projection of inferred neoplastic cells from snRNA-Seq (102,061 cells) after integration and annotation of cell states; AC, astrocyte; MES, mesenchymal; OPC, oligodendrocyte progenitor cell; NPC, neural progenitor cell; NEU, neural. **b)** Barplot of cell type proportions of neoplastic cells across dataset comparing initial resection and post-therapy samples. **c)** Gene signatures of GBM cell states^12^ overlaid on UMAP of neoplastic cells. Colors truncated at 1^st^ and 99^th^ percentiles for visualization. **d)** Expression of representative differentially expressed genes across neoplastic cell states in snRNA-Seq data. **e)** Gene set enrichment of top differentially expressed genes in each neoplastic cell state using biological process terms from the Gene Ontology database. **f)** CytoTRACE scores of inferred differentiation states on the UMAP projection of snRNA-Seq data (left) and across each cell state (right). Higher values indicate a more undifferentiated/stem-like state and lower values indicate a more differentiated state. **g)** Shifts in neoplastic cell state proportions for each patient between initial resection and post-therapy time points in snRNA-Seq (*n* = 15 paired samples). Post-therapy samples were merged for one patient with three longitudinal samples. A two-sided Wilcoxon signed-rank test for paired samples was used.

**Figure 3.**
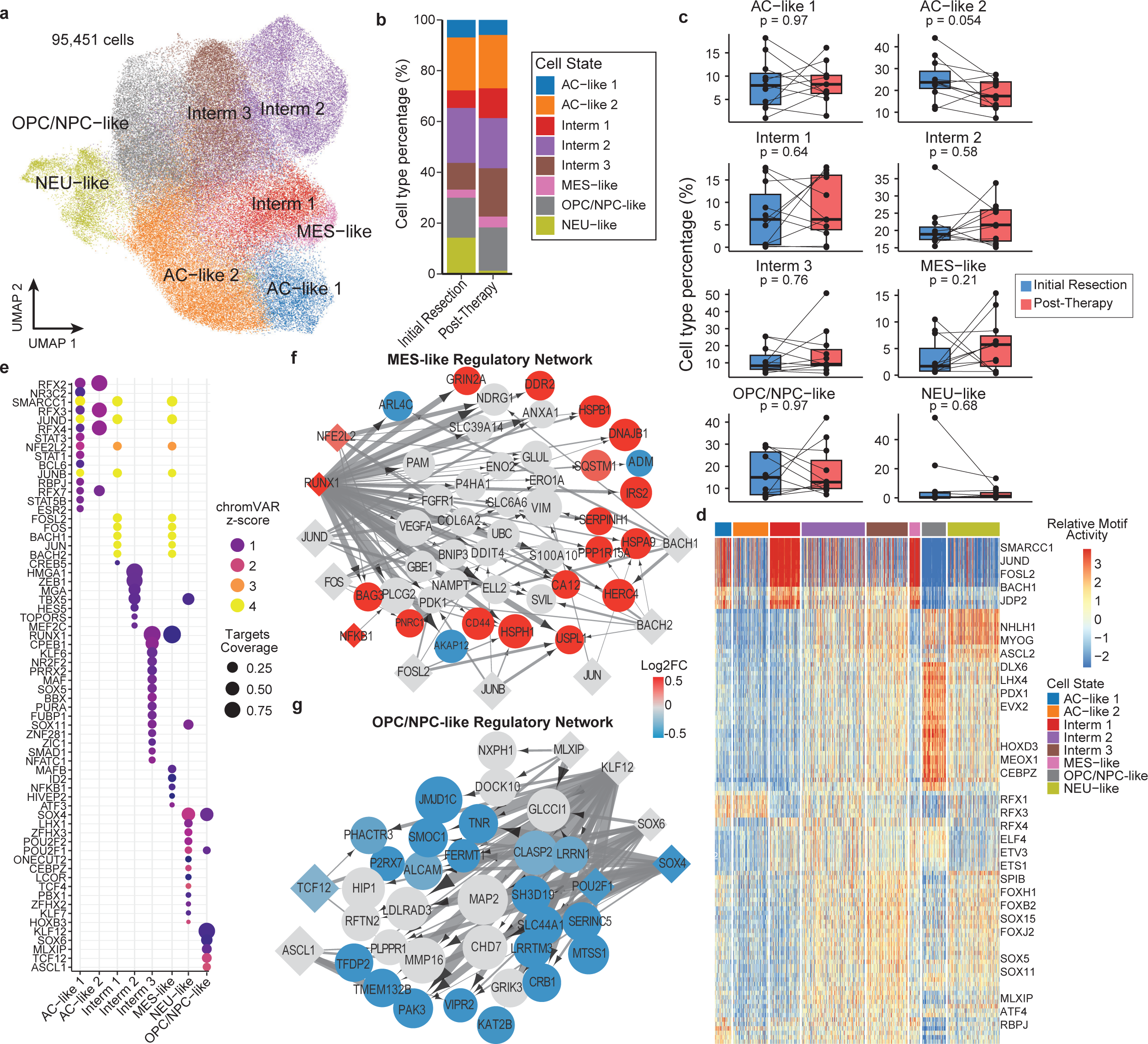
Transcriptional regulation of pHGG neoplastic cell states. **a)** UMAP projection of inferred neoplastic cells from snATAC-Seq (95,451 cells) after integration and identification of cell states defined in the snRNA-Seq data; AC, astrocyte; MES, mesenchymal; OPC, oligodendrocyte progenitor cell; NPC, neural progenitor cell; NEU, neural. **b)** Stacked barplot of cell type proportions of neoplastic cell states across dataset comparing initial resection and post-therapy samples. **c)** Shifts in neoplastic cell state proportions for each patient between initial resection and post-therapy time points in snATAC-Seq (*n* = 11 paired samples). Post-therapy samples were merged for one patient with three longitudinal samples. A two-sided Wilcoxon signed-rank test for paired samples was used. **d)** Heatmap of differential transcription factor (TF) motif accessibility in each pHGG neoplastic cell state. Values are z-score-normalized deviation scores calculated using chromVAR. The differential TF accessibility analysis was performed by a Wilcoxon rank-sum test, comparing chromVAR deviation score between each cell state and the other cell states. The top 20 differential TFs are displayed for each state. **e)** Overview of top 15 significant transcriptional regulators for each neoplastic cell state based on predicted enhancer-promoter interactions and TF-target gene pairs. The size of the dot indicates the fraction of the total gene targets in the network regulated by each TF. Color indicates chromVAR deviation z-score as in (**d**). **f-g**) Transcriptional regulatory networks (TRNs) for (**f**) MES-like state and (**g**) OPC/NPC-like state, showing top 50 upregulated genes and top 15 TFs in each TRN. Diamond nodes represent TFs and circle nodes represent target genes. Node size is proportional to the average gene expression for target genes and average chromVAR z-score for TFs. Node color is proportional to the average log_2_ fold change of the gene in that cell state post-therapy across all cells. Edge line thickness is proportional to the linear regression coefficient for the predicted enhancer-promoter interaction and the fraction of cells with chromatin accessibility at the enhancer peak.

### Pediatric gliomas exhibit distinct neoplastic cell states

We sought to characterize the neoplastic cell compartment and assess how these cell states change during progression and therapy. After identifying putative neoplastic cells via copy number variation (CNV) inference (**Supplementary Figure 3a-b, Methods**), we reintegrated these populations (**Extended Data Figure 4a-d)** and then examined whether the canonical cell states established by Neftel *et al*.^12^ in a cohort of IDH-wild-type adult and pediatric glioblastoma (GBM) can be applied to a molecularly diverse cohort of pHGG using the snRNA-Seq data. Assessing the gene signatures of these four states (astrocyte (AC)-like, mesenchymal (MES)-like, oligodendrocyte-progenitor (OPC)-like, neural-progenitor (NPC)-like) yielded several key observations. First, we identified two distinct AC-like populations (**Figure 2a-d**). These populations both expressed the astrocyte-defining marker, *GFAP,* and were most enriched in the AC-like gene signature (**Figure 2c**, **Extended Data Figure 4e**). Next, we identified a definitive MES-like state which expressed established mesenchymal marker genes (e.g., *CD44, VIM, ANXA1, NDRG1*) and angiogenesis genes (i.e., *VEGFA*) and was enriched in hypoxia response signatures (**Figure 2d-e, Supplementary Table 2**). Interestingly, while the mesenchymal cell state has only recently been identified in H3K27M-mutant glioma^20^, which these findings further support, we observe a low-frequency MES-like state in our single pediatric IDH-mutant glioma case (**Extended Data Fig. 4i**).

**Figure 4.**
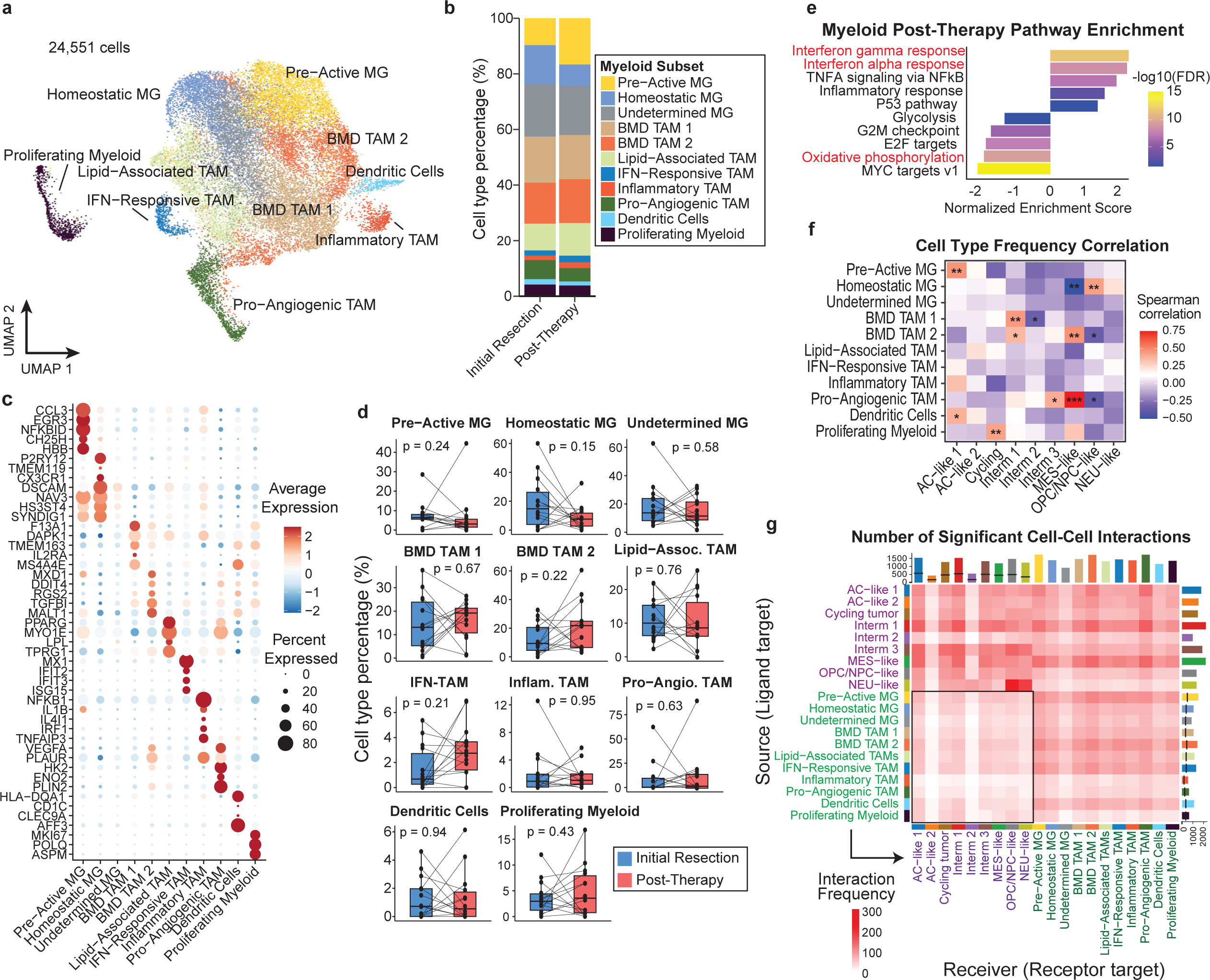
The myeloid response to progression and therapy. **a)** UMAP projection of annotated tumor-associated myeloid cell populations identified in integrated longitudinal pHGG snRNA-Seq atlas (24,551 cells). BMD, bone marrow-derived; MG, microglia. **b)** Stacked barplot of myeloid cell type composition across dataset comparing initial resection and post-therapy samples. **c)** Expression of representative genes across myeloid populations in snRNA-Seq data highlighting top differentially expressed genes. **d)** Shifts in myeloid cell type proportions for each patient between initial resection and post-therapy time points in snRNA-Seq (*n* = 15 paired samples). Post-therapy samples were merged for one patient with three longitudinal samples. A two-sided Wilcoxon signed-rank test for paired samples was used. **e)** Gene set enrichment analysis (GSEA) of Hallmark pathways comparing pathway-level differences in gene expression within myeloid cells overall between initial resection and post-therapy time points. A linear mixed model was used to identify differentially expressed genes between time points while accounting for individual patient variability. **f)** Heatmap of Spearman correlation coefficients between frequency of neoplastic cell states in the malignant population and frequency of TAM subtypes in the myeloid population across region-stratified samples in the snRNA-Seq data (n = 63). P-values are adjusted using the Benjamini-Hochberg method; *** p <0.001, ** p <0.01, * p <0.05. **g)** Heatmap showing number of significant interactions between myeloid and neoplastic cell populations across dataset. Interactions were inferred using LIANA^72^ and filtered for aggregated consensus rank (adjusted p-value < 0.05). Bars above the heatmap represent total number of significant interactions received (down columns) and bars to the right of the heatmap represent total number of significant interactions sent (across rows) for each cell type, with black lines indicating subset of interactions between myeloid and neoplastic cells. Box highlights interactions from myeloid cells to neoplastic cells.

We identified a population with high expression of both OPC-like and NPC-like gene signatures that we refer to as OPC/NPC-like, and a distinct population expressing neural genes (**Figure 2c-e**) that was restricted to IDH/H3-WT tumors (**Extended Data Fig. 4i**). Projecting the cells onto an atlas of the developing human fetal brain revealed that this population most closely resembled fetal excitatory neurons, rather than earlier neural progenitor phenotypes (**Extended Data Fig. 4f**), and pathway analysis supported the expression of neuronal pathways (**Figure 2e)**, thus this population was annotated as neuronal (NEU)-like. OPC/NPC-like cells expressed both known OPC-like genes (e.g., *FGF12*) and NPC-like genes (e.g., *TNR*), and NEU-like cells expressed some NPC-like marker genes (e.g., *SOX4, CD24*) (**Figure 2d**). Lastly, we identified three distinct intermediate cell states that lack specific enrichment of the canonical markers and identified a mixed population of cycling cells (**Extended Data Fig. 4e**). An analysis of neoplastic lineages via CytoTRACE^27^, which leverages transcriptional diversity to predict differentiation trajectories, supported the proneural to mesenchymal differentiation hierarchy^14,23^, and suggested that the two AC-like states lie on either ends of the differentiation spectrum (**Figure 2f**). The AC-like 1 population is the least differentiated neoplastic cell state while the AC-like 2 population is the most differentiated cell state and was found to also express some mesenchymal markers (e.g., *VIM, APOE*) in addition to canonical AC-like markers (e.g., *S100B, SPARC*) (**Figure 2d**).

The proportions of these cell states across samples were highly heterogeneous, with significant variation between patients and across time points (**Extended Data Fig. 4g-i**). However, examining neoplastic cell state proportions per-patient revealed no significant shifts in cell type composition across therapeutic time points (**Figure 2g**). This finding is in contrast with recent findings in adult IDH-WT GBM in which a significant increase in mesenchymal cells was observed after treatment and progression.^23^

### Transcription factors jointly regulate pHGG neoplastic cell states

We then sought to extend our characterization of these cell states via our single-cell chromatin accessibility data. After reintegrating and annotating the putative neoplastic population in our snATAC-Seq data (**Extended Data Fig. 5a-c, Supplementary Fig. 4a-d, Methods**), we captured all the cell states defined transcriptionally in the snRNA-Seq data and confirmed enrichment of chromatin accessibility for cell state-defining genes and significant concordance with snRNA-Seq (**Figure 3a-b, Supplementary Fig 4e-f**). Of note, we did not identify a distinct population of cycling cells in the snATAC-Seq data (**Supplementary Fig. 4c**). As expected, we observed significant heterogeneity between patients and therapeutic time points, and no significant shifts in neoplastic cell state post-therapy (**Extended Data Fig. 4d-f, Figure 3c**). However, we observed a decreasing trend in the AC-like 2 population in the majority of patient-matched pairs with borderline significance (p=0.054) (**Figure 3c)**.

**Figure 5.**
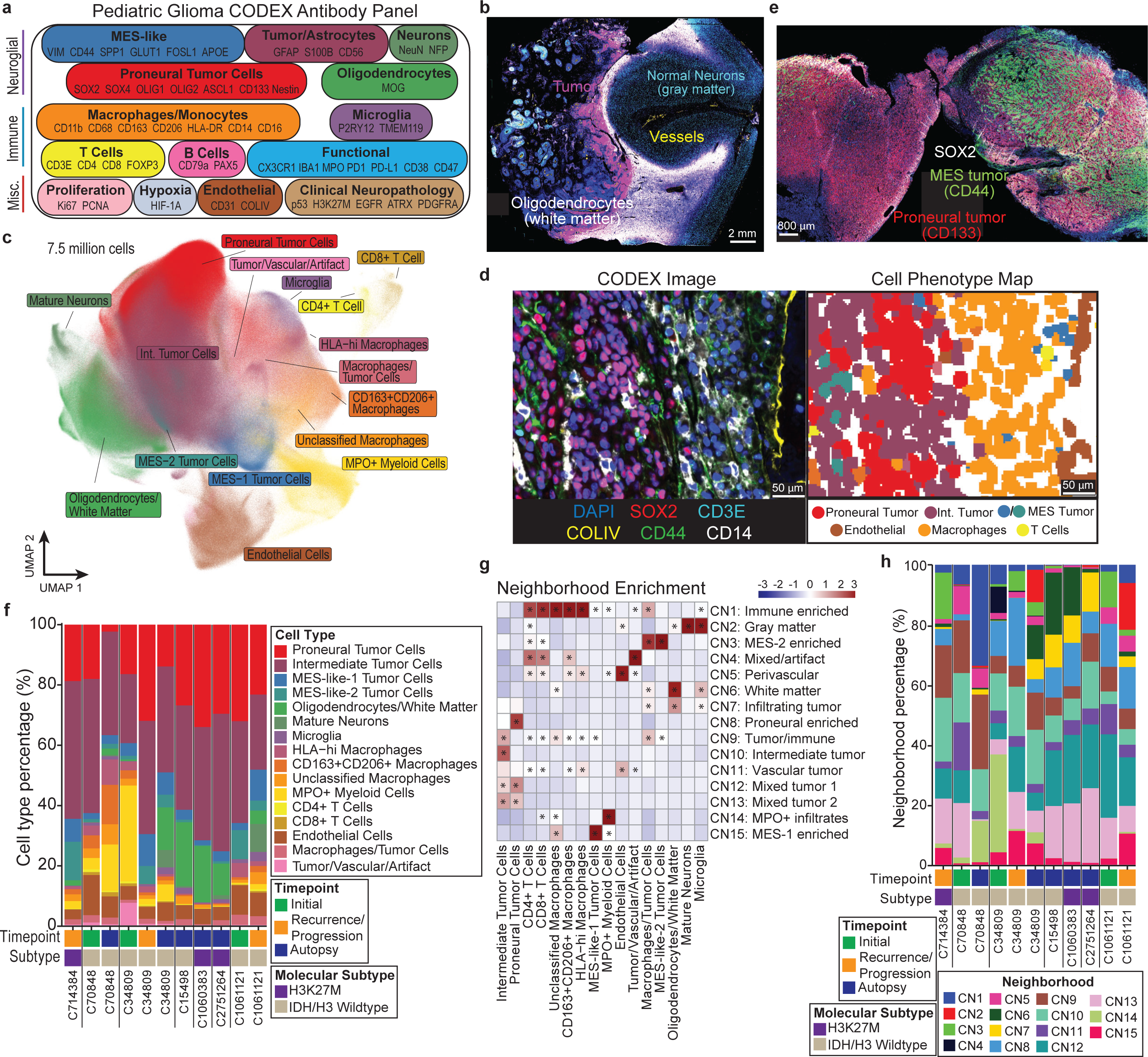
CODEX imaging reveals the spatial landscape of pHGG. **a)** Diagram showing the 51-antibody CODEX panel split by target cell population or cellular function. **b)** Representative CODEX image highlighting tumor mass and substructures of the normal brain. DAPI (blue), Collagen IV (yellow), Neu (cyan), SOX2 (magenta), MOG (white). **c)** UMAP projection of all 7.5 million cells in the pHGG CODEX dataset across 11 samples after annotation and filtering. **d)** CODEX image with selected fluorescent markers (left) paired with cell phenotype map (right). Segmentation masks of individual cells are colored by their identity. **e)** Representative CODEX image demonstrating spatially restricted tumor cell state populations. Proneural tumor cells are stained by CD133 (red) and SOX2 (white) and mesenchymal tumor cells stained by CD44 (green). **f)** Cell type proportions in each CODEX sample, indicating patient, therapeutic time point, and molecular subtype. **g)** Heatmap showing relative enrichment of the cell types present in neighborhoods, normalized across neighborhoods (by column). Significant positive cell type enrichments in each neighborhood were calculated using a hypergeometric test, adjusted using the Benjamini-Hochberg method. * p-adjusted <0.001. **h)** Neighborhood proportions in each CODEX sample, indicating patient, therapeutic time point, and molecular subtype.

We then aimed to identify the transcription factors that regulate each cell state, first by using chromVAR^28^ to assess differential accessibility of transcription factor motifs (**Supplementary Table 3**). Consistent with previous reports^23,29^, motifs for the AP1 family of transcription factors (e.g., FOSL2, JUN) were enriched in the mesenchymal state, along with SMARCC1, JDP2, and BACH1 (**Figure 3d**). Notably, these motifs were also enriched in the Intermediate 1 and AC-like 1 states. Both AC-like states were enriched for RFX factor motifs, and the OPC/NPC-like and NEU-like states were enriched for proneural transcription factors (e.g., ASCL2, NHLH1, LHX4) (**Figure 3d**).

Next, we constructed a transcriptional regulatory network (TRN) for each cell state by integrating our snRNA-Seq and snATAC-Seq data to predict state-specific enhancer-promoter interactions and transcription factor-target gene pairs (**Supplementary Table 4, Methods**). These TRNs revealed substantial cooperativity between transcription factors in regulating cell state-specific gene expression (**Figure 3e-g, Extended Data** Fig. 4g-j). This analysis nominated known and novel transcription factors. The RFX factors were predicted to regulate both AC-like cell states^30^. SOX4 was predicted to regulate both the NEU-like and OPC/NPC-like states through cooperation with other transcription factors including LHX1 and KLF12 respectively (**Figure 3e-f**). The AP1 factors^23^ and RUNX1 were predicted to jointly regulate the MES-like state. RUNX1 was predicted to target the top differentially expressed genes in the MES-like state and *RUNX1* expression was upregulated in MES-like neoplastic cells post-therapy (**Figure 3f**). While RUNX1 has been recognized as a contributor to mesenchymal GBM^31^, this analysis suggests that the RUNX1 transcription factor is a central regulator of the MES-like state. Interestingly, we observed that 39% of genes in the MES-like TRN were significantly upregulated within that population post-therapy (versus 9.3% downregulated), while 49% of genes within the OPC/NPC-like TRN were significantly downregulated (versus 1.4% upregulated). This suggests that although there is no population shift, the MES-like state phenotype may be strengthened post-therapy. Overall, this analysis revealed the overlapping transcriptional regulatory interplay underlying the spectrum of neoplastic phenotypes.

### Tumor-immune microenvironment is dominated by diverse myeloid populations

After defining the neoplastic cell states in pHGG, we next sought to characterize the immune microenvironment. T cells comprised ∼2% of the cells captured via snRNA-Seq and were primarily found in two post-therapy specimens (**Figure 1b-d**). The progressive H3G34V-mutant case was a notable outlier, with T cells representing ∼22% of cells captured (**Figure 1d**). A low T cell abundance with outliers up to ∼20% of total cell composition is consistent with adult GBM^23^. Myeloid cells comprised ∼9% of the snRNA-Seq data and were captured in each patient, so we selected this population for further analysis. After reintegration (**Extended Data Fig. 6a-c, Methods**), we identified 11 distinct myeloid populations that were manually annotated based on their differentially expressed genes and transcriptional regulons and demonstrated extensive heterogeneity between patients and therapeutic time points (**Figure 4a-c**, **Extended Data Fig. 6d-h, Supplementary Table 5**). Most samples contained a distribution of myeloid subpopulations, while a few samples were dominated by a single subtype (**Extended Data Fig. 6h**). Additionally, these cells formed a continuous phenotypic spectrum, including resident microglia and bone marrow-derived macrophage ontogenies (**Extended Data Fig. 6d**).

**Figure 6.**
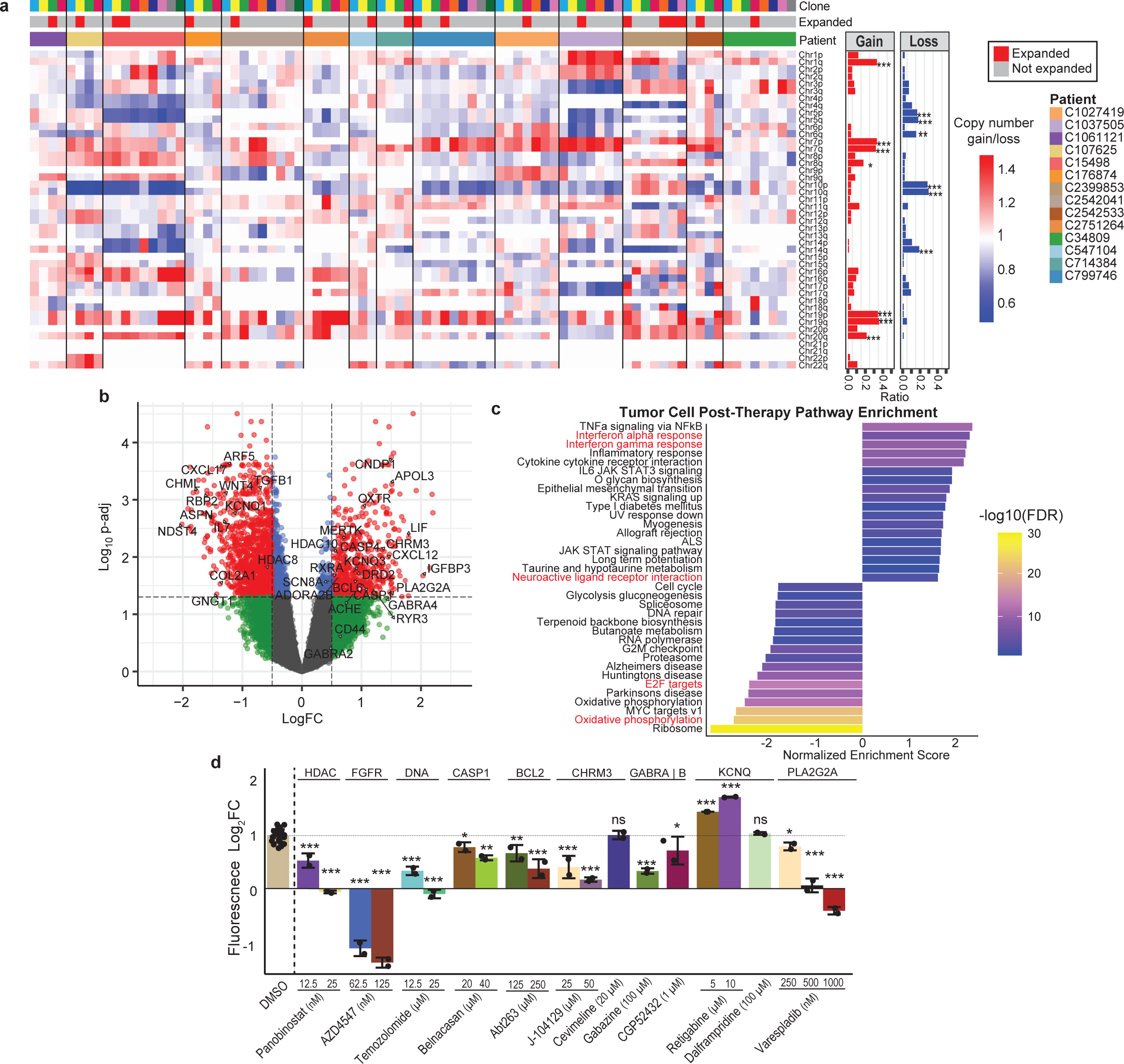
Identifying resistance mechanisms through *in vitro* drug screening. **a)** Left, heatmap of average CNVs within each tumor subclone at the chromosome arm level across 14 patients using Clonalscope. Clone color (top row) corresponds to the patient-specific subclone shown in **Extended Data Figure 9a**. Right, ratio of binarized copy number gain or loss for each chromosome arm, defined as having an average CNV >1.25 or average CNV <0.75 respectively. For each chromosome arm, a Fisher’s exact test was used to assess for recurrent copy number alterations, adjusted using the Benjamini-Hochberg method. *** p<0.001; ** p<0.01, * p<0.05. **b)** A linear mixed model was used to identify differentially expressed genes within neoplastic cells overall between initial resection and post-therapy time points accounting for individual patient variability. Volcano plot shows the log fold change and adjusted p-value for each gene included in the model, with selected genes labeled. **c)** Gene set enrichment analysis (GSEA) of Hallmark and KEGG pathways across all genes in (**b**) ranked by log fold change. **d)** Selected results from *in vitro* drug screening in human pHGG cell lines. Cells were treated with drugs at indicated concentrations, and growth was monitored using a fluorescent reporter 72 hours after drug treatment (*n* = 24 control, 2 drug-treated replicates each). Positive values indicate a net proliferation, while negative values indicate net cell death. Gene target or mechanism of action is indicated above the drugs. Significance is assessed via a two-sided Student’s *t* test for each condition compared to DMSO controls and adjusted for multiple hypothesis testing using the Benjamini-Hochberg method, with mean ± SD shown. *** p<0.0001; ** p<0.01, * p<0.05.

The myeloid subpopulations included tissue-resident microglia, dendritic cells, and multiple tumor-associated macrophage (TAM) subsets that have been previously characterized across multiple solid tumor types including adult glioma^32^. This includes pro-angiogenic TAMs differentially expressing *VEGFA* and glycolytic enzymes (i.e., *HK2, ENO2*), lipid-associated TAMs (*PPARG* and *LPL*), inflammatory TAMs (*NFKB1* and *IL1B*), interferon (IFN)-responsive TAMs (*IFIT2, IFIT3,* and *ISG15*), as well as two additional populations of putative bone marrow-derived macrophages, BMD TAM 1 (*F13A1, TMEM163, MS4A4E*) and BMD TAM 2 (*TGFBI, MALT1, RGS2*). Microglia were primarily stratified into pre-active microglia (*CCL3, EGR3, NFKBID*) and homeostatic microglia (*P2RY12, TMEM119*) (**Figure 4a-c**). Both pre-active and homeostatic microglia populations exhibited a trend of decreasing frequency post therapy in the majority of the samples, while bone-marrow derived macrophages tended to increase post therapy (**Figure 4d**). This is consistent with observations in adult GBM, in which a microglia to macrophage shift post therapy has been reported^23^. Given that myeloid population shifts were highly variable between patients, we applied a generalized linear mixed model approach (**Supplementary Table 6, Methods**) to identify pathway-level changes in pseudobulk myeloid cells during tumor progression. We observed upregulation of interferon and inflammatory response pathways, and downregulation of pathways related to proliferation and cellular metabolism (e.g., oxidative phosphorylation, E2F targets) (**Figure 4e**).

Glioma-associated macrophages have been previously demonstrated to differentially interact with neoplastic cell states, altering their activity and differentiation status^33^. Indeed, we observed differential correlations in frequency between neoplastic cell states and myeloid subtypes across tumor regions. The MES-like state was associated with pro-angiogenic TAMs, while the OPC/NPC-like state was associated with homeostatic microglia (**Figure 4f)**. Consequently, we aimed to elucidate how these pHGG-associated myeloid subpopulations interact with our newly defined pHGG-specific neoplastic cell states through an analysis of inferred ligand-receptor interactions (**Figure 4g, Supplementary Table 7, Methods**). We observed a broad and heterogeneous set of bidirectional cellular interactions. Importantly, myeloid cells were predicted to mediate multiple neoplastic cell functions both through direct contact and secreted factors. These interactions included processes involved in regulating growth and proliferation (e.g., SPP1-CD44, HBEGF-EGFR/ERBB2), cell adhesion and migration (e.g., FN1-ITGA3/ITGB1), and modulation of electrochemical or synaptic properties (e.g., NLGN1-NRXN3) (**Supplementary Fig. 5**). The AC-like 1 and Intermediate 1 cell states were predicted to receive intercellular signals most broadly across myeloid subtypes with receptors including FGFR1, IGF1R, and CD44, which is specifically known to interact with a range of ligands (e.g., HBEGF, PSEN1, SPP1, VEGFA) and has a critical role in adult glioma^34–36^. In contrast, the AC-like 2 and Intermediate 2 populations were the most inert neoplastic populations (**Figure 4g**). Additionally, the NEU-like population was predicted to be a significant ligand source for intra-neoplastic interactions with OPC/NPC-like and NEU-like cells (**Figure 4g**). Taken together, this analysis is the first comprehensive characterization of pHGG myeloid subtypes and suggests that TAM populations can differentially interact with neoplastic cell states and modulate multiple neoplastic cell intrinsic functions.

### Mapping the spatial landscape of pHGG

Gliomas are not only highly heterogeneous in terms of cell types and states, but complex topographic localization of neoplastic and immune populations yields spatial niches with distinct molecular functions and therapeutic vulnerabilites^37–41^. To characterize the spatial landscape of pHGG, we employed Co-Detection by Indexing (CODEX) spatial proteomics with a 52-plex panel (51 antibodies + DAPI) on 11 whole-slide formalin-fixed paraffin-embedded (FFPE) samples that had paired snRNA-Seq data, including three patient-matched longitudinal pairs (**Figure 5a, Supplementary Table 8, 9**). First, we confirmed appropriate antibody staining morphology and co-localization, and we manually removed areas with staining artifact (**Supplementary Fig. 6**). Then, we confirmed that our CODEX panel was able to resolve gross anatomical compartments including bulk tumor, gray matter, and white matter (**Figure 5b**). Finally, after segmenting single cells, computational integration, and clustering, we annotated over 7.5 million single cells (**Figure 5c-d, Supplementary Figure 7, Extended Data Fig. 7a-c, Methods**). We captured the primary axis of neoplastic cell states from proneural (high expression of SOX2, OLIG1, OLIG2) to mesenchymal/astrocytic (high expression of CD44, VIM, GFAP). Interestingly, we observed two distinct MES (mesenchymal)-like tumor populations. MES-like-1 tumor cells expressed the additional mesenchymal markers APOE, SPP1, and GLUT1, and were predominantly identified in peri-necrotic regions (**Supplementary Figure 7e**), and MES-like-2 tumor cells had the highest expression of the canonical marker CD44 (**Extended Data Fig. 7b-c**). Neoplastic cell states were distributed heterogeneously both between and within samples with regions of the tumor predominated by patches of either proneural or mesenchymal tumor cells (**Figure 5e**).

The immune populations were predominated by myeloid cells, consistent with the sequencing data, and similarly formed a continuous phenotypic distribution including microglia and macrophages (**Figure 5c, f**). We identified a macrophage population that strongly co-expressed classically immune suppressive markers CD163 and CD206, a second macrophage population characterized by high HLA-DR expression, and a large population of MPO^+^ myeloid cells (**Figure 5c**, **Extended Data Fig. 7b**). This MPO^+^ population had a high expression of HIF- 1A and was primarily found as large infiltrates in necrotic regions in several samples (**Supplementary Figure 8a, Extended Data Fig. 7b**). While we identified small populations of CD4^+^ and CD8^+^ T cells, inspection of the images revealed that T cells were predominantly located within vessels or concentrated in areas of hemorrhage. This demonstrates that blood contaminants in tissue may confound analysis of single-cell sequencing of rare immune populations (**Supplementary Fig. 8b)**. Lastly, we observed spatially restricted expression of immune checkpoint molecules including CD47 and PD-L1 (**Supplementary Fig. 8c)**.

### Myeloid cells are spatially colocalized with distinct tumor states

To systematically identify recurrent spatial patterns, we performed unsupervised neighborhood analysis and identified 15 cellular neighborhoods (CNs) which we manually annotated based on their relative enrichment of cell types (**Figure 5g**). These neighborhoods were heterogeneously distributed across samples and captured expected anatomic compartments including gray matter (CN2, predominantly mature neurons), white matter (CN6, mature oligodendrocytes), and infiltrating tumor regions (CN7, normal oligodendrocytes and tumor cells) (**Figure 5h, Supplementary Fig. 8)**. Additionally, this analysis highlighted localized regions predominated by different tumor cell states (i.e., proneural, intermediate, and mesenchymal neighborhoods), as well as MPO^+^ infiltrates, and a vascular neighborhood (**Figure 5h**). Tumor cells tended to co-localize with cells sharing the same phenotype, such as proneural tumor cells localizing with other proneural tumor cells. (**Figure 5e, g**). Notably, each mesenchymal tumor cell type was primarily enriched in its own cellular neighborhood (CN3, CN15) with a relative depletion of proneural or intermediate tumor cells, suggesting that mesenchymal tumor cells form localized niches that are distinct from other regions of bulk tumor (**Figure 5g**).

Immune cells were differentially localized across cellular neighborhoods. Microglia were enriched in areas of normal brain, primarily gray matter (**Figure 5g**), and were observed to be concentrated at the tumor-normal boundary (**Supplementary Fig. 8d**). Macrophages and T cells were jointly enriched in an immune-predominant neighborhood (CN1), a perivascular neighborhood (CN5), a vascular tumor neighborhood (CN11), and a tumor/immune neighborhood (CN9). Outside of these neighborhoods, unclassified macrophages were most enriched in the MES-like-1 neighborhood (CN15). Immune cells were relatively depleted in all neighborhoods that had a significant enrichment of proneural tumor cells (**Figure 5h**). Together, this is consistent with previous reports that macrophages are most enriched in the vicinity of MES-like glioma cells^33^, but further suggests that this is specific to some TAM subpopulations. This was supported by examining distances from tumor cells to myeloid cells, which revealed that MES-like-1 tumor cells were consistently enriched near MPO+ myeloid cells and unclassified macrophages, while MES-like-2 tumor cells and proneural tumor cells were both enriched near HLA-hi macrophages (**Extended Data Fig. 8a-e**). This analysis also revealed that MES-like-1 tumor cells were the furthest population from vasculature, while MES-like-2 tumor cells were closest to vasculature after immune cells, supporting a hypoxia-dependent stratification of mesenchymal cell states (**Extended Data** Fig. 8a, f)^12^.

### Tumor subclone dynamics reveal recurrent genomic alterations

We next aimed to apply our longitudinal data to identify mechanisms of therapeutic resistance. We first utilized large-scale copy number variations (CNVs) to trace tumor subclones across patient-matched samples with Clonalscope^42^ which integrates snRNA-Seq and matched WGS data. Neoplastic subclones were defined at the earliest time point for each patient and traced to the later therapeutic time points to assess populations that have expanded or regressed during treatment and progression (**Methods**). Through this approach we identified lineage-traced neoplastic subclones on 14 patients, ranging from 4 to 9 subclones per patient, with variable clonal dynamics across time points (**Extended Data Fig. 9a**). We identified recurrent CNVs including copy number gains on chromosomes 1q, 7p/7q, 8q, 19p/19q, and 20p and copy number losses on chromosomes 5p/5q, 6q, 10p/10q, and 14q (**Figure 6a**). Clustering of gene-level CNVs across tumor subclones revealed recurrent modules of highly correlated CNVs across patients, indicating that similar patterns of chromosome alterations dynamics occur during disease progression **(Extended Data Fig. 9b**). Notably, some alterations occurred more frequently on expanded subclones (e.g., gain on chr18p and chr19q, loss on chr14q), suggesting that these alterations may confer a survival advantage (**Extended Data Fig. 9c, Supplementary Table S10**).

### Longitudinal analysis uncovers tumor cell-intrinsic targets

We then applied an analogous generalized linear model approach (**Methods**) to identify genes and pathways that were upregulated across therapeutic time points over all neoplastic cells, accounting for individual patient variability. Despite not observing population-level cell state shifts in the neoplastic compartment, this analysis yielded 627 significantly upregulated genes and 1,551 significantly downregulated genes (adjusted p <0.05) (**Supplementary Table 11, Figure 6b**). Examining pathway-level changes revealed an upregulation of type I and type II interferon response pathways and the neuroactive ligand-receptor interactions gene set as well as downregulation of pathways related to cell proliferation and metabolism, primarily oxidative phosphorylation (**Figure 6c**).

We then aimed to utilize this neoplastic cell-specific longitudinal analysis to identify and validate tumor cell intrinsic drug targets for pHGG, assuming that consistently upregulated genes are related to therapy resistance. To prioritize gene targets, we screened differentially upregulated genes against multiple drug target databases and the Cancer Dependency Map (DepMap) as well considered their roles as receptors in the tumor microenvironment (**Extended Data Fig. 10a-c, Supplementary Tables 7, 12, 13, Methods**). After curating targets to validate, we screened over 20 pharmacological compounds to assess their impact on cell proliferation in a pHGG post-therapy cell line. Cell proliferation and viability were assessed via a fluorescent reporter 72 hours after drug treatment as a fold change of fluorescence intensity from the time of drug treatment and compared to the growth fold change of DMSO controls (**Methods**). We verified a cytotoxic effect of panobinostat (non-selective HDAC inhibitor), AZD4547 (FGFR inhibitor), and temozolomide (alkylating chemotherapy agent). Multiple genes related to apoptosis, pyroptosis, and inflammasome activation were upregulated including caspases (*CASP1, CASP4*), *BCL2L1,* and *BCL6.* Indeed, inhibition of CASP1 with belnacasan and inhibition of BCL-2 with ABT-263 reduced proliferation *in vitro* compared to DMSO controls **(Figure 6d**). Consistent with our pathway analysis, we observed significant upregulation of multiple genes involved in electrochemical and synaptic communication, which has been shown to support glioma progression and invasion^43,44^. This includes receptors for neurotransmitters (e.g., the top predicted target, CHRM3) as well as solute and ion (sodium and potassium) channels. Modulating their functions *in vitro* with small molecule antagonists and agonists confirmed the significance of electrochemical signaling in regulating pediatric glioma cell growth and survival. The selective CHRM3 antagonist, J-104129, resulted in significant cell death, although the cholinergic agonist, cevimeline, had no effect on proliferation. The selective GABA_A_ receptor antagonist, gabazine, and to a lesser extent, the selective GABA_B_ receptor antagonist, CGP52432, had a mild antiproliferative effect (**Figure 6d**). Interestingly, activating KCNQ potassium channels with retigabine significantly stimulated cell proliferation, while inhibiting KCNQ channels (4-aminopyridine) had no effect. Lastly, our screening nominated secretory phospholipase A2 (sPLA2) as a novel therapeutic target in pediatric glioma, with a dose-dependent cytotoxic response upon treatment with the PLA2-inhibitor varespladib (**Figure 6d**).

## Discussion

In this study, we profiled the single-cell transcriptional, chromatin-accessibility, and spatial landscape of pediatric high-grade glioma (pHGG) longitudinally under standard therapy. We defined a set of pediatric neoplastic cell states and identified their transcriptional regulatory networks. Similarly, we characterized the tumor immune microenvironment and identified a diverse spectrum of tumor-associated macrophage (TAM) subtypes and employed a 51-marker CODEX panel that revealed differential tumor-immune co-localization.

The longitudinal patient-matched samples provide critical insight into the molecular mechanisms of tumor progression and therapy. Mesenchymal transformation has been described as a hallmark of progressive GBM, analogous to epithelial-to-mesenchymal transition in carcinomas^23,25,45,46^. In pHGG, we indeed observed a spectrum of proneural to mesenchymal differentiation states that resembled those characterized in adult glioma. Interestingly, we also observed distinct astrocyte-like states on each end of the differentiation hierarchy, suggesting that astrocytic programs are maintained in a subset of stem-like pHGG neoplastic cells. Additionally, we did not identify any significant shifts in neoplastic cell states, suggesting an important distinction from adult GBM.

Our framework for identifying tumor cell-intrinsic drug targets implicated several mechanisms of therapy resistance and uncovered novel targets. Synaptic electrochemical signaling through multiple receptors has been increasingly implicated in adult and pediatric glioma progression^44^, including acetylcholine^47–49^, dopamine^50–52^, and GABA^53–55^. We found that neuroactive signaling is broadly upregulated in pHGG, suggesting that tumor cells may become increasingly dependent on synaptic activity over time. We also identified sPLA2 as a novel target in pHGG. Phospholipases are enzymes that hydrolyze phospholipids into precursor fatty acids, which have roles in cell signaling, metabolism, and inflammation. Phospholipase A2 has been implicated in multiple cancer types including colorectal cancer^56^, skin cancer^57^, and adult glioblastoma^58,59^, in which it has been shown to inhibit apoptosis and activate EGFR signaling^60^.

Overall, our study sheds light onto the molecular mechanisms of pHGG, but there are important limitations. Primarily, the small size and frequent inoperability of these tumors necessitate a relatively small and heterogeneous cohort with some samples collected post-mortem. Thus, additional profiling is necessary to elucidate the specific effects of different molecular subtypes and chemotherapeutic agents on longitudinal changes. Crucially, further *in vitro* and *in vivo* studies are expected to elucidate microenvironment-dependent mechanisms of resistance.

### Statistics and reproducibility

No statistical method was used to predetermine sample size. All available longitudinal specimens at the Children’s Hospital of Philadelphia meeting the inclusion criteria were profiled, and all data meeting standard QC thresholds were included. The two-sided Wilcoxon signed-rank test for paired samples was used to compare percentages of cell initial resection and post-therapy specimens. A two-sided Student’s *t* test was used to compare cell growth for *in vitro* experiments. The Fisher’s exact test was used to assess for recurring copy number alterations in the tumor subclone analysis, and a hypergeometric test was used to assess cell type enrichment in spatial neighborhoods. Both were adjusted for multiple hypothesis testing via the Benjamini-Hochberg method. A logistic regression model was used to identify differentially expressed genes in tumor cells across cell states and time points, and the Wilcoxon rank-sum test was used to identify differentially accessible transcription factor motifs in tumor cells and differentially expressed genes and regulons across myeloid cell types and adjusted using the Bonferroni correction. Distance analysis in CODEX data was conducted using a one-sided permutation test (**Methods**).

## Methods

### Human biospecimens

Primary samples were obtained from patients with high-grade glioma banked at the Children’s Hospital of Philadelphia (CHOP) Childhood Cancer Research (CCCR) Registry. The patient selection was built based on specimen availability. Biorepositories were obtained with parent informed consent according to the Declaration of Helsinki and Institutional Review Board approval from all participating centers. All patients underwent an initial tumor resection after histopathological diagnosis of high-grade glioma before receiving treatment, followed by a secondary surgical resection or sample acquisition at autopsy. Germline DNA from either blood or skin samples were acquired from the Children’s Brain Tumor Network (CBTN) at CHOP. Patient sample information and relevant clinical metadata is provided in **Supplemental Table 1**.

### Single-nucleus RNA sequencing (snRNA-Seq)

Single nuclei suspensions immediately underwent library preparation using the Chromium Single Cell 3’ Reagent Kit v3 or V3.1 (10x Genomics) according to the manufacturer’s instructions. Library quality was assessed using the Bioanalyzer Agilent 2100 with the High Sensitivity DNA chip (Agilent Technologies, 5067-4626). Indexed libraries were pooled and sequenced on an Illumina NovaSeq 6000 using sequencing parameters 28:8:0:87 (read1:i5:i7:read2, bp) with an average sequencing depth of 50,000 read pairs per nucleus.

### Single-nucleus assay for transposase-accessible chromatin using sequencing (snATAC-Seq)

Single nuclei suspensions immediately underwent library preparation with the Chromium Next GEM Single Cell ATAC Reagent kit V1.1 (10x Genomics) as per manufacturer’s user manual. Library quality was assessed using the Bioanalyzer Agilent 2100 with a High Sensitivity DNA chip (Agilent Technologies, 5067-4626). Indexed libraries were pooled and sequenced on an Illumina NovaSeq 6000 using sequencing parameters 49:8:16:49 (read1:i5:i7:read2, bp) with an average sequencing depth of 50,000 read pairs per nucleus.

### Processing and quality control filtering of snRNA-Seq data

Read count matrices for snRNA-Seq data were generated from raw FASTQ files using Cell Ranger v3.1.0. Reads were aligned to the GENCODE Release 34 (GRCh38.p13) transcriptome reference. The resulting count matrices were processed and analyzed using Seurat v4^61^. Quality control filtering was applied to each cell, using filters of 500 < nFeature_RNA < 8000 and mitochondrial read percentage < 10%. Poor quality samples containing fewer than 500 cells passing quality control thresholds were excluded from downstream analysis. For three samples of borderline but passable quality (7316-339, 7316-7545, and 7316-7559) we instead used filters of 300 < nFeature_RNA < 8000 and mitochondrial read percentage < 20%. Doublets were called and removed using the DoubletFinder package (v3)^62^ using a doublet proportion estimate of 7.5%.

### Sample integration, clustering, and cell type annotation of snRNA-Seq data

Initial snRNA-Seq data processing was performed using the Seurat v4 package. To aid in identification of malignant and non-malignant cell populations, two published high-grade glioma snRNA-Seq datasets^22,63^ were included with snRNA-Seq data from the present study in the following integration and annotation protocol. Due to memory constraints in Seurat v4, data were randomly downsampled so as not to exceed 200,000 total cells, preferentially downsampling cells from samples with a higher cell count to preserve cells in samples with lower cell counts. Cell cycle scores were computed using the *CellCycleScoring* method with annotated cell cycle genes (2019 update). Integration was performed by reciprocal principal component analysis (RPCA) at a patient level. In detail, each patient was normalized by *SCTransform* (v2) with regression of mitochondrial percentage, S score, and G2M score by Gamma-Poisson generalized linear model. A total of 3,000 features were chosen by *SelectIntegrationFeatures* followed by PCA. The *FindIntegrationAnchors* function was run using top 30 PCs. Following integration, PCA was repeated on integrated features with *RunUMAP* and *FindNeighbors* computed using the top 30 PCs. Louvain clustering was performed by *FindClusters* at a resolution of 0.6. Cluster annotation was performed by manual review of canonical cell type-defining genes, allowing for identification of normal cell type populations including immune and stromal cells as well as a heterogenous and admixed population of other neural and glial cells whose neoplastic versus normal status was inferred by downstream copy number alteration analysis.

As increased computational capacity became possible, after annotation using the downsampled data, the remaining cells were added to the downsampled dataset. These cells were first normalized with *SCTransform* and integrated through projection with Seurat v5^64^ using *FindTransferAnchors* with *dims* = 1:50, followed by *MapQuery* using the integrated PCA and integrated assay with default parameters. To support the cell type annotations in the full dataset, we projected the entire dataset onto an integrated reference atlas of adult glioblastoma^65^. Briefly, the reference atlas was log normalized, and a PCA was recomputed using the published variable features. The data was projected using the *FindTransferAnchors* function with *dims =* 1:50, followed by the *TransferData* function with default parameters in Seurat v5.

### Inference of neoplastic versus normal cells by copy number alteration analysis

Neoplastic versus normal cell annotation was inferred by the presence or absence, respectively, of copy number alterations (CNA) detected from snRNA-Seq data using a dockerized implementation of InferCNV^66^ (https://hub.docker.com/r/trinityctat/infercnv, version tag 1.11.1). Due to computational constraints, the downsampled dataset as described above was used for all analysis of neoplastic cells in the snRNA-Seq data. Input parameters included *cutoff* = 0.1 (recommended for 10x Genomics snRNA-Seq data), as well as *cluster_by_groups* = FALSE and *analysis_mode* = "subclusters’’ in order to cluster cells by distinct copy number profiles. All samples for a given patient were run together in order to capture CNA clusters that may be shared between different tumor regions or timepoints. Unambiguous normal cell clusters identified during the Seurat integrated analysis of snRNA-Seq data were aggregated into three separate normal cell categories (specifically, mature neuron/glial, white blood cells, and vascular cells) which were then used as normal reference populations for InferCNV. Note that aggregation was required in order to meet the minimum cell count requirement for InferCNV across all patients. The remaining non-reference cells were annotated as “neoplastic” if the CNA profile of their corresponding InferCNV cluster matched CNAs detected by WGS from the same patient and were considered to be “normal” otherwise. This comparison with WGS data, performed manually, was an additional quality control step to ensure that putative CNAs inferred from snRNA-Seq match true CNAs detected by WGS of DNA.

### snATAC-Seq data processing

snATAC-Seq data for each sample was first demultiplexed using CellRanger-ATAC v.1.1.0 (10x Genomics). The fastq files were then processed using the *process* module of scATAC-pro (v1.4.4)^67^ with the default parameters. Briefly, the raw reads were aligned to the hg38 genome assembly. Peaks were called using MACS2^68^. Barcodes with more than 2,000 total fragments, < 20% mitochondrial reads, and >25% fraction of reads in peaks (FRiP) were identified as cells. The peak-by-cell count matrix was constructed and used for downstream analyses.

### snATAC-Seq data integration

To integrate data from all patients, we first merged the peaks from different samples if two peaks are within 500bp of each other by the *mergePeaks* module of scATAC-pro. The peak-by-cell count matrix was then reconstructed based on the merged peaks using the *reConstMtx* module of scATAC-pro. Matrices from all samples were concatenated and loaded into Seurat with an extra *ChromatinAssay* added. The data was processed using Signac^69^ as follows: The Seurat object was split by sample ID and each sample was then processed through *FindTopFeatures* (with the minimum cutoff equal to 1% the number of cells present in the subset), *RunTFIDF* and *RunSVD* of Signac. *FindIntegrationAnchors* function was run with parameters *reduction=rlsi* and *dims* = 2:30 with samples 4036 and 4037 as reference, which were from the patient with the greatest number of immune cells as found in the snRNA-Seq data. The anchor features were defined as peaks that are accessible in more than 3% of cells in at least one of the patients. Then, Signac *IntegrateEmbedding* function was run with default parameters. The cells were further clustered with the *FindNeighbors* and *FindClusters* (with *resolution* = 0.8) functions in Seurat. For visualization, the UMAP was constructed using *RunUMAP* with *reduction* = “integrated_lsi” and *dims* = 2:30.

### Construction of transcriptional regulatory network

The transcriptional regulatory network for each neoplastic cell state was constructed as previously described^70^ with minor modifications. We first co-embedded the snATAC-Seq and snRNA-Seq data per sample using the standard Seurat pipeline. Then we identified metacells using hdWGCNA^71^ with parameters *k*=20, *max_shared* = 5, *min_cells* = 50, *reduction* = “pca” and *ident.group* = “seurat_clusters.” Metacells containing between 4-16 snRNA-Seq cells were kept for further analysis. The gene-by-metacell expression matrix and the peak-by-metacell accessibility matrix were calculated as the average normalized expression and normalized accessibility of all cells within the metacell, respectively. Metacells from different samples were then combined and the Enhancer-Promoter (EP) interactions were predicted using a linear regression model for each gene on metacells, with the gene expression in each metacell as the dependent variable, and the accessibility of the peaks within +/- 500kb of the gene promoter as the independent variables. Significant EP interactions were defined based on a peak regression coefficient > 0.1 and Benjamini-Hochberg-adjusted p-value < 0.05. Predicted TF-target genes pairs were defined if the TF motif was present at the enhancer of a predicted EP interaction and both the TF and target gene were expressed in at least 20% of the cells within a given cell state.

### Cell-cell communication analysis

To assess ligand-receptor interactions between cell populations in the tumor microenvironment, we implemented the Ligand-Receptor Analysis Framework (LIANA) v0.1.12^72^, which infers cell-cell communication using a consensus of 16 cell signaling database resources and 5 CCC methods (Natmi^73^, Connectome^74^, LogFC Mean, SingleCellSignalR^75^, CellphoneDB^76^) with default parameters. The neoplastic cell states and myeloid subpopulations as annotated above along with the remaining non-neoplastic populations were included. We considered the consensus rank generated via Roust Rank Aggregation as the significance p value to predict the intercellular crosstalk between each pair based on the expression level of known receptors and ligands in the respective clusters and filtered interactions to those with p-value < 0.05. The number of significant interactions between cell populations was quantified, and the most relevant interactions were manually selected to plot.

### Malignant subclone analysis

To study the evolution of malignant subclones in the patient-matched longitudinal samples, we applied Clonalscope (v1.0.0)^42^, which utilizes both snRNA-Seq data and paired WGS data. Clonalscope identifies copy number variation (CNV) segments with a Hidden Markov Model (HMM) from the paired WGS data, and then estimates the fold change of CNV segments at a single cell level using the snRNA-Seq data with a Poisson model. Then, it identifies tumor subclones through a Bayesian non-parametric clustering process based on the estimated CNVs. Clonalscope was run with default parameters on 14 of the 16 patients. Patient C70848 was excluded due to having a single time point and patient C1060383 was excluded due to an insufficient amount of non-neoplastic cells for the Clonalscope algorithm. The required normal/reference cells were defined by manual inspection of the inferCNV profiles as described above. Paired WGS data augments identification of CNV segments, improving malignant subclone delineation. WGS data was first analyzed by CNVkit as described and iteratively refined by (1) merging continuous segments that share the same copy number state (amplification, neutral, or loss) and (2) merging each with neighboring segments if its size is <5% of both neighboring segments and if both neighboring segments share the same copy number state. This process denoises the WGS-defined CNV segments for use with Clonalscope.

Clonalscope was then applied to estimate CNV profiles of single cells at the earliest time point for each patient, through a non-parametric clustering process. The estimated mean CNV profile of each subclone is utilized as a prior to trace similar subclones or discover new subclones from subsequent time points. For each patient, the shifts in malignant subclone proportions were visualized using clevRvis (v0.99.6)^77^, with the *fishPlot* function using a spline fit. Clones were defined as having expanded if their percentages increased over time and comprised at least 10% of the malignant population at the latest time point. The average values of estimated CNVs were summarized for each chromosome arm. CNV gain or loss was binarized as follows: average CNV >1.25 was defined as a copy number gain and <0.75 as a copy number loss. For each chromosome arm, a Fisher’s exact test was used to assess for recurring copy number alterations comparing the gain or loss of each chromosome segment relative to the gain or loss of all other segments and adjusted using the Benjamini-Hochberg method.

### CODEX staining

CODEX staining was done using the sample kit for PhenoCycler-Fusion (Akoya, 7000017) according to Akoya’s PhenoCycler-Fusion user guide with modifications to include a photobleaching step and overnight incubation with antibodies at 4°C. FFPE samples were sectioned at 5 μm thickness and mounted onto charged slides (Leica, 3800080). Sample slides were baked overnight at 60°C and allowed to cool to room temperature. Sample slides were deparaffinized in Xylenes (Sigma, 534056) twice and rehydrated in a graded series of ethanol concentrations (2 times 100%, 90%, 70%, 50%, 30% and 4 times ddH2O). Antigen retrieval was performed in 1x Dako Target Retrieval Solution, pH 9 (Dako, S2367) with a pressure cooker for 20 minutes. After equilibrating to room temperature, sample slides were washed 2 times with ddH2O and once with 1x PBS before being submerged in a four-well plate containing 4.5% H2O2 and 20mM NaOH in PBS (bleaching solution) for photobleaching. The four-well plate was sandwiched between two broad-spectrum LED light sources for 45 minutes at 4°C. After 45 minutes, sample slides were transferred to a new four-well plate with freshly-made bleaching solution and photobleached for another 45 minutes at 4°C. Sample slides were washed 3 times in PBS and then 2 times in hydration buffer. Sample slides were equilibrated in staining buffer for 30 minutes and incubated in the antibodies (Supplemental Table S5) diluted in staining buffer plus N Blocker, G Blocker, J Blocker, and S Blocker overnight at 4°C. After antibody incubation, sample slides were washed 2 times in Staining Buffer and fixed for 10 minutes in 1.6% paraformaldehyde (Electron Microscopy Sciences, 15710) storage buffer. Sample slides were washed 3 times in PBS and incubated in ice cold methanol for 5 minutes. After incubation in methanol, sample slides were washed 3 times in PBS and incubated in final fixative solution (1000uL of PBS + 20uL of Akoya’s final fixation reagent) for 20 minutes at room temperature.

The sample slides were then washed 3 times in PBS and stored in storage buffer prior to imaging.

### CODEX imaging

CODEX reporters were prepared according to Akoya’s PhenoCycler-Fusion user guide and added to a 96-well plate. The PhenoCycler-Fusion experimental template was set up for a CODEX Run using Akoya’s PhenoCycler Experiment Designer software according to Akoya’s PhenoCycler-Fusion user guide. Details on the order of fluorescent CODEX Barcodes and microscope exposure times can be found in **Supplemental Table S3**. The PhenoCycler-Fusion experimental run was performed using Akoya’s Fusion 1.0.8 software according to Akoya’s PhenoImager Fusion user guide. Images were taken and pre-processed (stitching, registration, background subtraction) with Akoya’s PhenoImager Fusion microscope using default settings. Final images were evaluated, and selected samples were reimaged with adjusted exposure times based on manual review. After imaging, slides were stained with hematoxylin and eosin (H&E) and imaged at 40x resolution.

### CODEX data segmentation

Nuclear segmentation with a fixed pixel expansion of 4 pixels (equivalent to 2 µm) was performed using Mesmer^78^ for each image to enable the capture of cytoplasmic and membrane markers while limiting lateral spillover. Maxima threshold and interior threshold were each set to 0.3. To generate the necessary input of a two-channel TIFF, we used DAPI for the nuclear channel and a composite channel of GLUT1, CD3e, CD14, and CD68 for the membrane channel although the nuclear segmentation was used. Mean pixel intensity was extracted from each cell segmentation mask, yielding a cell by protein matrix which was carried forward for analysis in Seurat v5^79^. Cells with very low or high raw DAPI expression (<10 or >250 on a UINT8 scale) were removed. Each image was manually cropped to exclude large areas of artifact including tissue folding and detachment, debris, and edge artifact. All marker channels including DAPI, but not blank channels, were retained in the cell by protein matrix of each Seurat object for each sample.

### Cellular neighborhood analysis

Neighborhood analysis was performed as previously described^80^ using the final cell type annotations, and as implemented by the imcRtools package^81^. Briefly, a k-nearest neighbors graph from all cells was constructed using the *buildSpatialGraph* function in imcRtools with k = 20, which calculates the neighborhood composition of each cell with a sliding window. These windows are clustered using k-means clustering with respect to their proportions of cell types with 15 clusters. Statistical significance of cell type enrichment within each neighborhood was calculated using a hypergeometric test. The p-value was calculated based on the following four numbers: (1) the number of cells of a given type in the neighborhood; (2) the total number of cells in the neighborhood; (3) the number of cells of a given type in the CODEX dataset; and (4) the total number of cells in the CODEX dataset. P-values were adjusted for multiple hypothesis testing using the Benjamini-Hochberg method and significance was defined as p-adjusted < 0.001.

### *In vitro* drug screening

Pediatric high-grade glioma cell line 7316-913 was obtained through the Children’s Brain Tumor Network and underwent histopathologic, molecular, and genomic characterization as previously described^82^. Glioma cells were stably transduced with a lentiviral nuclear red fluorescent protein under the EF1a promotor (Sartorius, cat. 4476, Göttingen, Germany) for visualization in live imaging assays. Spheroid cultures were maintained in DMEM/F-12 medium supplemented with 1% glutaMAX (Gibco, cat. 35050061), 100 U/mL penicillin-streptomycin (cat. 15140122), 1X B-27 supplement minus vitamin A (Gibco, Cat. 12587010), 1X N-2 supplement (Gibco, cat. 1752001), 2.5 ng/mL human epidermal growth factor (PeproTech, cat. AF-100-15-B), 2.5 ng/mL human basic fibroblast growth factor (PeproTech, cat. 100-18B), and 0.5μg/mL heparin (StemCell, cat. 07980). Glioma cells were plated at 500 cells per well in 384 well ultra-low attachment plates (S-Bio, cat. MS-9384UZ) in 50μL of media and allowed to form spheroids overnight. Plated cells were subsequently treated with pharmacological compounds in duplicate. Compounds were obtained from the following sources: Selleckchem (Abt263, cat. S1001; Panobinostat, cat. S1030; AZD4547, cat. S2801; Belnacasan, cat. S2228; Temozolomide, cat. S1237; Cevimeline, cat. S6432; Dalfampridine, cat. S5028; CGP52432, cat. S0303; Gabazine, cat. E1247; Retigabine, cat. S4734; Varespladib, cat. S1110), R&D Systems (J-104129, cat. 2507), Thermo Scientific Pierce (DMSO, cat. 20688). Compounds were added as 20μl of a 3.5x working solution for each drug/dilution. Drug concentrations were selected based on prior literature characterizing these compounds in cell line models. Cellular proliferation and viability were monitored via Incucyte Live Imaging technology with imaging every 8 hours.

### Data availability

Data from this study have been deposited at the Human Tumor Atlas Network (HTAN) data portal: https://data.humantumoratlas.org/. For the snRNA-Seq, snATAC-Seq and WGS data this includes sequencing reads and processed data including read alignments, gene-by-cell or peak-by-cell matrices, and variant call files. For the CODEX data, this includes multi-channel images, segmentation masks, and marker-by-cell matrix. For all data types, Seurat objects with annotations and reductions are provided for each data type (shown in **Figures 1, 5**) and subset analyses (shown in **Figures 2, 3, 4)**. The linkage between HTAN patient IDs and sample IDs is provided in **Supplementary Table 1**.

### Code availability

Source code will be made public upon publication, and any code can be made available to the reviewers upon request.

## Acknowledgements

The authors acknowledge the Children’s Brain Tumor Network for collecting tissue and curating clinical information used in this study and the Children’s Hospital of Philadelphia Pathology Core, Flow Cytometry Core, High-Throughput Sequencing Core, and Research Information Services for providing technical support. We are grateful to all the patients and their families who volunteered to participate in this study.

## Author contributions

J.H.S., D.A.O., W.Y., N.A., T.D.R,.K.C., K.T., K.C. conceived and designed the study. S.M., M.S., M.K., A.R., P.B.S., K.C., provided patient samples, J.H.S., D.A.O., W.Y., J.R., A.P.R, J.X., S.B., Y.S., D.W., C.E.H. performed computational and statistical analyses. J.H.S., C.H.C, A.M.Z., A.T., C.E.H., S.B., K.J.A., T.D.R performed experiments. J.H.S., D.A.O., W.Y., J.R., J.X., S.B., A.E.B., R.S.V., C.E.H., N.M.A. performed data interpretation and biological analysis. D.A.O., N.Z., T.D.R., A.N.V., K.C., K.T. provided funding and supervised the study. J.H.S, D.A.O., W.Y., K.T. wrote the manuscript with input from all authors.

## Additional Information

Supplementary information is available for this paper.

Correspondence and requests for materials should be addressed to Kai Tan. Reprints and permissions information is available at www.nature.com/reprints

## Extended Data Figure Legends

**Extended Data Figure 1.**
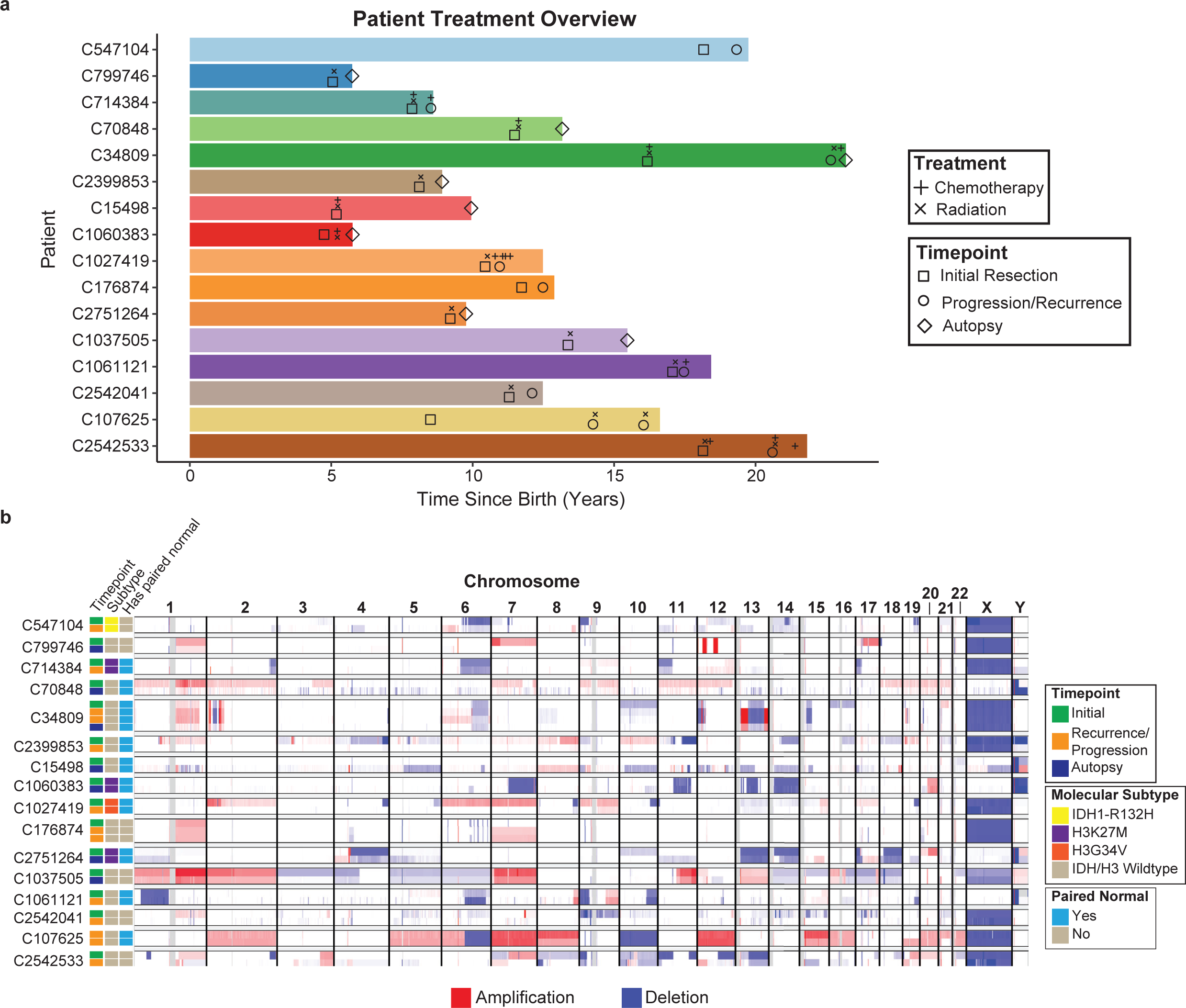
Overview of longitudinal patient cohort. **a)** Timeline of specimen collection and patient treatments if available. Patients were between 4 and 24 years of age. All patients received radiation therapy. Some patients received chemotherapy including temozolomide, bevacizumab, pembrolizumab, vemurafenib, and irinotecan. Samples were collected from an initial resection after histologic diagnosis of high-grade glioma, and then through a secondary post-therapy resection or at autopsy. **b)** Copy number alterations assessed through whole genome sequencing (WGS) for each patient at all available therapeutic time points. Average sequencing depth is 91x per sample.

**Extended Data Figure 2.**
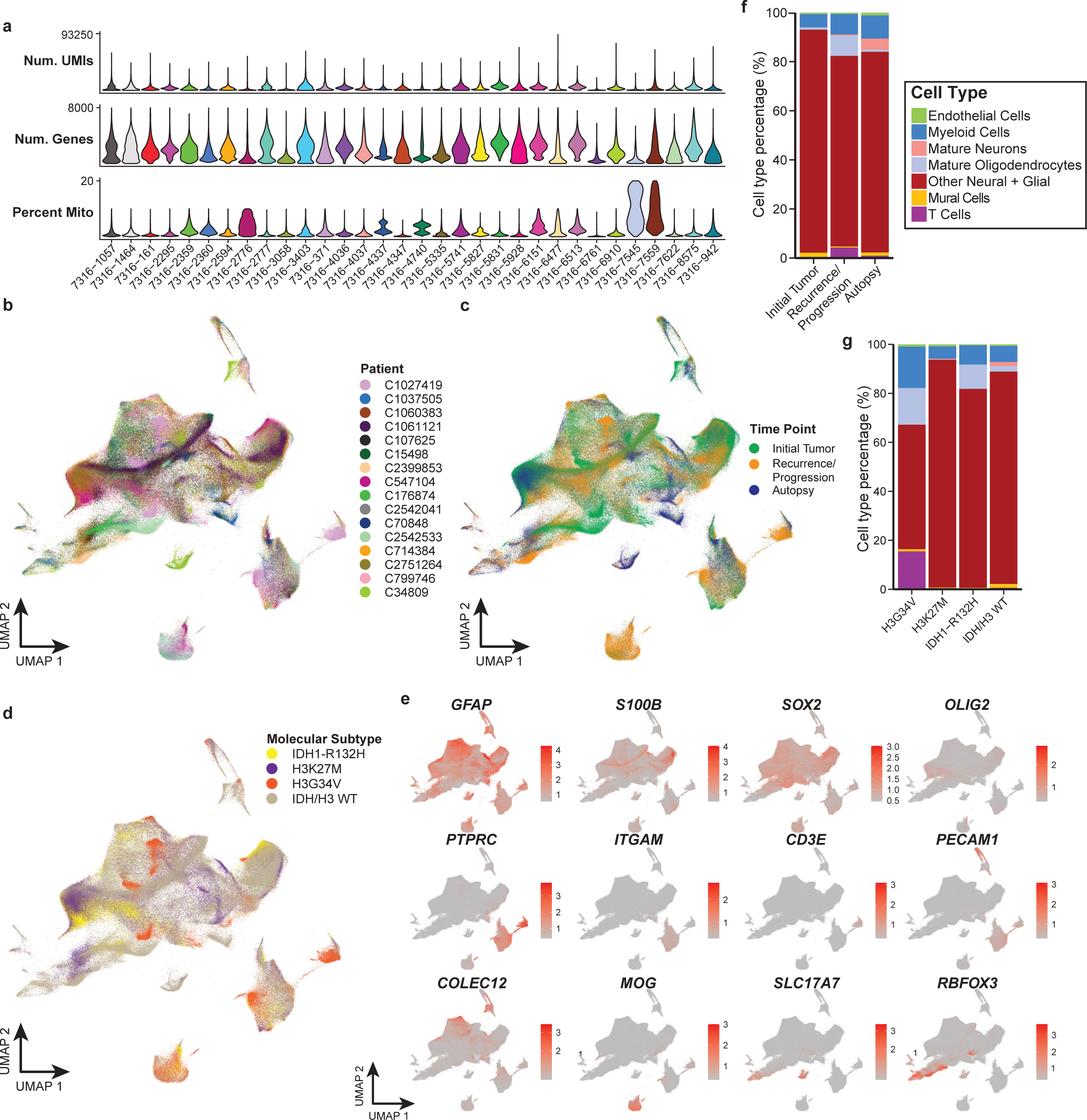
Generation and integration of snRNA-Seq pHGG atlas. **a)** Violin plots of quality control (QC) metrics for each of specimen in the integrated snRNA-Seq dataset. Most specimens were sequenced at two regions, yielding 63 total samples. QC metrics include number of unique molecular identifiers (UMIs), number of unique genes captured after quantitation, and percent of reads originating from mitochondrial genes. **b-d**) Integrated UMAP projection of snRNA-Seq data colored by (**b**) patient, (**c**) time point, and (**d**) molecular subtype. **e)** Expression of marker genes on UMAP of snRNA-Seq data supporting annotation of major cell types. Colors truncated at 1^st^ and 99^th^ percentiles for visualization. **f-g**) Stacked bar plots of cell type proportions across dataset stratified by (**f**) time points separated by initial resection, recurrence/progression (secondary surgical resection), or post-mortem acquisition at autopsy, and (**g**) molecular subtypes defined by driver mutations in IDH or core histone proteins.

**Extended Data Figure 3.**
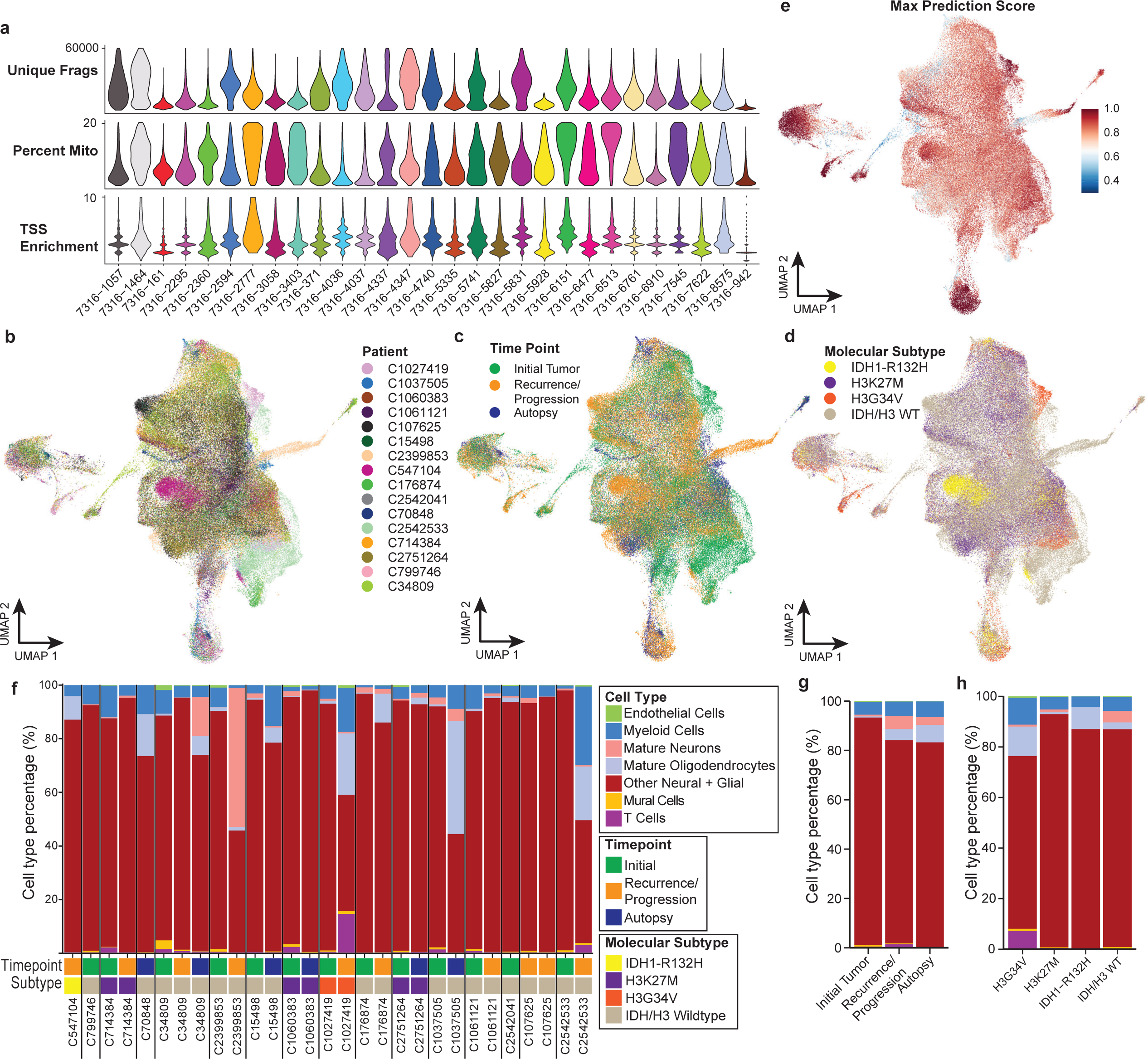
Generation and integration of snATAC-Seq pHGG atlas. **a)** Violin plots of quality control (QC) metrics for each of specimen in the integrated snATAC-Seq dataset (32 total samples). QC metrics include number of unique fragments, mitochondrial genes, and transcription start site (TSS) enrichment of fragment reads in each cell. **b-d**) Integrated UMAP projection of snATAC-Seq data colored by (**b**) patient, (**c**) time point, and (**d**) molecular subtype. **e)** Confidence scores of label transfer predictions using snRNA-Seq to annotate the major cell types in the snATAC-Seq, demonstrating high concordance between the two data modalities. **f)** Cell type proportions in snATAC-Seq data across each patient and therapeutic time point, along with the molecular subtype. **g-h**) Stacked bar plots of cell type proportions across dataset stratified by (**g**) time points separated by initial resection, recurrence/progression (secondary surgical resection), or post-mortem acquisition at autopsy, and (**h**) molecular subtype defined by driving mutations in IDH or core histone proteins.

**Extended Data Figure 4.** Integration and distribution of transcriptionally-defined neoplastic cell states in pHGG. **a-c**) Integrated UMAP projection of inferred neoplastic cells in snRNA-Seq data colored by (**a**) patient, (**b**) time point, and (**c**) molecular subtype. **d)** UMAP of inferred neoplastic cells colored by predicted cell cycle phase. **e)** Predicted neoplastic cell state identity based on canonical cell state modules previously defined by Neftel *et al*.^12^ Cells are assigned to the highest scoring cell state. AC, astrocyte; MES, mesenchymal; NPC, neural progenitor cell; OPC, oligodendrocyte progenitor cell. **f)** Neoplastic cells were computationally projected onto a dataset of the developing fetal human brain^14^. Barplot shows projected cell type proportions for each cell state. tRG, truncated radial glia; uRG, unknown radial glia; IPC, inhibitory neuronal progenitor cell; RG, radial glia; EN, excitatory neuron; IN, interneuron; Astro, astrocyte; GPC; glial progenitor cell; OLC, oligo-lineage cells. **g)** Neoplastic cell state proportions in snRNA-Seq across each patient and therapeutic time point, along with the molecular subtype. **h-i**) Stacked bar plots of cell type proportions across dataset stratified by (**h**) time points separated by initial resection, recurrence/progression (secondary surgical resection), or post-mortem acquisition at autopsy, and (**i**) molecular subtype defined by driving mutations in IDH or core histone proteins.

**Extended Data Figure 5.**
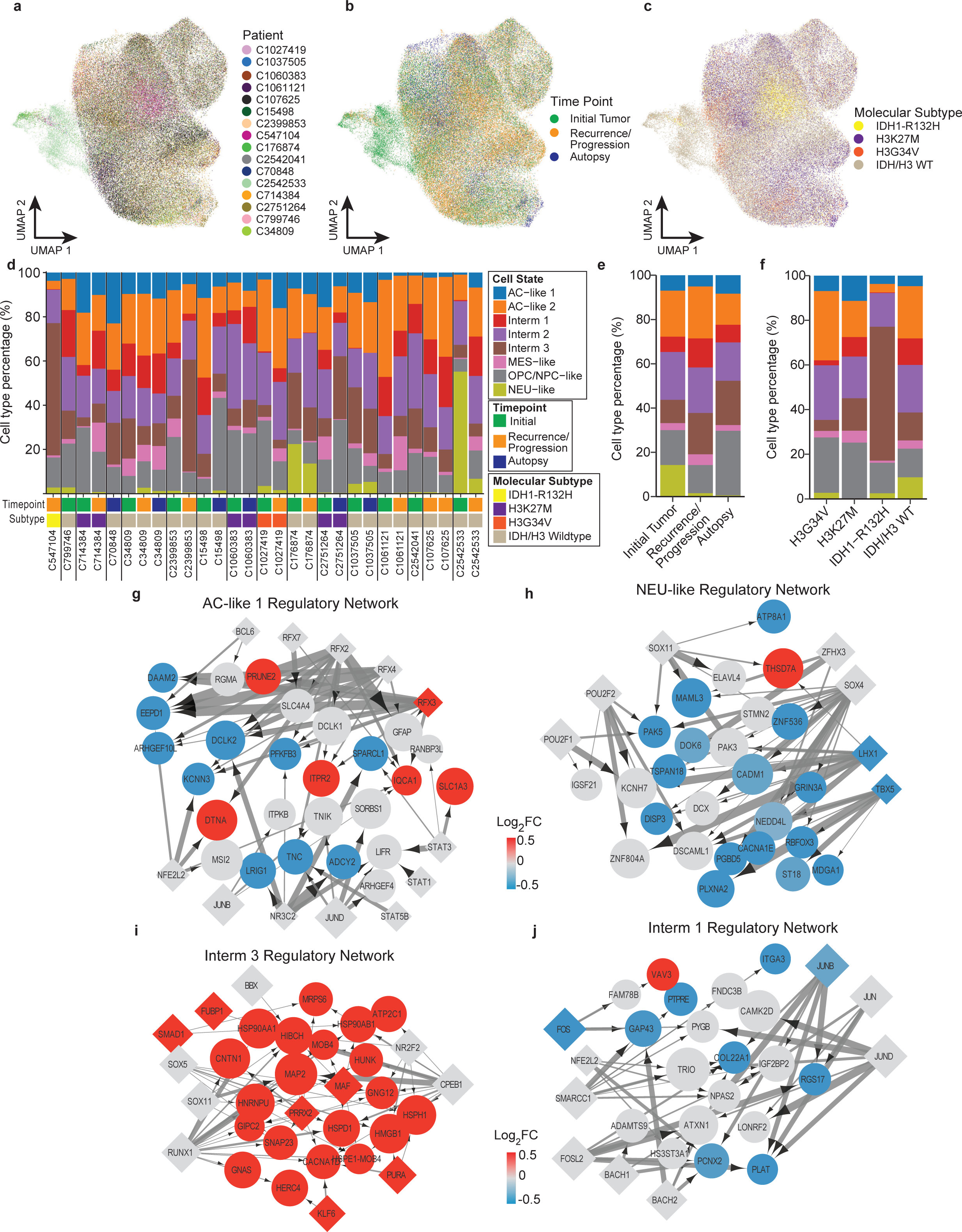
Integration and regulatory network analysis of malignant cell states using snATAC-Seq data. **a-c**) Integrated UMAP projection of inferred neoplastic cells in snATAC-Seq data colored by (**a**) patient, (**b**) time point, and (**c**) molecular subtype class. **d)** Neoplastic cell state proportions in snATAC-Seq across each patient and therapeutic time point, along with the molecular subtype. **e-f**) Stacked bar plots of cell type proportions across dataset stratified by (**e**) time points separated by initial resection, recurrence/progression (secondary surgical resection), or post-mortem acquisition at autopsy, and (**f**) molecular subtype defined by driving mutations in IDH or core histone proteins. **g-j**) Transcriptional regulatory networks for (**g**) AC-like 1 state, (**h**) NEU-like state, (**i**) Interm 3 state, and (**j**) Interm 1 state, showing top 50 upregulated genes and top 15 TFs in each state. Diamond nodes represent transcription factors and circle nodes represent target genes. Node size is proportional to the average gene expression for target genes and average chromVAR z-score for TFs. Node color is proportional to the average log_2_ fold change of the gene in that cell state post-therapy across all cells. Edge line thickness is proportional to the linear regression coefficient for the predicted enhancer-promoter interaction and the fraction of cells with chromatin accessibility at the enhancer peak. The AC-like 2 and Interm 2 state networks are not shown as they include less than 5 significant TF-gene pairs.

**Extended Data Figure 6.**
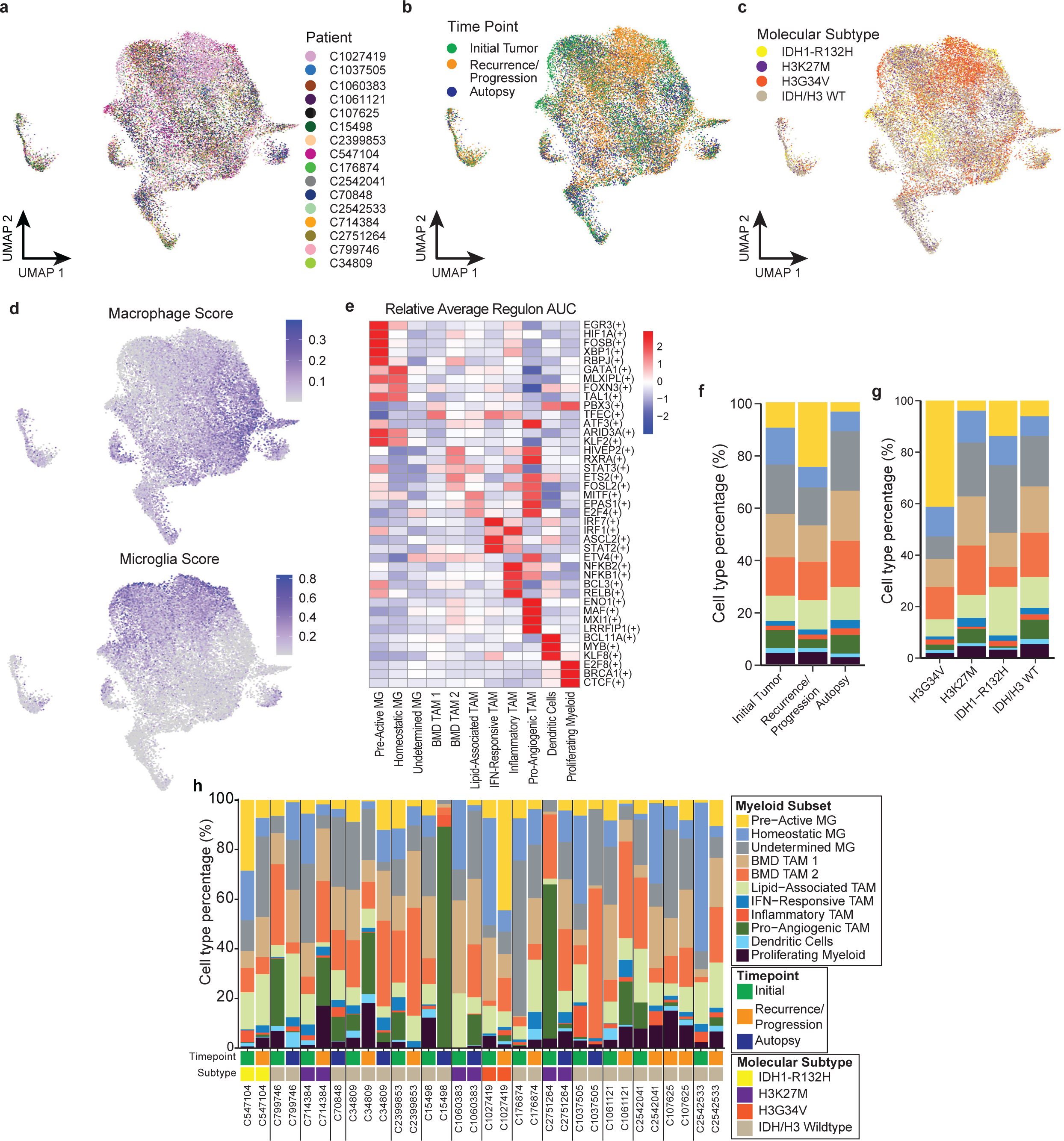
Integration and distribution of myeloid populations in snRNA-Seq. **a-c)** Integrated UMAP projection of myeloid cells in snRNA-Seq data colored by (**a**) patient, (**b**) time point, and (**c**) molecular subtype class. **d)** UMAP colored by signature scores for bone marrow-derived (BMD) macrophages (left) and microglia (right) as previously defined^83^. Colors truncated at 1^st^ and 99^th^ percentiles for visualization. **e)** Transcription factor regulon activity calculated by SCENIC^84^. Heatmap shows average regulon AUC value for top differentially active regulons in each myeloid subpopulation. **f-g**) Stacked bar plots of cell type proportions across dataset stratified by (**f**) time points separated by initial resection, recurrence/progression (secondary surgical resection), or post-mortem acquisition at autopsy, and (**g**) molecular subtype defined by driver mutations in IDH or core histone proteins. **h)** Myeloid cell type proportions in snRNA-Seq across each patient and therapeutic time point, along with the molecular subtype. BMD, bone marrow derived. MG, microglia.

**Extended Data Figure 7.** Integration and annotation of CODEX data. **a)** Left, integrated UMAP projection of all 7.5 million cells in pHGG CODEX atlas colored by sample identifier. Right, pie chart showing the contribution of each sample to the dataset. **b)** Heatmap showing the average centered log ratio (CLR)-normalized expression of each marker per cell type scaled by marker (across rows) and clustered by marker and cell type. Marker names listed in yellow indicate antibodies that had high quality staining on the majority of samples and were subsequently used for integration and clustering. **c)** CODEX images (left) with selected markers and corresponding cell phenotype masks (right) showing appropriate co-labeling of certain markers and examples of annotated cell types. Grey cell masks refer to all other segmented cells in the final analysis (after removal of imaging artifacts and clusters of red blood cells).

**Extended Data Figure 8.**
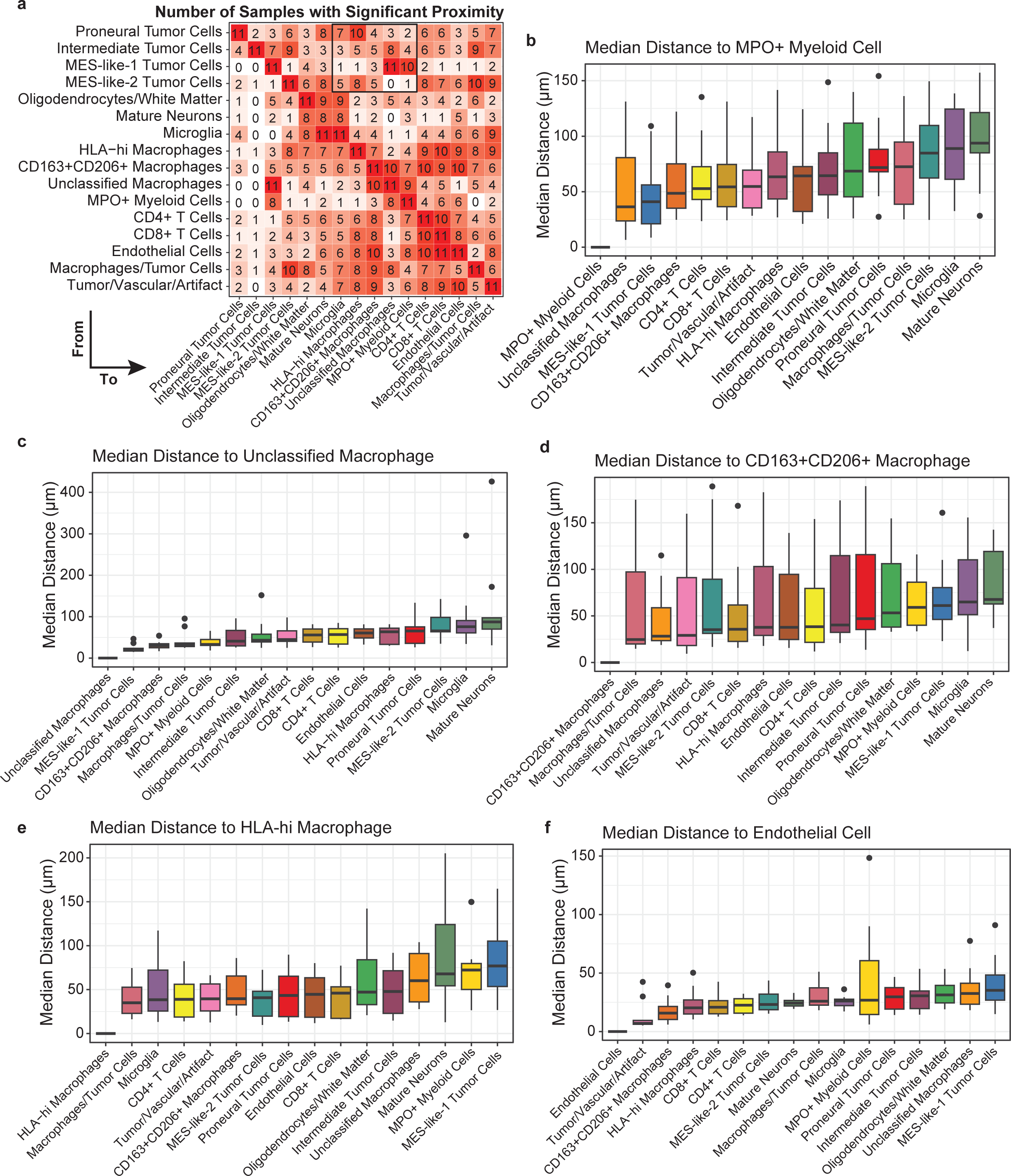
Tumor cell staters are differentially localized near myeloid subtypes. **a)** Heatmap tabulating number of samples (out of 11 total) in which there is a significant proximity of the source cell (rows) to the target cell (columns). Significance was assessed by a one-sided permutation test. Black box highlights proximity from tumor cell states to myeloid subtypes. **b-f**) Median distances in each sample from source cell type (x-axis label) to (**b**) MPO+ myeloid cells, (**c**) unclassified macrophages, (**d**) CD163^+^CD206^+^ macrophages, (**e**) HLA-hi macrophages, and (**f**) endothelial cells.

**Extended Data Figure 9.**
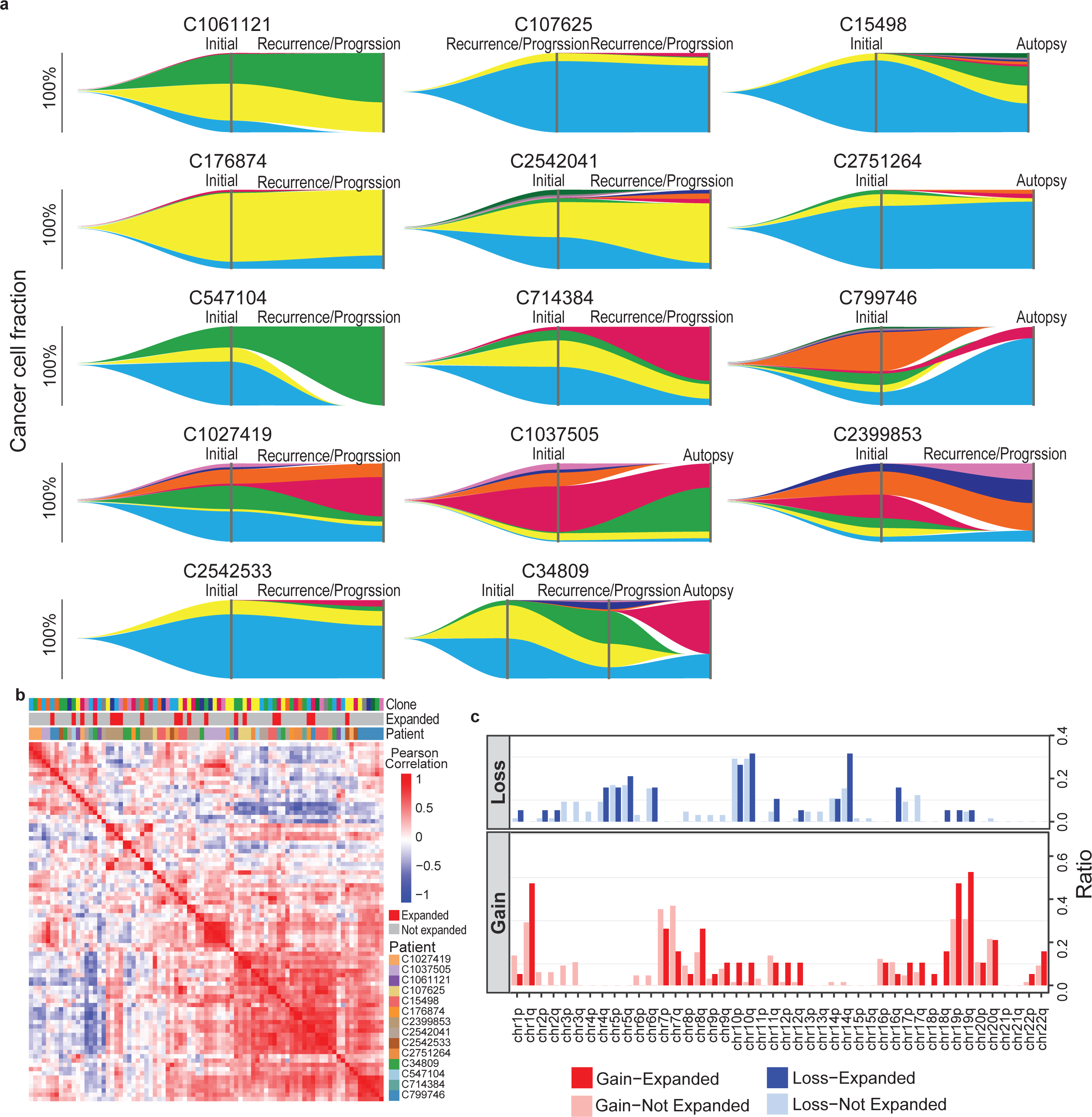
Tumor subclone dynamics across patients. **a)** Fishtail plots showing shifts in tumor subclone populations across longitudinal time points for 14 patients. **b)** Correlation heatmap of subclones based on estimated copy number variation (CNV) at the gene level. The CNV states of chromosome segments in each subclone were used to infer gene-level CNVs for comparison across the cohort. Clone colors indicate subclones identified in (**a**). Expanded subclones are indicated. **c)** Ratio of binarized copy number alteration (gain or loss) at the chromosome arm level comparing expanded and non-expanded clones. The ratio represents the number of subclones with copy number gain or loss among all 84 subclones identified across patients. Expanded subclones were defined if they increased in proportion across time points and comprise at least 10% of the tumor population at the latest time point.

**Extended Data Figure 10.**
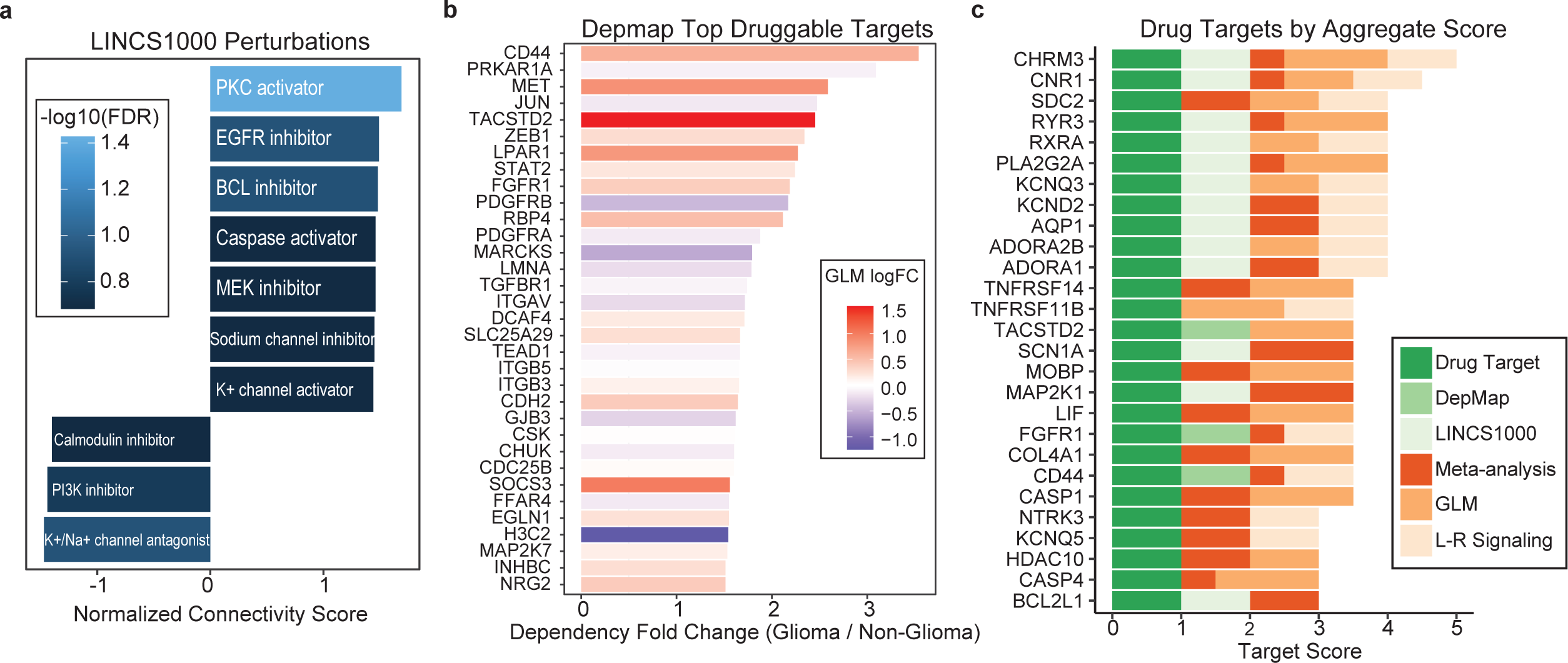
Drug target selection. **a)** Representative drug mechanisms nominated by LINCS1000 using top upregulated and downregulated time point-specific genes (**Methods**). Perturbation results were filtered for false discovery rate <0.25 and normalized connectivity score >0.6. **b)** Top glioma-specific targets predicted from DepMap screening. Dependency scores in glioma versus non-glioma cell lines were ranked by fold change (mean dependency in glioma / mean dependency in non-glioma cell lines). Color indicates log fold change of expression post- therapy using the generalized linear mixed model analysis. **c)** Top gene targets by aggregate ranking score (**Methods**). Criteria includes screening against 3 drug databases, LINCS1000 compound perturbations, and DepMap ,as well as two orthogonal methods of differential expression analysis (time point-specific generalized linear mixed model and per-patient meta-analysis) and participation in ligand-receptor signaling as a receptor target.

