## Supplementary Figures for "A longitudinal single-cell and spatial multiomic atlas of pediatric high-grade glioma"

### Supplementary Table Legends

**Supplementary Table 1.** Chart summarizing available patient characteristics and clinical information of the pHGG cohort profiled in this study, including demographic information, details of histologic diagnosis, reported medical conditions, treatments received and survival times.

**Supplementary Table 2.** List of differentially expressed genes for all annotated neoplastic cell states. A logistic regression model was used comparing normalized gene expression in each cell type against all other cell types. Upregulated and downregulated genes are included with a log2-fold change magnitude threshold of 0.1 and adjusted p-value <0.05.

**Supplementary Table 3**. Top differentially accessible motifs in each neoplastic cell state as calculated by chromVAR. Results are filtered by delta ≤0.01, where delta is the difference between the mean motif deviation in a specific neoplastic cell state and the rest of cells.

**Supplementary Table 4.** Complete metrics of transcriptional regulatory networks for each cell state. This includes peak locations and connectivity strengths of inferred enhancer-promoter interactions and TF-target gene relationships. (**Methods**).

**Supplementary Table 5.** List of differentially overexpressed genes for all annotated myeloid subtypes. A Wilcoxon rank-sum test was used comparing normalized gene expression in each cell subtype across all other myeloid cells, filtered by log fold change of 0.1, minimum expression in 10% of cells., minimum expression difference (cluster versus others) of 10%, and adjusted p-value <0.05.

**Supplementary Table 6.** Output of generalized linear model assessing differentially expressed genes between diagnosis and post therapy time points across entire myeloid population (**Methods**), filtered by adjusted p-value <0.25. Table includes log fold change between time points, average expression across all samples, and statistics based on the Limma-Voom framework (**Methods**).

**Supplementary Table 7.** Results of the LIANA cell-cell interaction analysis. For each significant ligand-receptor pair (robust aggregate rank; p-adjusted <0.05), the result include output metrics and rank from 5 methods (Natmi, Connectome, LogFC Mean, SingleCellSignalR, CellphoneDB), summarizing the expression magnitude and specificity of the gene pairs.

**Supplementary Table 8.** List of antibodies used in this study, including vendors and clone information.

**Supplementary Table 9.** Imaging parameters including the exposure times and cycle/channel assignment for each antibody in the panel.

**Supplementary Table 10.** Copy number variation (CNV) segments of lineage-traced subclones that expanded across therapeutic timepoints, as derived by Clonalscope. Clones were defined as having expanded if their percentages increased over time and comprised at least 10% of the malignant population at the latest time point. CNVs passing the threshold of >1.25 (copy number gain) and <0.75 (copy number loss) are shown for each expanded subclone. Expanded clone ID includes patient identifier and clone number.

**Supplementary Table 11.** Output of generalized linear model assessing differentially expressed genes between diagnosis and post therapy time points across entire neoplastic population (**Methods**), filtered by adjusted p-value <0.25. Table includes log fold change between time points, average expression across all samples, and statistics based on the Limma-Voom framework (**Methods**).

**Supplementary Table 12.** List of top drug perturbations from the LINCS1000 databases. Differentially upregulated and downregulated genes of neoplastic cells across timepoints (**Supplementary Table 1**1) were used as input. Perturbations were filtered for statistical significance (FDR <0.25) and normalized connectivity score (NCS >0.6). For each drug perturbation, table includes the cell line, drug dose, mechanism of action, gene targets, and number of genes whose expression was significantly altered.

**Supplementary Table 13.** Drug target prioritization analysis, filtered for genes passing criteria of 1) being a predicted drug target based on 3 drug target databases and 2) overexpressed in the post-therapy timepoint. Additional columns include differential expression scores and screening against the LINCS1000 and DepMap databases (**Methods**). Columns C-L provide component scores used to calculate drug target score. Columns M-P: results of per-patient meta-analysis; Columns Q-R: results of GLM analysis; Columns T-V: drug database screen; Columns W-Y: dependency scores and fold change from DepMap.

### Supplementary Figure Legends

**Supplementary Figure 1. Full dataset integration and annotation**

- **a)** UMAP projection of full dataset highlighting the additional cells that were included after initial annotation. Due to computational constraints, a downsampled subset of the full snRNA-Seq data was used to integrate samples and annotate major cell types. The remaining data was then projected onto the annotated data set, aligning to the annotated cell types with high confidence. The full snRNA-Seq dataset includes over 400,000 cells. Cells are colored by projection confidence; grey cells indicate initial dataset.
- **b)** Stacked barplot showing proportions of each cell type in the final dataset depending on whether the data was from the initial annotation or projected post-hoc.
- **c)** The full snRNA-Seq pHGG dataset was projected onto an integrated atlas of adult IDH-wild type glioblastoma (Ruiz-Moreno *et al.*). UMAP of cells is colored by predicted cell type. AC-like, MES-like, OPC-like, NPC-like indicate malignant cell states.
- **d)** UMAP colored by projection confidence scores from (**c**).

**Supplementary Figure 2. Annotation of main cell types in snATAC-Seq dataset.**

- **a)** Gene activity scores of key marker genes on UMAP of snATAC-Seq data supporting annotation of major cell types. Colors truncated at 1$^{st}$ and 99$^{th}$ percentiles for visualization.
- **b)** Gene activity signatures of excitatory neurons, astrocytes, and oligo-lineage cells from normal developing brain (Couturier *et al.*) supporting major cell type annotations. Colors truncated at 1$^{st}$ and 99$^{th}$ percentiles for visualization.

**Supplementary Figure 3. Strategy for identifying neoplastic cells using inferCNV.**

**a)** Two representative examples showing the procedure used to identify neoplastic cells in the snRNA-Seq data. Left, raw copy number variation (CNV) profiles from inferCNV for each patient using the snRNA-Seq data. Right, processed CNV profiles binarized as copy number gain, loss, or neutral per region. For each patient, top panel shows reference non-neoplastic cells (immune, vascular, and mature/non-tumor neuroglial cells) and bottom panel shows neuroglial mixture (including putative neoplastic and non-neoplastic cells).

**b)** UMAP projection of snRNA-Seq data with inferred neoplastic cells (left) and normal cells (right) highlighted based on inferCNV results.

**Supplementary Figure 4. Annotation of neoplastic cell states in snATAC-Seq dataset.**

**a)** Gene activity signatures of GBM cell states (Neftel *et al.*) overlaid on UMAP of neoplastic cells. Colors truncated at $1^{st}$ and $99^{th}$ percentiles for visualization.

**b)** Confidence scores of label transfer predictions using snRNA-Seq to annotate neoplastic cell states in the snATAC-Seq data, revealing that the epigenetic landscape of neoplastic cells does not directly reflect the transcriptional state.

**c)** UMAP of neoplastic cells colored by predicted cell cycle phase, based on gene activity scores, demonstrating inability to delineate cell cycle stages in chromatin accessibility sequencing.

**d)** Gene activity signatures of differentially expressed genes for each cell state as determined by snRNA-Seq supporting snATAC-Seq annotation strategy. Colors truncated at $1^{st}$ and $99^{th}$ percentiles for visualization.

**e)** Gene activity scores of representative genes across neoplastic cell states in snATAC-Seq demonstrating chromatin accessibility at top differentially expressed genes defined transcriptionally through the snRNA-Seq data.

**f)** Correlation of cell state proportions for each sample that has paired snRNA-Seq and snATAC-Seq data, with Pearson's correlation coefficient. Each point represents the frequency of that neoplastic cell state as a fraction of all neoplastic cells in each modality.

**Supplementary Figure 5.** Select statistically significant ligand-receptor interactions predicted by LIANA (aggregate rank/p-adjusted <0.05) from myeloid subtypes to neoplastic cell states, demonstrating differential and specific interactions through which myeloid cells can modulate tumor cell functions. Dot size represents specificity of the interaction, using the NATMI edge specificity score. Color represents expression magnitude, using the SingleCellSignalR LR score.

**Supplementary Figure 6. Validation of CODEX antibody markers.** Greyscale images of each of the 51 antibody markers on the panel, demonstrating appropriate staining morphology and subcellular localization as well as high ratio of antibody staining to background signal. Each marker is shown at two locations with varying levels of magnification to highlight tissue-level and subcellular staining patterns. Representative images are chosen from samples across the cohort.

**Supplementary Figure 7. Overview of cell type and neighborhood annotations in CODEX dataset.** Top, hematoxylin and eosin (H&E)-stained image of the exact slide profiled via CODEX image. Slides were stained after completion of CODEX imaging. Middle, cell type label masks for all segmented and annotated cells that remained in the final dataset after filtering and removal of artifacts. Bottom, cell neighborhood masks for all cells corresponding to the middle panel.

**Supplementary Figure 8. Representative CODEX images.**

- **a)** Representative example of $MPO^{+}$ myeloid cells infiltrating a region of necrosis. DAPI (blue), CD11b (green), MPO (yellow), HIF1A (red).
- **b)** Two examples demonstrating that T cells present in the data are confounded by their intravascular position (top) or clustered in areas of hemorrhage (bottom). DAPI (blue), Collagen IV (white), CD3E (magenta).
- **c)** Examples of heterogeneous distribution of immune checkpoint markers, PD-L1 and CD47 in two different CODEX samples.
- **d)** Example of spatial stratification of immune populations, highlighting microglia preference for the tumor-normal boundary, preponderance of macrophages in the tumor, and monocytic cells localized in and around vasculature. DAPI (blue), TMEM119 (green), CD68 (magenta), CD14 (orange).
- **e)** Example of peri-necrotic aggregate of MES-like-1 tumor cells with positive staining for APOE and SPP1. Top, CODEX images; bottom left, H&E image; bottom right, cell phenotype mask.

#### Supplemental Figure 1

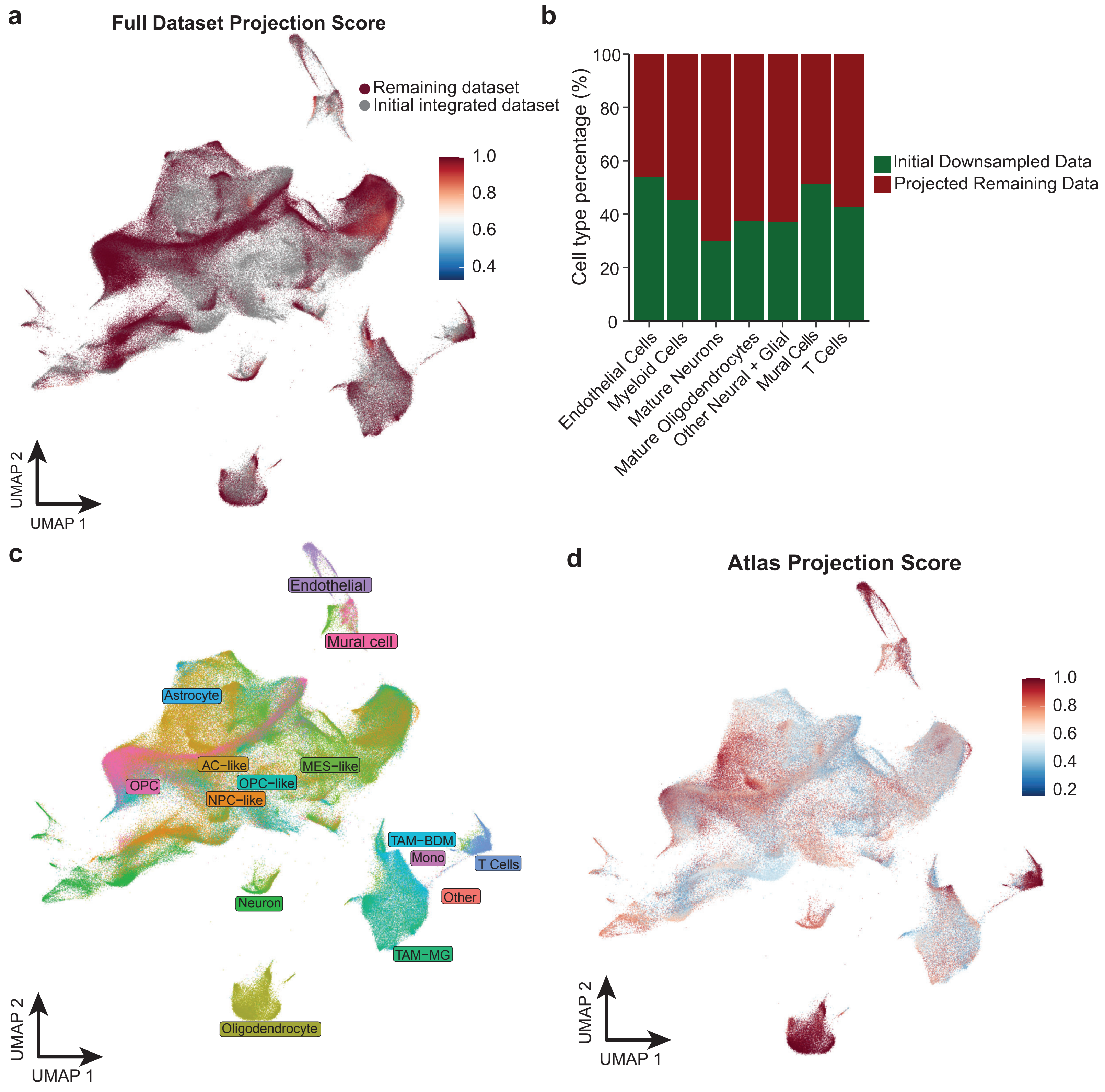

**Supplemental Figure 2**

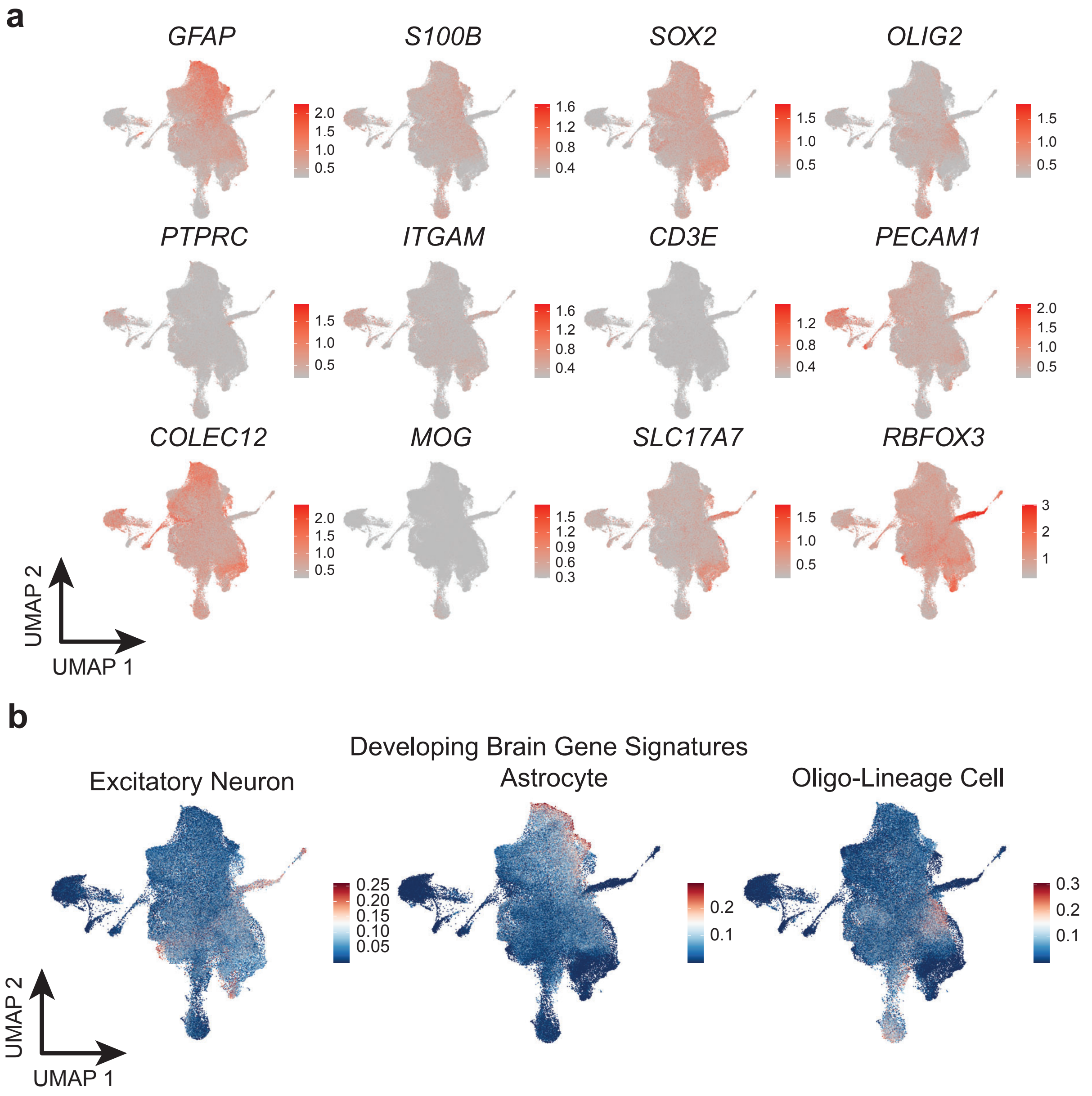

Supplemental Figure 3

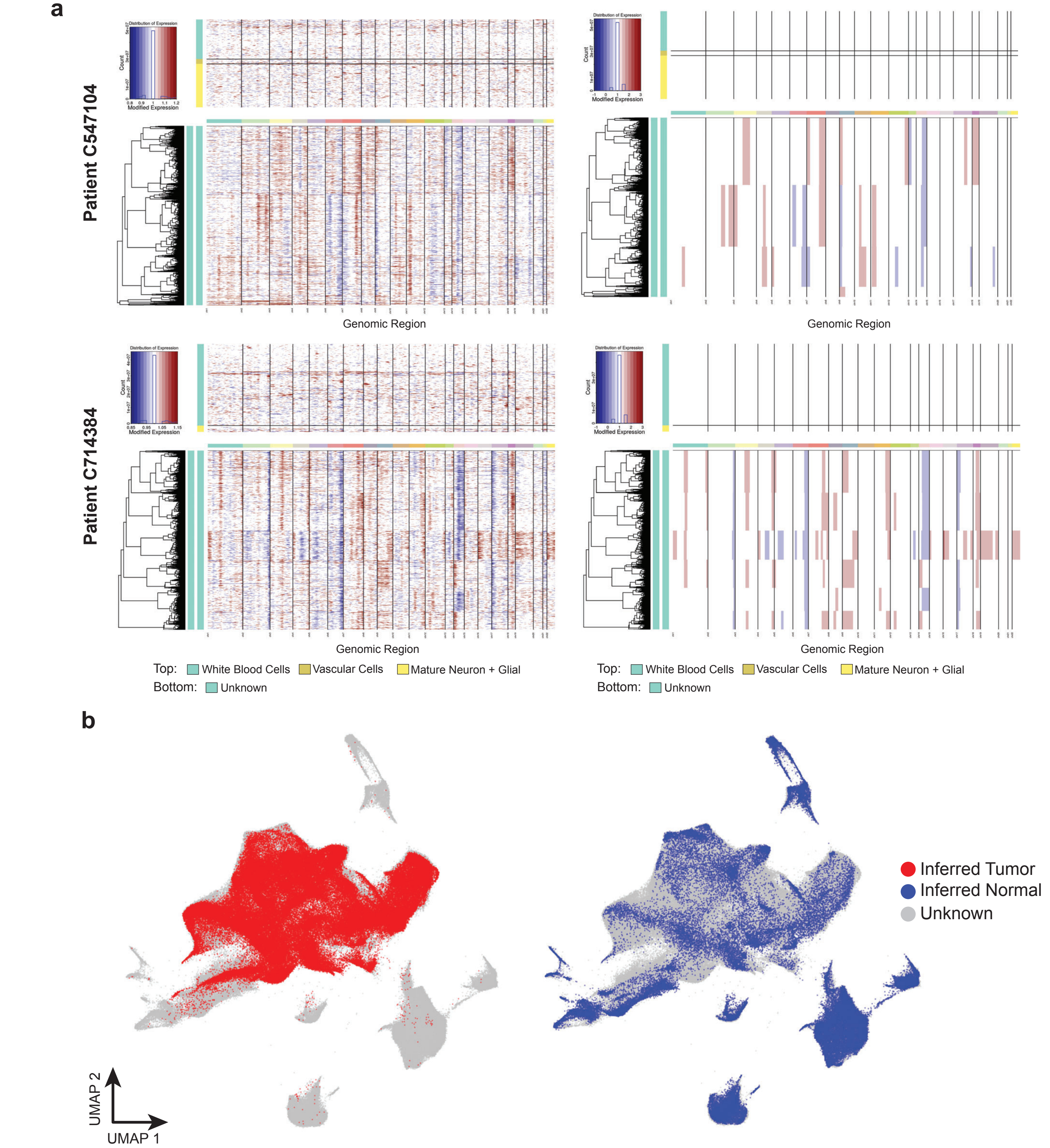

#### Supplementary Figure 4

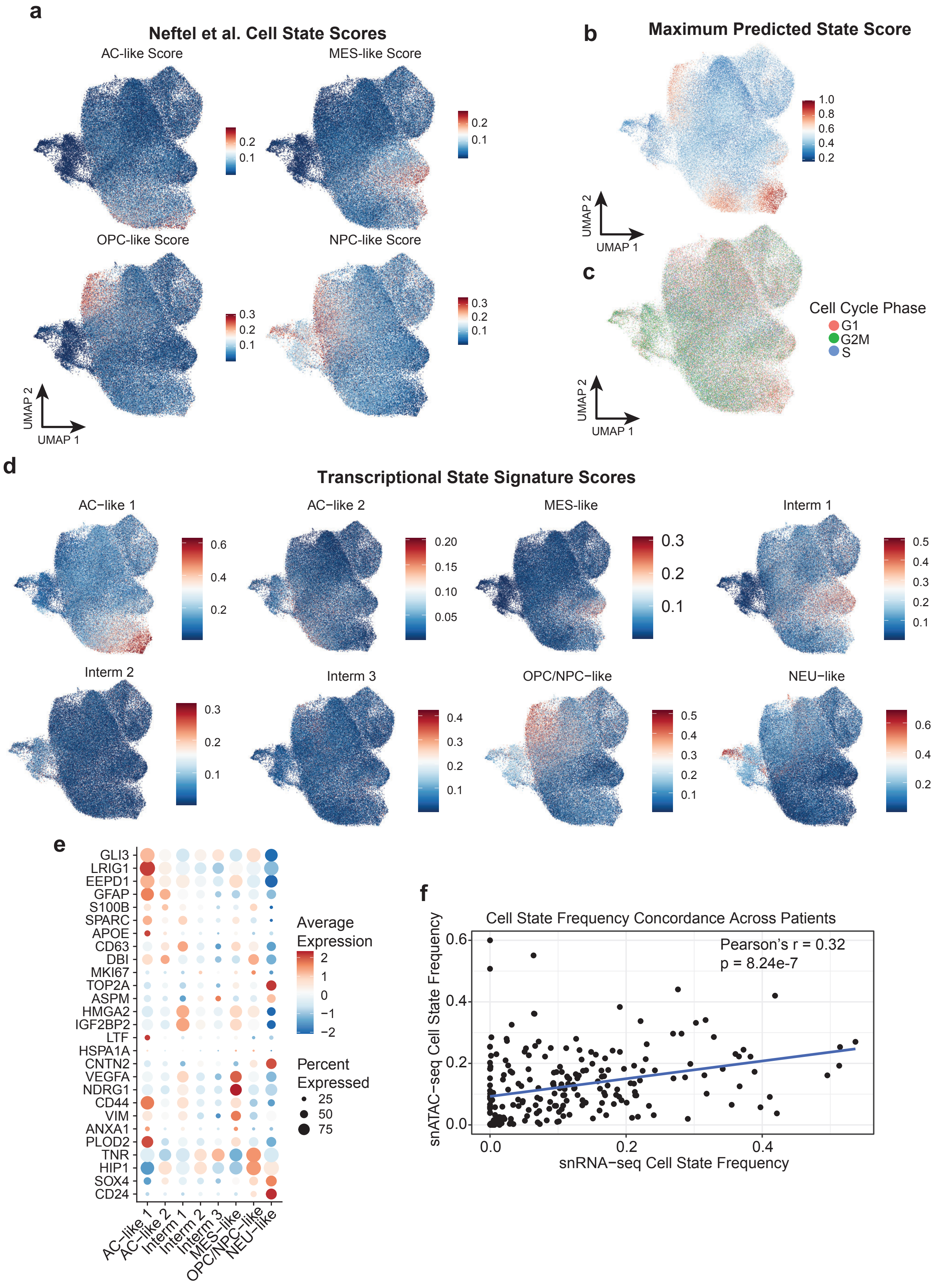

**Supplemental Figure 5**

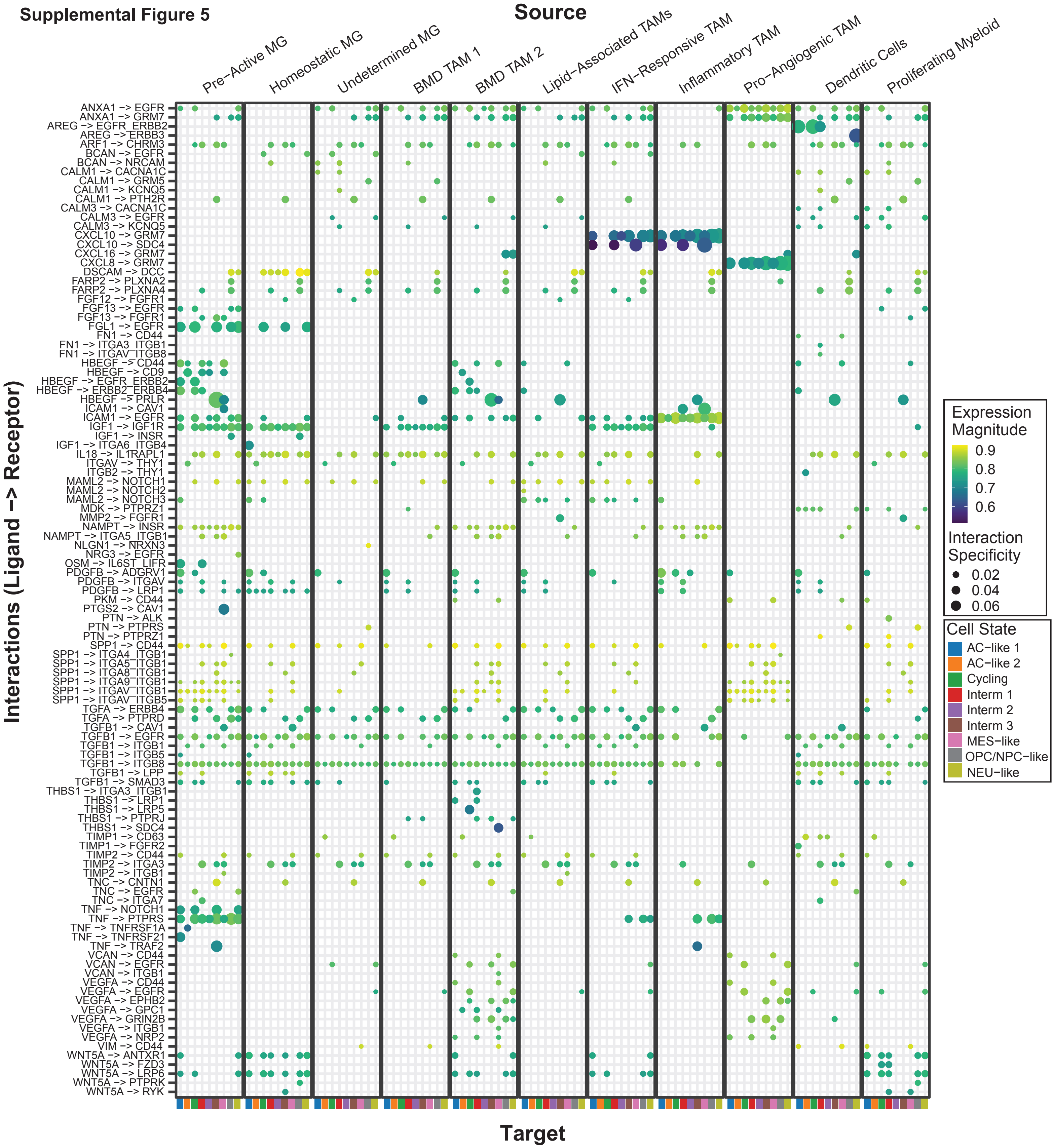

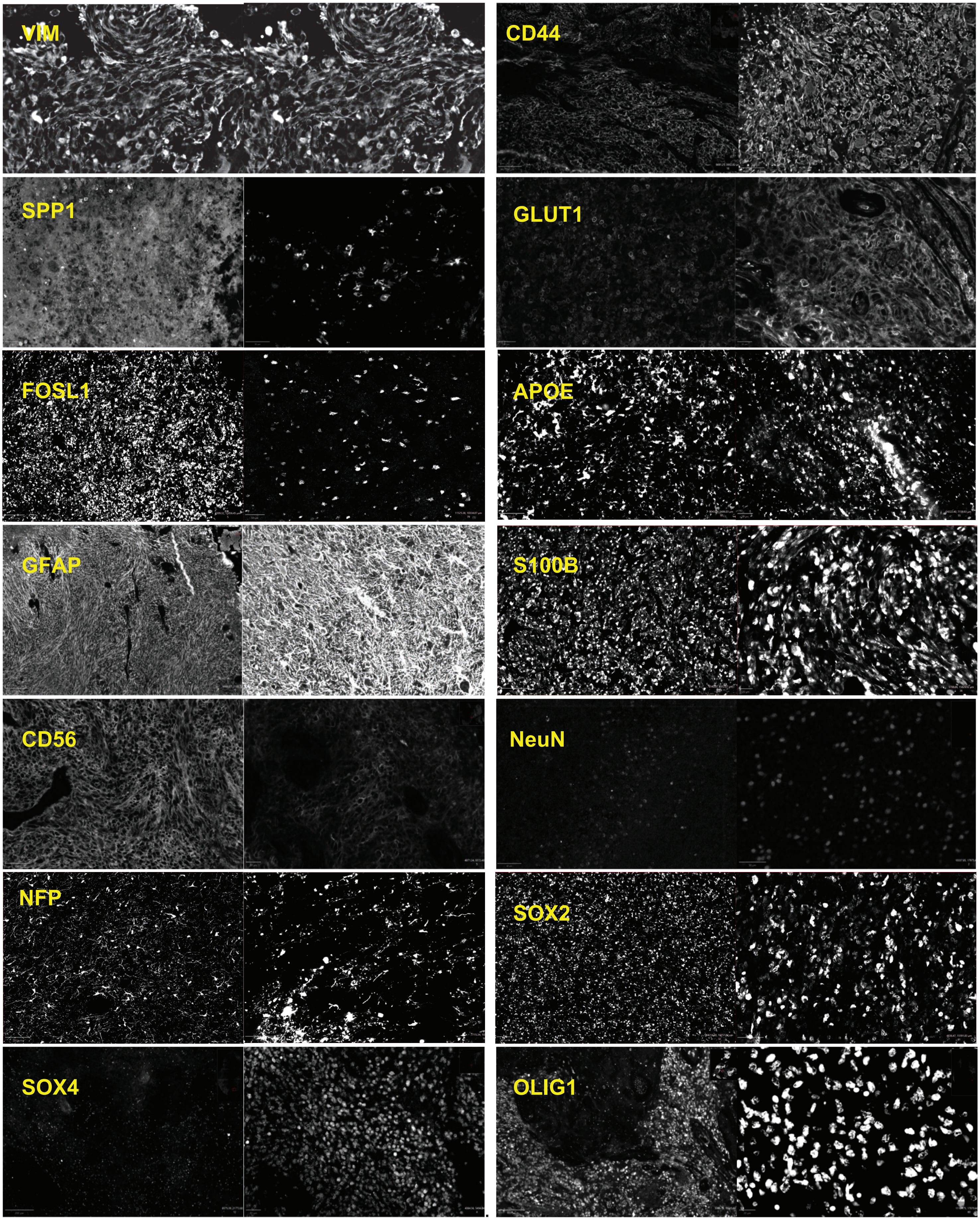
VIM
CD44
SPP1
GLUT1
FOSL1
APOE
GFAP
S100B
CD56
NeuN
NFP
SOX2
SOX4
OLIG1

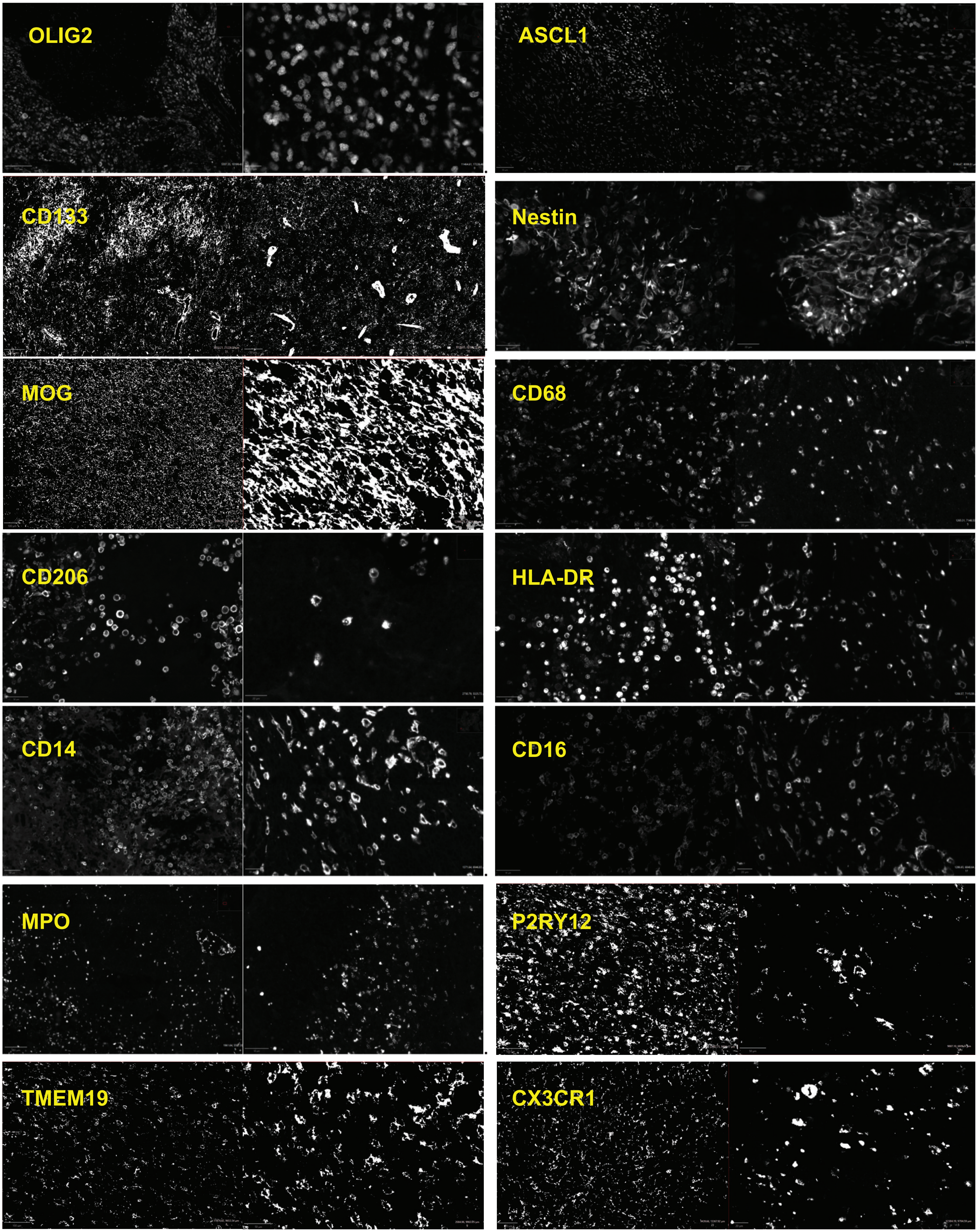
OLIG2
ASCL1
CD133
Nestin
MOG
CD68
CD206
HLA-DR
CD14
CD16
MPO
P2RY12
TMEM19
CX3CR1

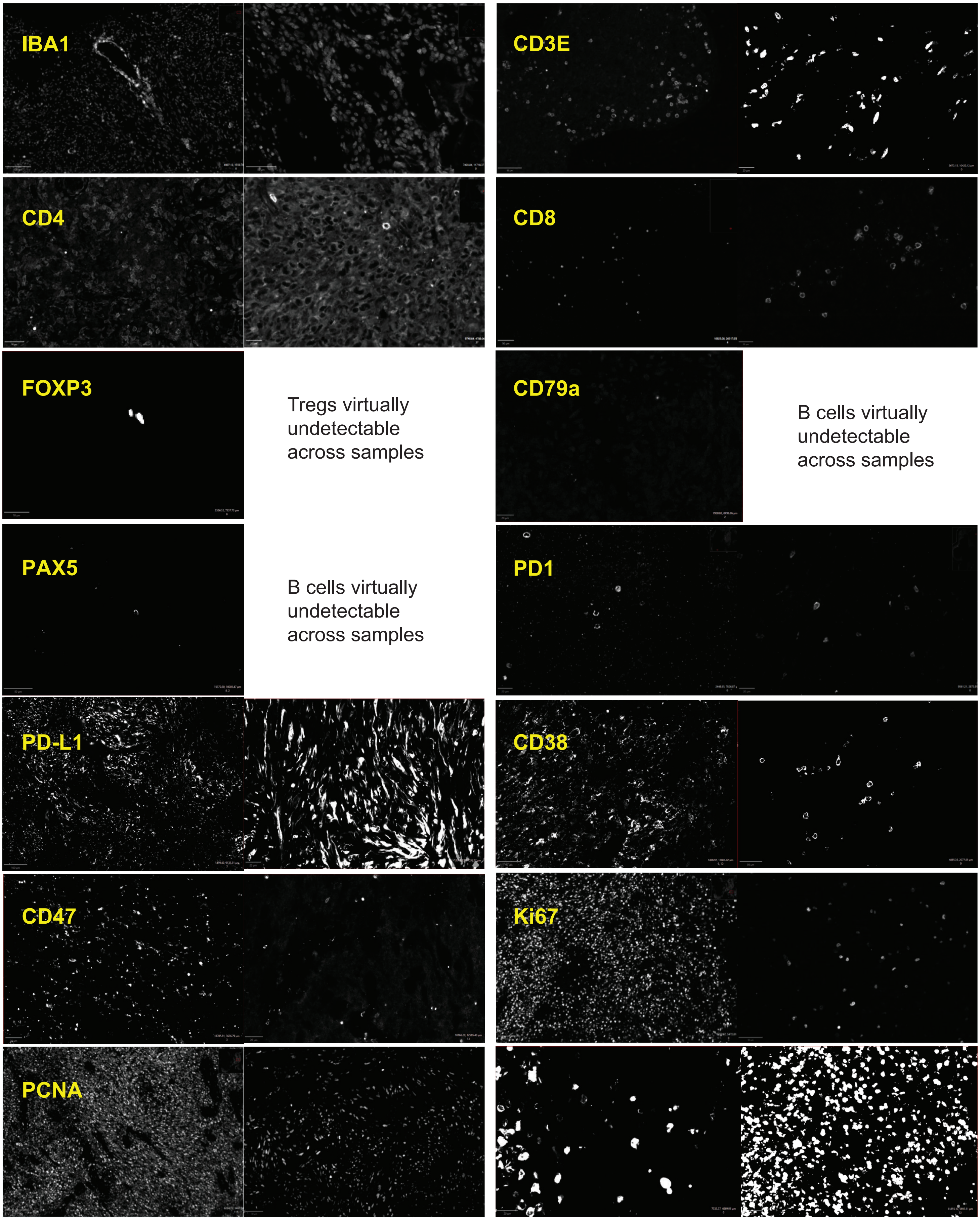

IBA1
CD3E
CD4
CD8
FOXP3
Tregs virtually undetectable across samples
CD79a
B cells virtually undetectable across samples
PAX5
B cells virtually undetectable across samples
PD1
PD-L1
CD38
CD47
Ki67
PCNA

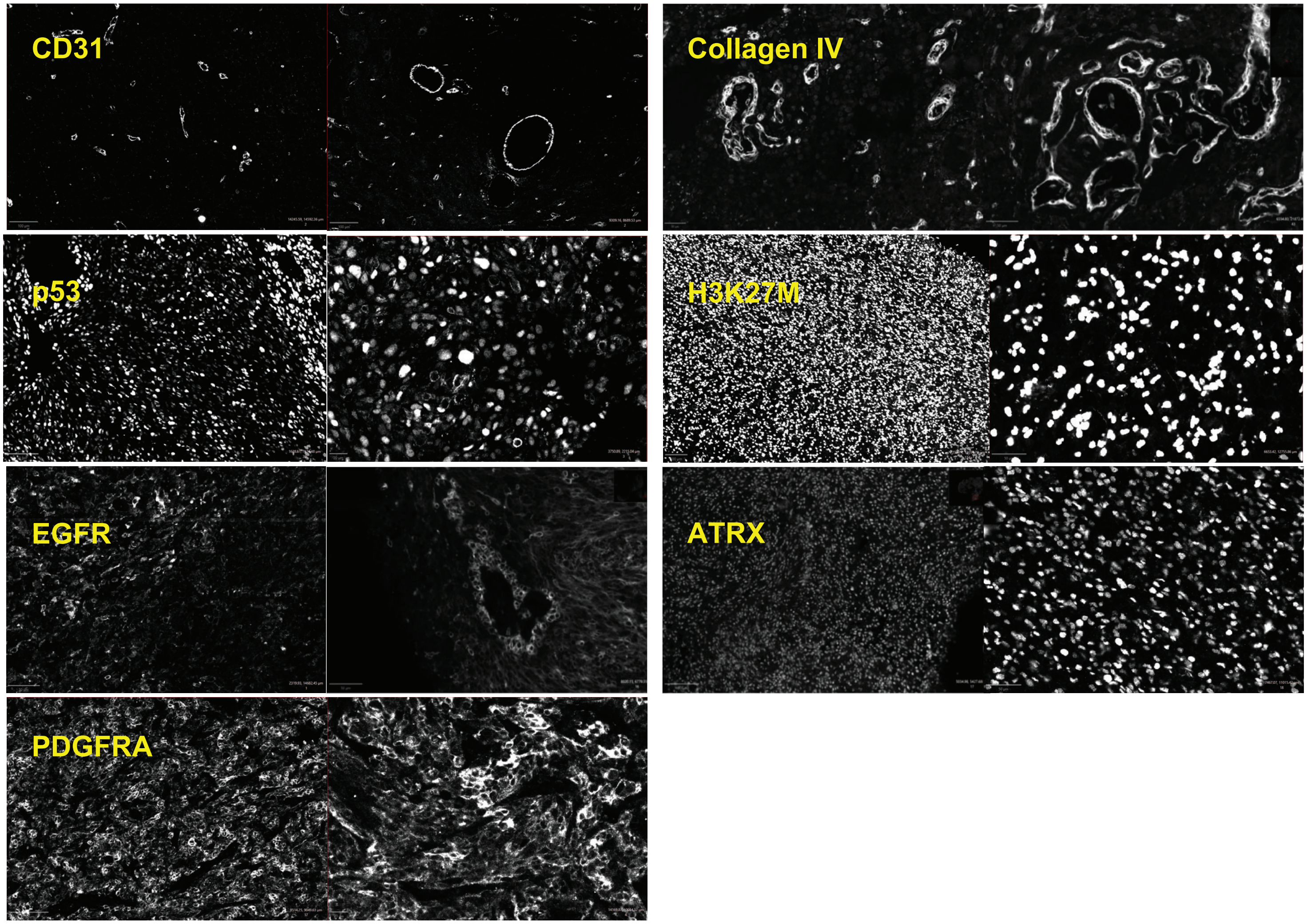
CD31
Collagen IV
p53
H3K27M
EGFR
ATRX
PDGFRA

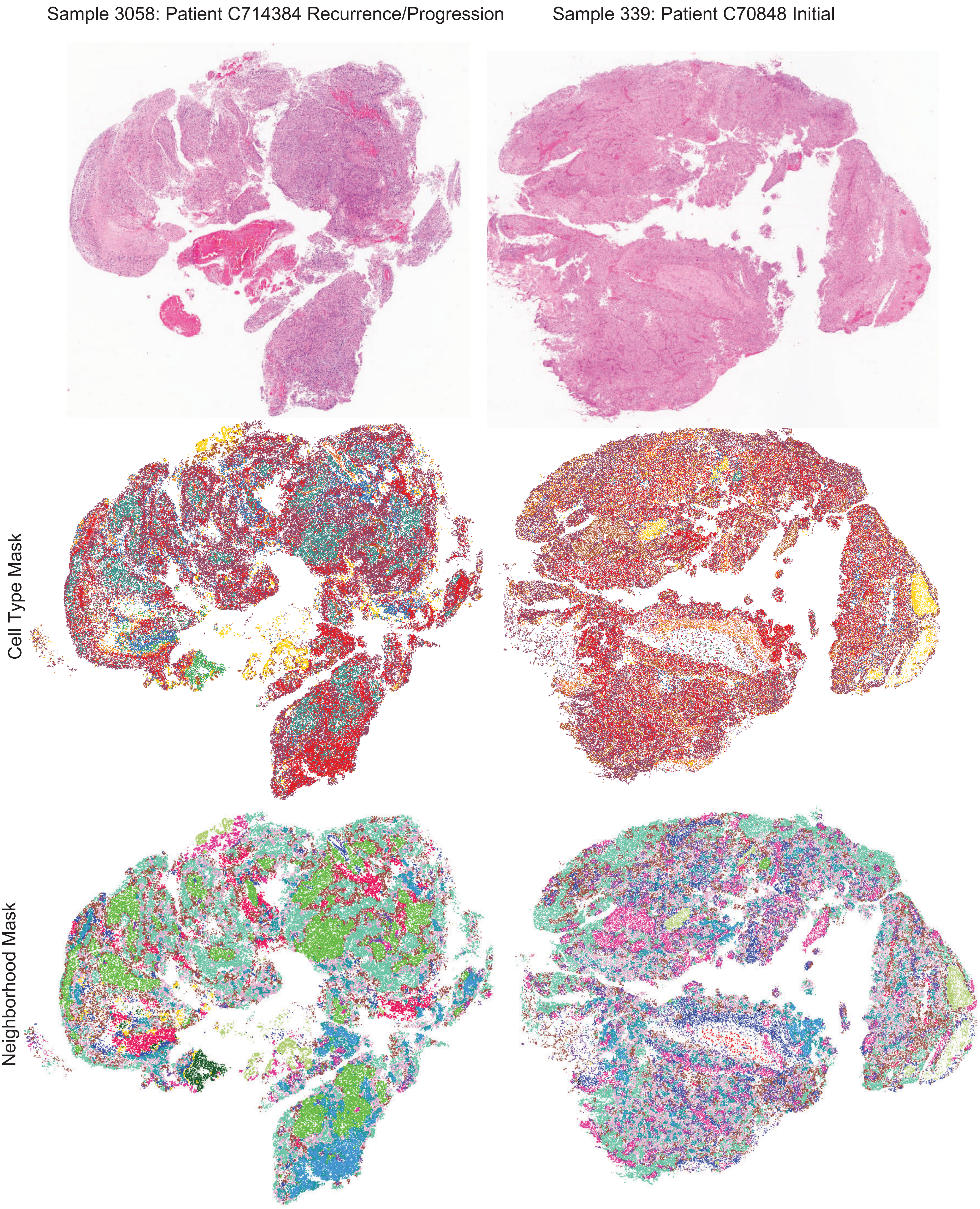

Sample 3058: Patient C714384 Recurrence/Progression
Sample 339: Patient C70848 Initial
Cell Type Mask
Neighborhood Mask

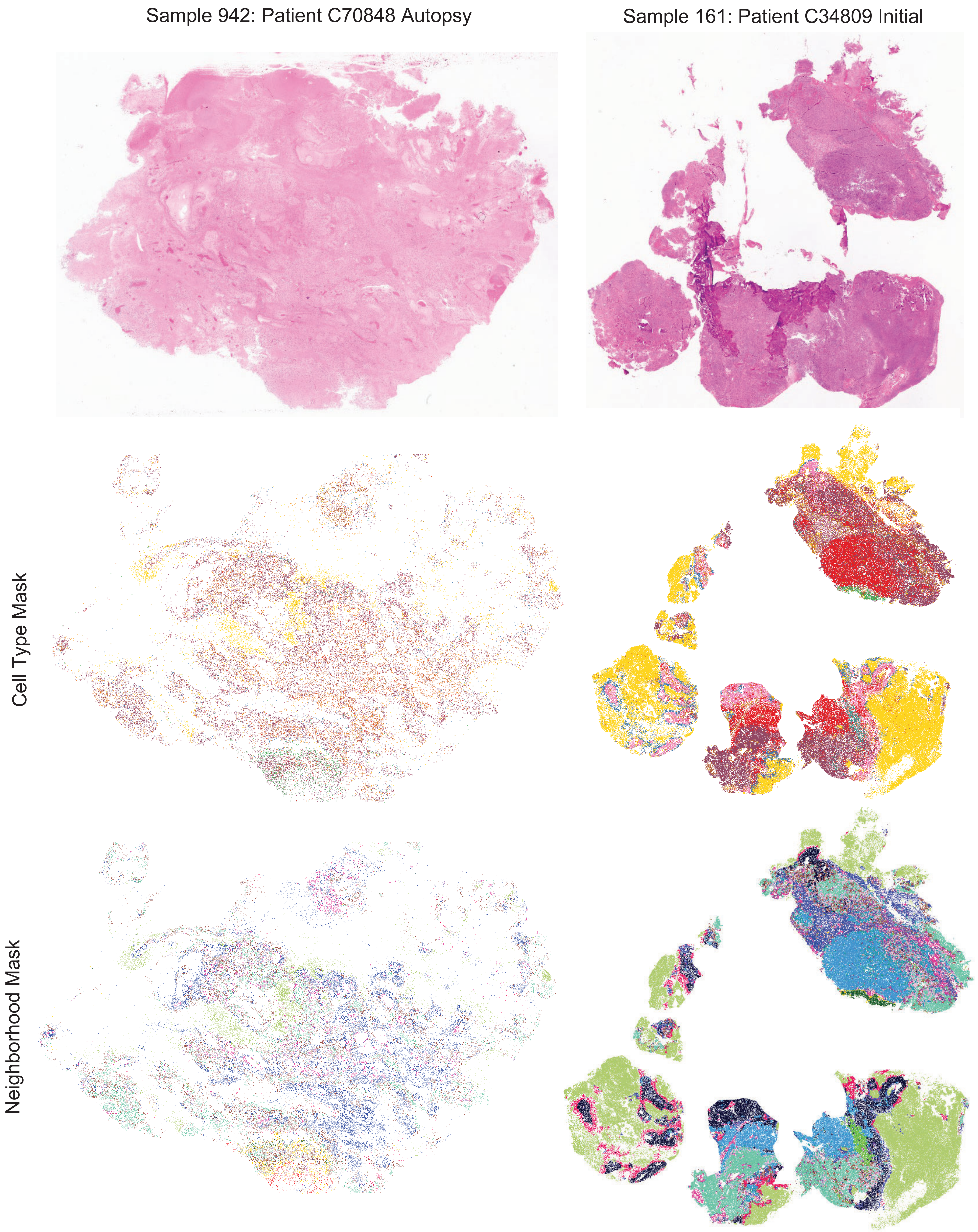
Sample 942: Patient C70848 Autopsy
Sample 161: Patient C34809 Initial
Cell Type Mask
Neighborhood Mask

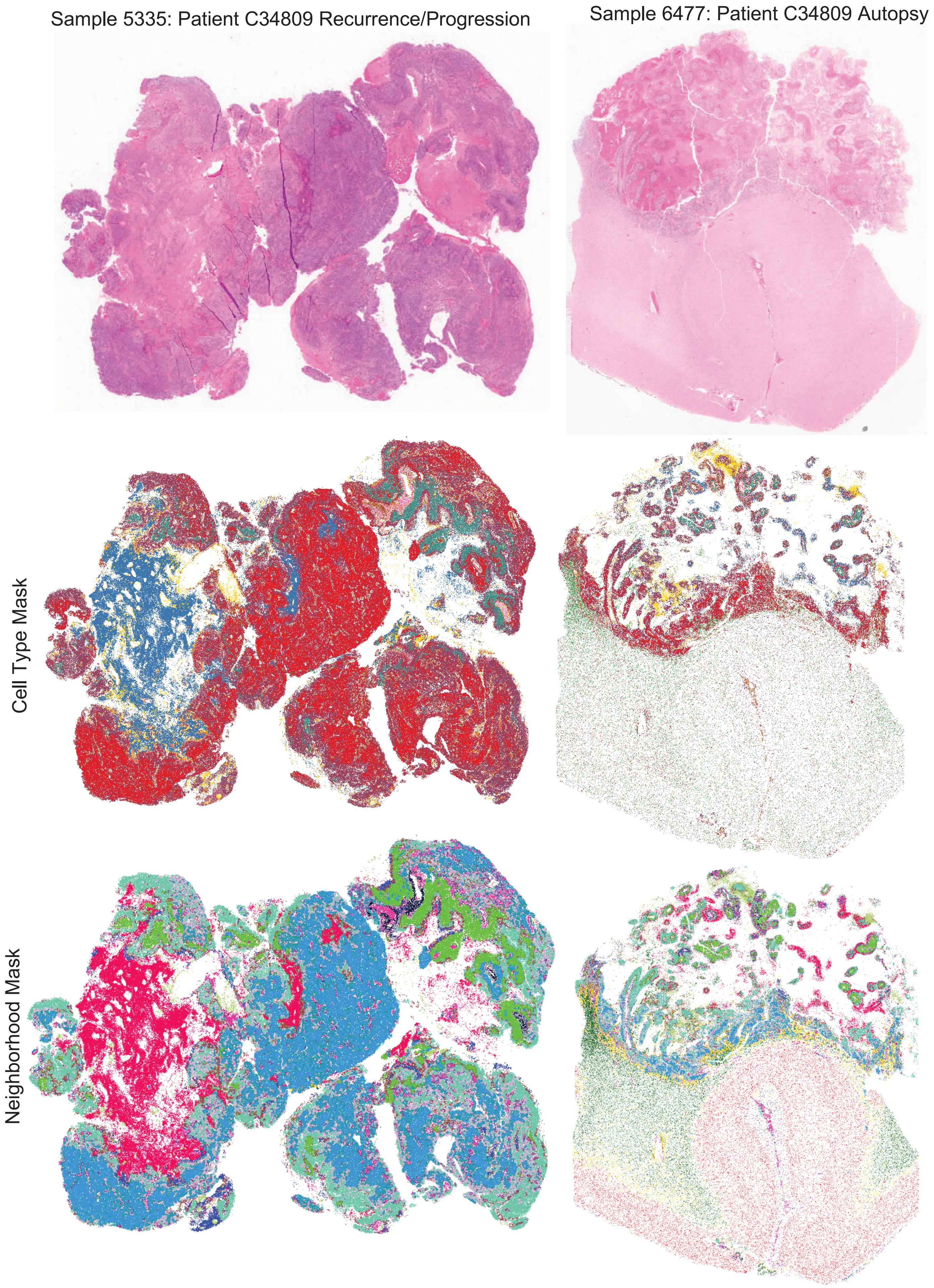
Sample 5335: Patient C34809 Recurrence/Progression
Sample 6477: Patient C34809 Autopsy
Cell Type Mask
Neighborhood Mask

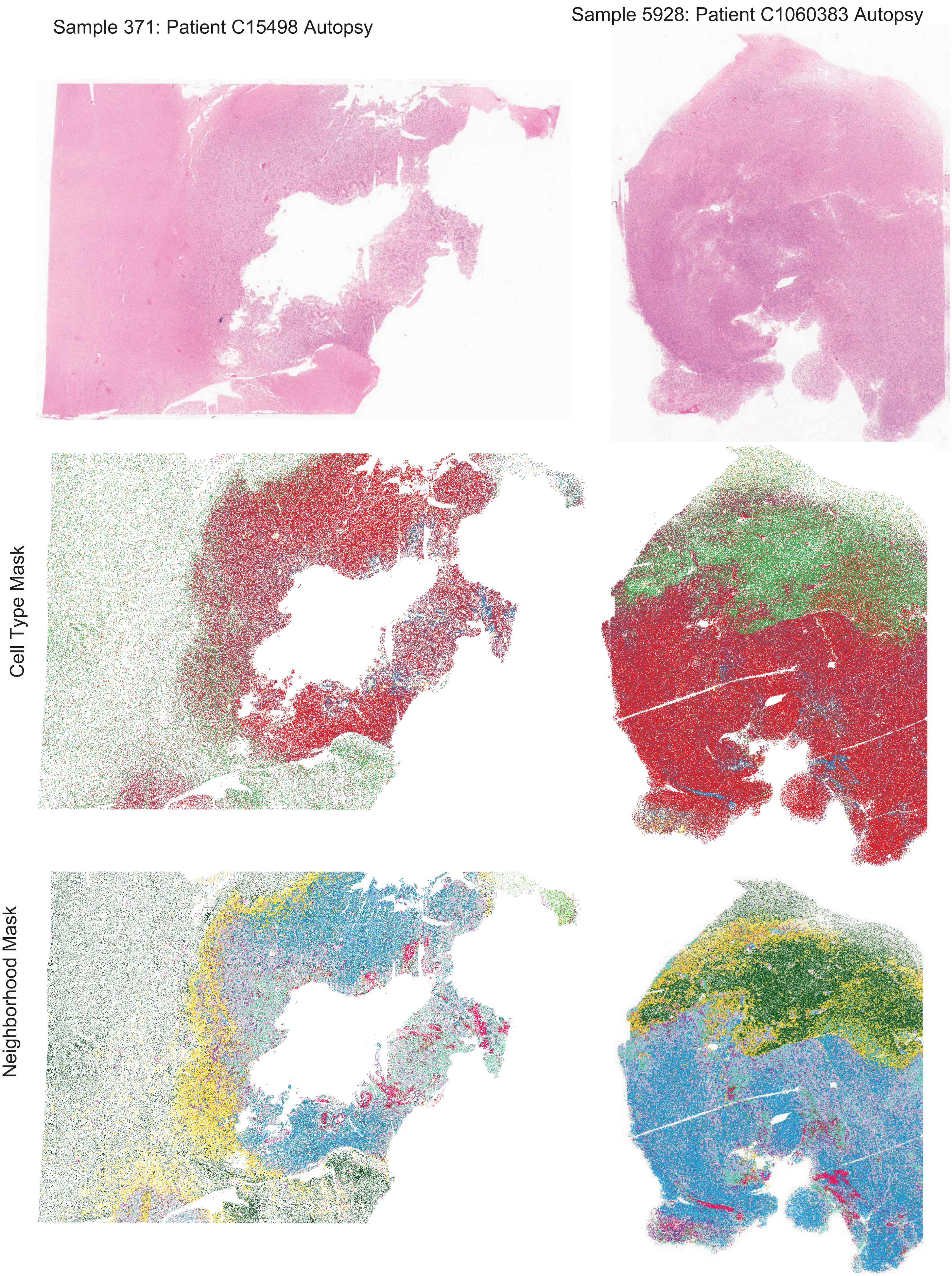
Sample 371: Patient C15498 Autopsy
Sample 5928: Patient C1060383 Autopsy
Cell Type Mask
Neighborhood Mask

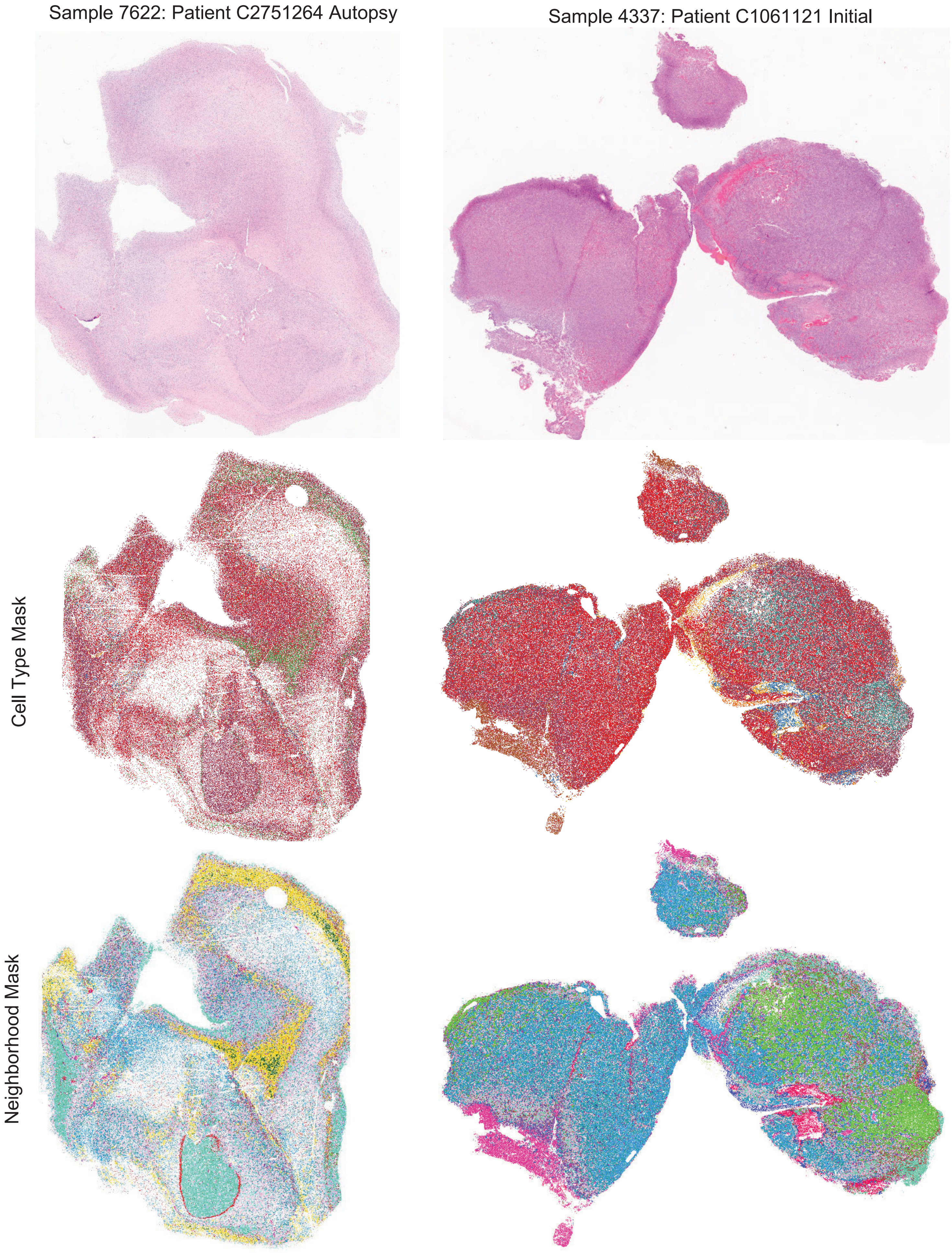
Sample 7622: Patient C2751264 Autopsy
Sample 4337: Patient C1061121 Initial
Cell Type Mask
Neighborhood Mask

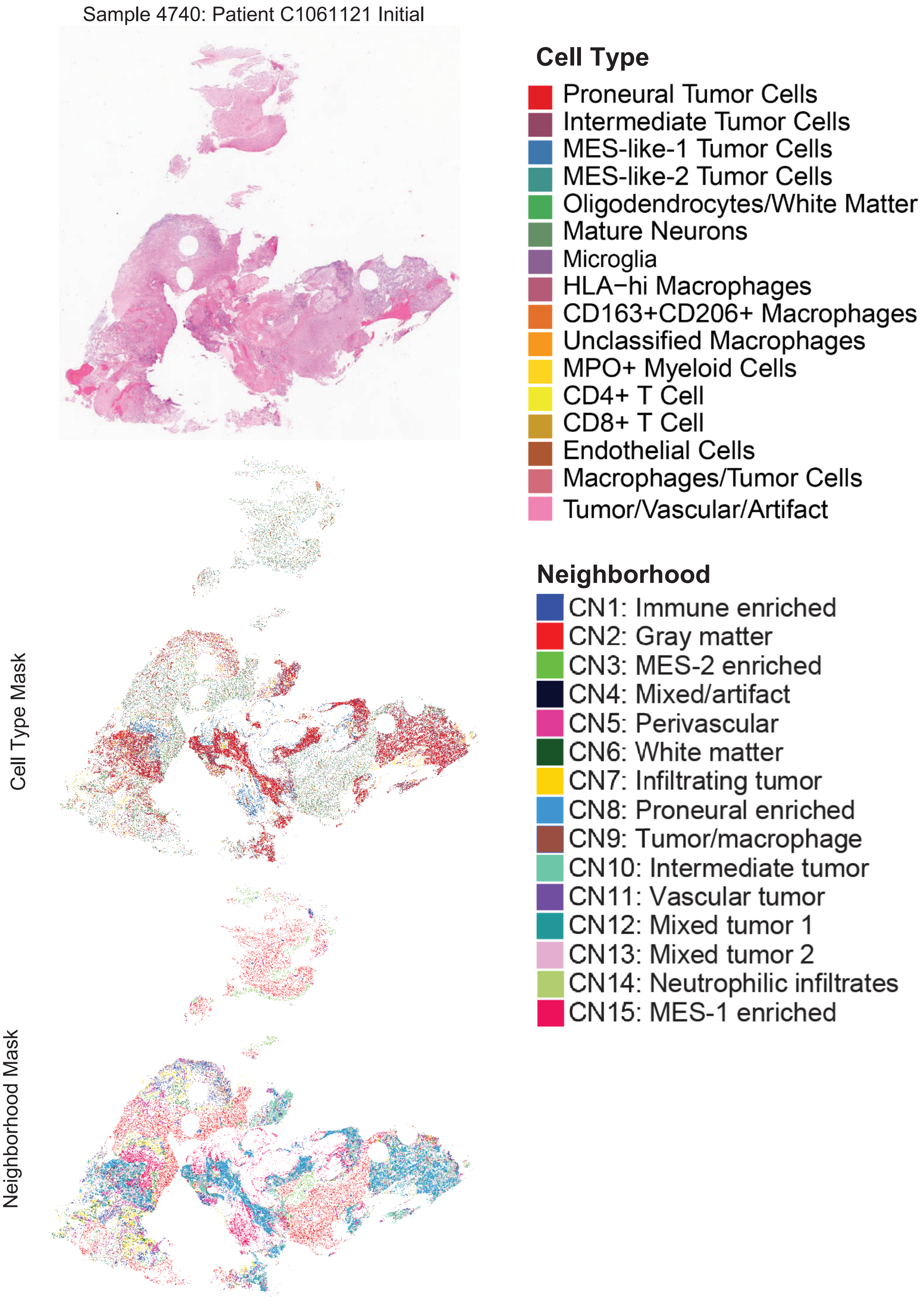
Sample 4740: Patient C1061121 Initial
Cell Type Mask
Neighborhood Mask
Cell Type
Proneural Tumor Cells
Intermediate Tumor Cells
MES-like-1 Tumor Cells
MES-like-2 Tumor Cells
Oligodendrocytes/White Matter
Mature Neurons
Microglia
HLA−hi Macrophages
CD163+CD206+ Macrophages
Unclassified Macrophages
MPO+ Myeloid Cells
CD4+ T Cell
CD8+ T Cell
Endothelial Cells
Macrophages/Tumor Cells
Tumor/Vascular/Artifact
Neighborhood
CN1: Immune enriched
CN2: Gray matter
CN3: MES-2 enriched
CN4: Mixed/artifact
CN5: Perivascular
CN6: White matter
CN7: Infiltrating tumor
CN8: Proneural enriched
CN9: Tumor/macrophage
CN10: Intermediate tumor
CN11: Vascular tumor
CN12: Mixed tumor 1
CN13: Mixed tumor 2
CN14: Neutrophilic infiltrates
CN15: MES-1 enriched

### Supplementary Figure 8

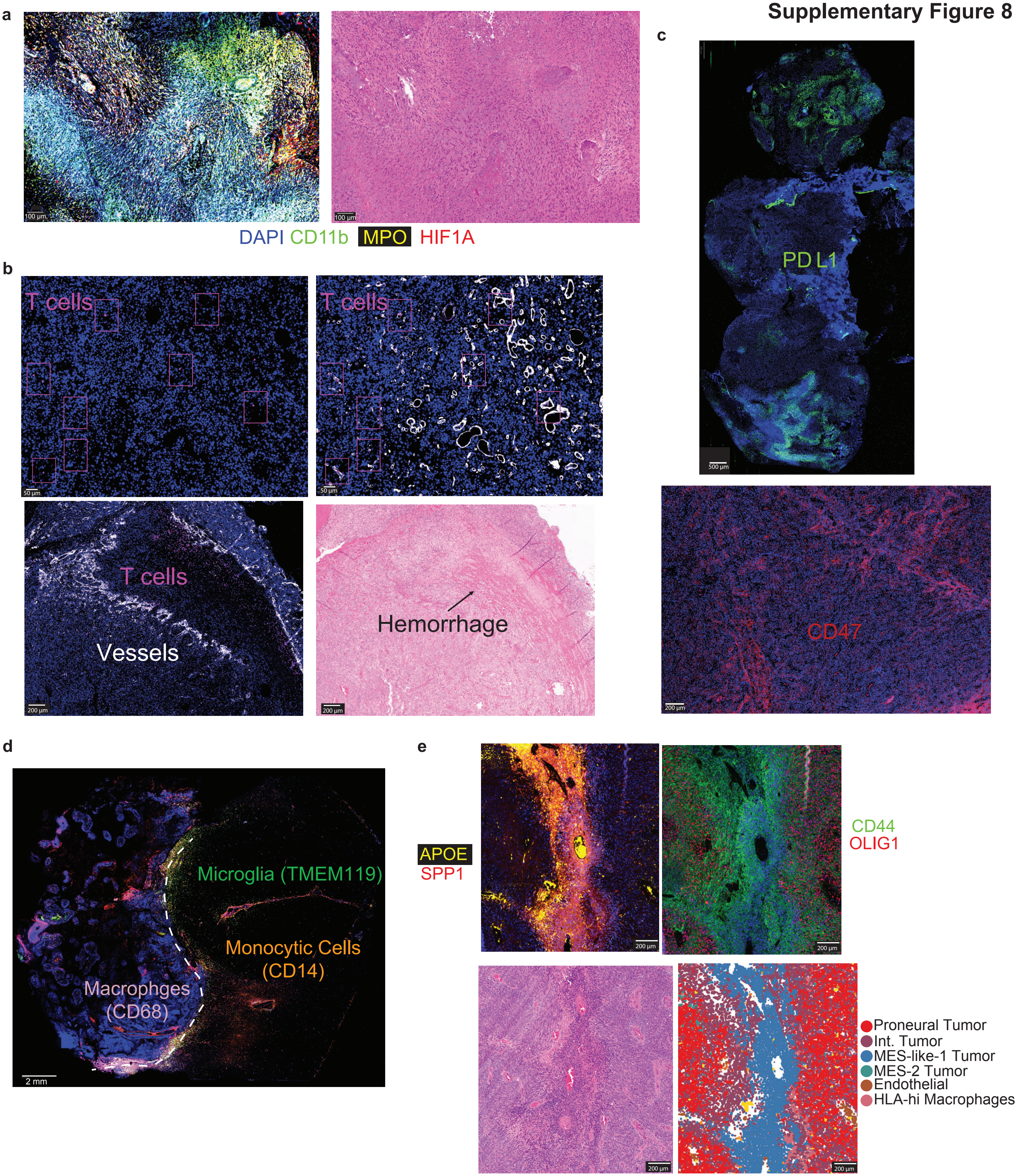
