## Supplementary Methods for "A longitudinal single-cell and spatial multiomic atlas of pediatric high-grade glioma"

**Tissue and sample preparation for single-nucleus RNA and ATAC sequencing**

Fresh frozen tumor samples were embedded in optimal cutting temperature (OCT) compound and cryosectioned into 40 um tissue scrolls. Cryosections (4-6 scrolls) were homogenized by using a 2-mL Dounce homogenizer (MilliporeSigma, D8938) with 1 mL ice-cold Nuclei Extraction Buffer (NEB, consisting of 1X PBS, 20mM Tris-HCl, 320mM sucrose, 5mM CaCl2, 3mM MgAc2, 0.1mM EDTA, and 0.1% TrionX-100). The tissue was homogenized 10 times with pestle A and 10-12 times with pestle B on ice. After ensuring that the tissue had been dissociated completely, an additional 500 µl of NEB buffer was added into each sample, then transferred from the homogenizer to a 1.7 mL Eppendorf tube with filtration using a 40 µm cell Strainer (Falcon, 352340). Filtered samples were incubated on ice for 5 mins. Nuclei were centrifuged at 500-600g for 7 mins at 4 °C and the pellet was washed twice with 1 mL of ice-cold Wash Buffer (1X PBS, 1% BSA, and 0.2U/µl RNAse inhibitor). The washed nuclei pellet was resuspended in an appropriate volume of resuspension buffer (1X PBS and 0.04% BSA), filtered through a 40 µm cell Strainer (Falcon, 352340) and nuclei were manually counted using a hemocytometer before proceeding to library preparation.

**Whole genome sequencing (WGS)**

The Quick-DNA Microprep Plus Kit (Zymo Research, D4074) was used to extract genomic DNA (gDNA) from nuclei isolated from human pediatric high-grade glioma (pHGG) tissue samples as per the manufacturer’s instructions. Sequencing libraries were generated from 1-100 ng of gDNA using the Illumina DNA Prep, (M) Tagmentation kit (Illumina, 20018705) according to manufacturer’s instructions. Bead-linked transposomes in tagmentation buffer were added to each sample at 55°C for 15 minutes in a thermal cycler to tagment gDNA and add adapter sequences. After completion of the reaction, Tagmentation Stop Buffer was added to stop the tagmentation reaction. Post-tagmentation cleanup was then performed to remove tagmented and adapter-tagged DNA from beads. This step was followed by the addition of i7 and i5 index adapters to amplify tagmented DNA. The number of PCR cycles was chosen according to directions in the user guide. The i7 and i5 indices were provided in the Nextera DNA CD Indexes kit (Illumina, 20018707). Amplified libraries were then purified using a double-sided bead purification method as outlined in the user guide. The average fragment size of purified libraries was confirmed using the Agilent 2100 Bioanalyzer with the High Sensitivity DNA kit (Agilent Technologies, 5067-4626) and library concentrations were measured using the KAPA library quantification kit (KAPA, KK4835). DNA libraries were pooled and sequenced on an Illumina NovaSeq 6000 using 150x150 bp paired-end reads.

**Whole-genome sequencing analysis**

Whole-genome sequencing (WGS) data processing and analysis was performed using published Gabriella Miller Kids First Data Resource Center (KFDRC) workflows using their corresponding public applications as implemented on the Cavatica - Seven Bridges Genomics cloud computing platform. Sequence alignment was performed using the KFDRC Alignment and GATK HaplotypeCaller application (Revision 0). Genotyping was performed using the Single Sample Genotyping Workflow application (Revision 0). Somatic mutation and structural variant calling for tumors with corresponding normal samples was performed using the KFDRC Somatic Variant Workflow application (Revision 0). Somatic mutation and structural variant calling for tumors without corresponding normal samples was performed using the KFDRC Tumor Only Pipeline (<https://github.com/kids-first/kf-tumor-workflow>, v0.1.1-beta), which incorporated a custom panel of normals (PON) to filter common technical artifacts. This PON file was built using the 10 germline WGS samples that were sequenced as part of this study and generated using a dockerized implementation of GATK (https://hub.docker.com/r/broadinstitute/gatk, version tag 4.1.1.0). The full repository of KFDRC workflows may be found here: <https://github.com/kids-first/>.

**snRNA-Seq malignant cell state analysis**

To further dissect the heterogeneity of the malignant cells, the large cluster of “other neural and glial” cells were extracted and re-integrated using Seurat v3^1^ reciprocal PCA integration. We first subset the neuroglia cells that were predicted to be malignant by CNV analysis from the downsampled Seurat object and split the object by sample ID using the *SplitObject* function. Each sample was processed individually using the standard Seurat pipeline, and then we selected 2,000 variable features with the *SelectIntegrationFeatures* function followed by running *FindIntegrationAnchors* function with *reduction* = “rpca” and *dims* = 1:30 followed by the *IntegrateData* function with *k.weight* = 50 and *dims* = 1:30. We then ran the *ScaleData*, *RunPCA and FindNeighbors* with default parameters and *FindClusters* functions with *resolution* = 0.3 and *RunUMAP* with dims = 1:30 on the *integrated* assay. To annotate each cluster of the malignant cells, we calculated the signature scores of canonical cell states defined by Neftel et al.^8^ as well as cell cycle S and G2M phase scores using the *AddModuleScore* function in Seurat.

Differentially expressed genes (DEGs) between each cluster and within each cluster across time points were calculated using the *FindAllMarkers* function with *assay* = “RNA”, *logfc.threshold* = 0.25, *min.cells.features* = 20, *max.cells.per.ident* = 500, *test.use* = “LR”, and *latent.vars* = c(“nCount_RNA”, “percent.mito”, “precent.ribo”, *only.pos* = TRUE). A pathway analysis was conducted using *gseGO* function in R package ClusterProfiler^9^ with DEGs ranked by average log2 fold change of expression and *ont* = “BP”. Each cluster was then manually annotated based on these signature scores, DEGs and pathways.

Neoplastic cells were further characterized by projection on a reference map of the developing fetal brain^10^. Briefly, the reference dataset was first reprocessed using *SCTransform* with the top 5,000 variable features using Seurat v4, and a PCA was computed. Then, the pHGG neoplastic cells were projected using the *FindTransferAnchors* function with *dims* = 1:50 followed by the *MapQuery* function with default parameters.

Lastly, the differentiation trajectory of malignant cells was inferred using CytoTRACE (v0.3.3)^11^ with default parameters using the log-normalized snRNA-Seq counts.

**Cohort-level analysis of pre- versus post-treatment gene expression changes**

*Method 1 (GLMM)*: In order to test for significant pre-treatment versus post-treatment gene expression changes assessed in snRNA-Seq data at a patient cohort level, we applied a generalized linear mixed model (GLMM) framework based on Limma-Voom (v3.46.0)^12^ and Differential expression for repeated measures (dream package, v0.4.2)^13^. Notably, dream extends upon the Limma-Voom framework by building in support for random effects modeling, which was used here to avoid pseudoreplication when patients had multiple samples from the same timepoint. Two separate GLMM analyses were performed in a pseudobulk manner based upon snRNA-Seq count matrices computed on either 1) neoplastic cells inferred by CNA or 2) myeloid cells from each snRNA-Seq sample. Prior to Limma-Voom and dream, samples with fewer than one million total reads in their corresponding pseudobulk count matrices were excluded. A counts per million (CPM) threshold of 10 was used to call genes as expressed or not expressed, and a gene was only kept if it met this threshold in at least 10 samples. Timepoint (diagnostic versus first post-treatment sample) was treated as a fixed effect and patient ID was treated as a random effect. One patient (C15498) was excluded from analysis after it was revealed during clinical data review that both timepoints were obtained post-radiation therapy due to a prior history of mixed high-grade/low-grade glioma. Genes with FDR < 0.05 by the Benjamini-Hochberg method were considered statistically significant. A gene set enrichment analysis (GSEA) was conducted to compare differentially expressed pathways over therapy. All genes from the GLM analysis were sorted by LogFC to use as input to pre-ranked GSEA using the fgsea package^14^. Pathways were sourced from the Molecular Signatures Database (MsigDB) Hallmark gene sets^15^ and the KEGG database^16^.

*Method 2 (Meta-analysis*)*:* For each patient, genes were assigned a nominal p-value using the *FindMarkers* function in Seurat v4, comparing pre- versus post-treatment expression changes in a pseudobulk manner using only neoplastic cells as inferred by the presence of CNA in both timepoints. The resulting two-sided p-values were converted to their corresponding absolute Z-score and assigned a sign based on the direction of fold-change in pre- versus post-treatment analysis for each patient. The resulting signed Z-scores for each patient were then combined by Stouffer’s Z-score method of meta-analysis, resulting in a composite signed Z-score and corresponding two-sided p-value. Genes with FDR < 0.05 by the Benjamini-Hochberg method were considered statistically significant.

**snATAC-Seq cell type annotation**

Cells in snATAC-Seq were annotated using the label transfer pipeline of Seurat v3^20^ with minor modifications. First, the gene activity score per cell was computed by summarizing fragments overlapping with the gene body or gene promoter (+/- 2kb around the TSS). The ACTIVITY assay was then added to the Seurat object through the *CreateAssayObject* function in Seurat using the gene activity score matrix, followed by log normalization through the *NormalizeData* function. Next, the *FindTransferAnchor* function was run using the downsampled snRNA-Seq Seurat Object as *reference* and the snATAC-Seq Seurat object as *query*, the highly variable genes of snRNA-Seq as *features,* ACTIVITY as the *query.assay,* and *reduction* = “cca.” Finally, the *TransferData* function was run with the annotated cell types in snRNA-Seq as *refdata*, *weigth.reduction* = “integrated_lsi”, and *dims* = 2:30. The snRNA-Seq “other neural and glial” population was further annotated as neoplastic/normal based on the inferCNV results before label transfer. Confident annotations were defined as cells having a maximum prediction score >0.9, and each cluster was assigned to the most abundant confident cell type annotation within that cluster. The predicted labels were further validated through the signature scores calculated based on signatures genes from Couturier et al.^10^ data for each type of the cells. The cells which had been assigned to non-neoplastic neuroglial cells were re-assigned as mature neurons based on the signature score (**Supplementary Figure 2b)**.

**Transcription factor motif analysis**

Transcription factor (TF) motif enrichment scores was computed using chromVAR^21^ for each cell. The cisBP core TFs “human_pwms_v2” from the chromVARmotif R package^22^ were used as the TF database. TF motifs enriched in each snATAC-Seq cell cluster were identified by performing a one-sided Wilcoxon rank-sum test comparing chromVAR deviation score between the cluster and other clusters after downsampling each comparison group to 500 cells each.

**snATAC-Seq malignant cell state analysis**

To further dissect malignant cells in snATAC-Seq data, we extracted the large cluster of neural and glial cells and re-integrated them by Signac pipeline, using the same process used to integrate the complete samples. Malignant cells were identified via label transfer as described above and were clustered using the *FindClusters* (*resolution* = 0.75). Briefly, we performed label transfer by Seurat as described above, using the cell state annotation of malignant cells in snRNA-Seq data as the reference. The clusters with > 5% of cells having a maximum cell state prediction score >0.6 were assigned to the most abundant cell state within the cluster. Remaining clusters were annotated manually based on a combination of signature scores derived by the DEGs from each snRNA-Seq cell state using the ATAC gene activity assay and chromVAR TF motif analysis (**Supplementary Figure 1**).

**Myeloid subtype analysis**

Tumor-associated myeloid populations were investigated by subsetting the macrophage/microglia population from the full snRNA-Seq dataset and splitting this object by patient ID using the *SplitObject* function. Each sample was processed individually using SCTransform regressing out nCount_RNA, mitochondrial read percentage, and ribosomal read percentage. We then selected 3,000 variable features with the *SelectIntegrationFeatures* function followed by running *FindIntegrationAnchors* function with *reduction* = “rpca”, *dims* = 1:30, and *k.anchor =* 20 followed by the *IntegrateData* function with *dims* = 1:30. We then ran the *RunPCA, RunUMAP, FindNeighbors,* and *FindClusters* functions on the *integrated* assay with *dims* = 1:30 and *resolution =* 0.1. We then projected this data onto the GBM atlas^6^ as described above and observed two clusters that represented misclustering of neuroglial cells, confirmed by specific expression of astrocyte and oligodendrocyte identity genes. These clusters were removed, and the remaining cells were reprocessed and reintegrated with the same procedure as above and reclustered with *resolution* = 0.6. To annotate each cluster of the myeloid cells, we first calculated the signature scores of canonical microglia and bone marrow-derived macrophages defined by Müller et al^25^ as well as cell cycle S and G2M phase scores using the *AddModuleScore* function in Seurat. Differentially expressed genes (DEGs) for each cluster were calculated using the *FindAllMarkers* function with *test.use = “*wilcox”*, min.pct =* 0.10*, min.diff.pct =* 0.10*, logfc.threshold =* 0.10, and clusters were annotated manually based on these gene signatures and DEGs.

**SCENIC transcriptional regulatory network (TRN) analysis**

The Single-Cell Regulatory Network Inference and Clustering (SCENIC) algorithm^26^ was used to identify transcription factors regulating myeloid cell states. It was run using the pySCENIC implementation as previously described^27^ on the scRNA-seq data, after conversion of the Seurat object to loom format. The GRNBoost2 algorithm was used for GRN inference. To predict transcription factor regulons, we used the human v9 cisTarget motif collection and the hg38_refseq-r80 databases with the 500bpUp100Dw and TSS+/-10kb search spaces. All relevant databases were obtained from: https://resources.aertslab.org/cistarget/. The pySCENIC implementation of AUCell was used to score the activity of regulons for each cell. The *SCopeLoomR* package (https://github.com/aertslab/SCopeLoomR) was then used to extract the regulons and AUCell matrix from the resulting loom file, and the final AUC matrix was added to the initial Seurat object for visualization and analysis. Differential regulons were identified using the *FindAllMarkers* function using the Wilcoxon rank sum test with “*logfc.threshold* = 0.01.”

**CODEX antibody conjugation**

Akoya antibodies were purchased pre-conjugated to their respective CODEX Barcode (**Supplemental Table S2**). All other antibodies were custom conjugated to their respective CODEX barcode (**Supplemental Table S2**) according to Akoya’s PhenoCycler-Fusion user guide using the antibody conjugation kit (Akoya, 7000009) following manufacturer’s protocol. Briefly, 50 μg of carrier-free antibodies were concentrated by centrifugation in 50kDa MWCO filters (EMD Millipore, UFC505096) and incubated in the antibody disulfide reduction master mix for 30 minutes. Buffer exchange of the antibodies was then performed to stop the reduction reaction by centrifugation, addition of conjugation solution, and an additional centrifugation. Respective CODEX barcodes resuspended in nuclease free water and conjugation solution were added to the concentrated antibody and incubated for 2 hours at room temperature. Conjugated antibodies were purified by 3 buffer exchanges with purification solution. Antibody storage buffer (100 μl) was added to collect the concentrated purified antibodies. Successful conjugation was confirmed using the Agilent 2100 Bioanalyzer with the Agilent Protein High Sensitivity kit (Agilent Technologies, 5067-1580), following the manufacturer’s instructions.

**CODEX data integration and annotation**

Individual Seurat objects were merged, which was then normalized by CLR normalization in Seurat with *margin* = 2. The Seurat object was again split by sample, and downsampled using the *SketchData* function with *method =* “LeverageScore” with 50,000 cells per sample. The data was scaled, and a principal component analysis was computed. The samples were integrated using the *IntegrateLayers* function with RPCA integration using the first 15 dimensions. The integration features were selected based on manual inspection of each marker across samples to identify markers with consistent staining across the majority of the samples. Cell cycle genes were excluded, and DAPI was included to help identify autofluorescence artifact. The markers selected were: DAPI, CD31, CD44, IBA1, NFP, CD4, ATRX, APOE, CD56, CD163, GFAP, CD11b, S100B, CD206, OLIG1, CD133, SOX2, Vimentin, MPO, HLA-DR, P2RY12, CD8, NeuN, Collagen IV, CD14, CX3CR1, MOG, SPP1, CD3e, CD68, Nestin, CD16, TMEM119, OLIG2, GLUT1.

Cells were clustered using the *FindNeighbors* function on the integrated assay with the top 15 principal components followed by *FindClusters*, and a UMAP was computed for visualization. Based on marker expression, clusters were assigned to an initial annotation of endothelial cells (CD31, Collagen IV), neuroglial cells including tumor cells (SOX2, OLIG2, Vimentin, GFAP, MOG, NFP), myeloid cells (CD68, CD11b, CD14, P2RY12, TMEM119) , or T cells (CD3, CD4, CD8). The remaining cells were projected onto this sketched integration using *ProjectIntegration* and *ProjectData* with the same features and dimensions as above. Subsequently, the neuroglia, myeloid, and T cell clusters were subjected to a second round of annotation. They were first subset, downsampled through sketching, scaled, integrated, clustered, and the remaining cells projected using the same parameters as above. Cell type masks were generated for each category and manually evaluated and labeled in Napari v0.4.18. Clusters which were found to represent apparent artifacts including necrotic cells, autofluorescence artifact, tissue folding, paraffin wax drops, and red blood cells were excluded. Blank channels were used to help identify artifacts. CD4^+^ and CD8^+^ T cells could not be distinguished via clustering, so manual thresholding based on CD8 was used to distinguish them. All other clusters were labelled based on their marker expression and general morphology. After filtering, cells were subjected to one more round of sketching, integration, and projection in order to build the final UMAP for visualization.

**Cell proximity analysis**

We measured the proximity of each cell type to every other cell type using the median distance through the spatstat R package^39^. We used the *ppp* and *psp* functions to create point and line segment patterns and quantified spatial proximity using a table of cell type annotations and locations with the *distfun* function. We randomly downsampled the source cell type populations to the larger value between 0.25*number of cells or 50,000 cells and restricted the observation window to a rectangular area bounded by the target cells with the maximum x and y coordinates. The source cell type was skipped if there were fewer than 10 of that cell type in a given sample. We conducted a one-sided permutation test without replacement to determine if the source cell type is significantly close to each target cell type in each sample. We permuted the labels 100 times and calculated their median distance across the 100 permutations. The proportion of distances smaller than that of the cell type is the p-value, which was defined as significant if p<0.05.

**Prioritization of drug targets for *in vitro* drug screening**

To identify targetable genes for pharmacological manipulation of tumor cell-intrinsic functions, we developed a framework that incorporates transcriptional analysis with drug databases. Drug/target data from two independent drug targets databases (TTD^40^, DrugIDB^41^), as well as a third database (OpenTargets^42^) that focuses on next-generation targets were overlapped with differentially gene expression data from our linear mixed model analysis. Targetable gene products were given a score of +1 if a resulting hit occurred from any of the three databases. To search for drugs that could modify therapeutically induced transcriptional profiles, we inputted top time point-specific DEGs (n=93 upregulated, 74 downregulated) into the LINCS1000^43^ database (https://clue.io/query#l1000) under default parameters. This method predicts perturbations that alter the collectively inputted transcriptional profile. Perturbation results across cell lines were filtered for compound-based perturbations with defined targets, false discovery rate (FDR) <0.25, and effect size (magnitude of normalized connectivity score >0.6). Both initial time point, and post-therapy-specific effects were considered on the basis that upregulated transcriptional profiles may simultaneously include positive effects of therapy in addition to the development of resistance mechanisms and requires experimental study. Thus, any gene targeted through the filtered perturbations was given a score of +1. Next, to predict glioma-specific genes that could mediate cytotoxicity when targeted, we downloaded cell line dependency data from the Cancer Dependency Map (DepMap)^44^ portal and searched for genes which showed increased dependency in glioma cell lines (n=83) compared to non-glioma cell lines (n=1070). Genes with negative dependency scores in glioma cell lines (mean dependency score < -0.1) and dependency mean fold-change >1.5 were assigned +1 to the overall score.

We further prioritized genes based on two methods of differential expression analysis, prioritizing genes that were upregulated post-therapy in order to identify additional single gene dependencies that could be targeted. First, using the linear model analysis, upregulated DEGs were assigned a score as follows accounting for degree of upregulation and statistical significance: logFC >0 and p-adjusted <0.05: +0.5, logFC >0.5 and p-adjusted <0.05: +1, logFC >0.5 and p-adjusted <0.001: +1.5, logFC >1 and p-adjusted <0.05: +1.5 (only the highest score added). Second, a patient-wise meta-analysis approach was used orthogonally, requiring a z-score >0 and then assigning a score based on the number of patients in which the gene was upregulated post-therapy: 8-9 patients: +0.5, 10-11 patients: +1, >11 patients,+1.5 (only the highest score added). Lastly, given the significance of the cellular interactions in maintaining glioma proliferation, we added a score of +1 to any genes that were predicted to be receptors in any significant ligand-receptor interactions (aggregate rank, p <0.05). Genes were then filtered to include only genes that were known drug targets and were differentially upregulated by either the linear model or meta-analysis approach, resulting in a list of 483 genes with scores ranging from 1.5 to 5. Genes with scores of 3 or greater (83 genes) were manually interrogated to identify target for *in vitro* study.
